# Discovery of Host-Directed Small Molecules with Broad Anti-Leishmanial Efficacy

**DOI:** 10.1101/2025.11.04.686469

**Authors:** Elizabeth G. Gurysh, M. Shamim Hasan Zahid, Monica M. Johnson, Antonio Landavazo, Ojas A Namjoshi, Joseph W Wilson, Devika M. Varma, Ryan N. Woodring, Aaron T. Hendricksen, Joseph F. Vath, Baiyi Quan, Erica N. Pino, Michael C. Fitzgerald, Eric M. Bachelder, Bruce E. Blough, Kristy M. Ainslie

**Affiliations:** Division of Pharmacoengineering and Molecular Pharmaceutics, Eshelman School of Pharmacy, University of North Carolina, Chapel Hill, NC; Center for Drug Discovery, RTI International, Research Triangle Park, Durham, NC; Department of Chemistry, Duke University, Durham, NC; Joint Department of Biomedical Engineering, UNC School of Medicine, University of North Carolina, Chapel Hill, NC; Department of Microbiology and Immunology, UNC School of Medicine, University of North Carolina, Chapel Hill, NC

## Abstract

Leishmaniasis, a neglected tropical disease affecting nearly 10% of the global population, suffers from limited therapeutic options and rising drug resistance. To address this, we developed 343 analogs of AR-12, a compound that has previously illustrated host-directed anti-leishmanial effects. Primary screening using a luminescence-based assay revealed 66 analogs with greater selectivity than the parent compound, AR-12. Sixteen promising candidates, selected for high potency (IC₅₀ < 1 µM) or high selectivity (>15), underwent secondary screening via Giemsa staining. Four lead compounds (53, 134, 197, and 354) demonstrated therapeutic indices greater than 40. Tertiary assays confirmed their broad *in vitro* efficacy against both *Leishmania donovani* and *L. mexicana*. Notably, 197 exhibited potent host-directed activity and proteomic analysis identified lysozyme as a mechanistic target, implicating it in the host-mediated clearance of intracellular parasites. These findings highlight the dual host– and pathogen-directed mechanisms of these compounds and support their potential as the basis for new therapeutic strategies. Further optimization and clinical exploration of these leads are warranted to meet the urgent need for effective leishmaniasis treatments.

**AUTHOR SUMMARY:** Leishmaniasis is a parasitic infectious disease with limited therapeutic options and rising drug resistance. Drugs that act directly against *Leishmania* can further drive drug resistance. Host-directed therapies work to enable the host responses to enhance pathogen clearance and reduce disease progression. Host-directed therapies both limit the emergence of new drug resistance and combat drug resistant infections. In this work we screened a library of 343 AR-12 analogs for host-directed anti-leishmanial activity. This work identified 4 hit compounds. We then did proteomic analysis to understand the mechanism of action of the primary lead compound, identifying the protein lysozyme as having a role in host-directed activity against *Leishmania* infection.

## INTRODUCTION

Leishmaniasis, an infection caused by the parasites of the *Leishmania* genus, is classified by the WHO as a neglected tropical disease. Nearly 10% of the world’s population is at risk of acquiring one form of leishmaniasis. Worldwide it is estimated that there are 12 million active cases of leishmaniasis, with approximately 1 million new cases occurring each year. Among parasitic infections, this disease is responsible for the highest number of DALYs (Disability adjusted life years; a measure of health burden) after malaria. Visceral leishmaniasis (VL) is the most clinically serious form of leishmaniasis and is fatal without chemotherapeutic intervention. For the last 70 years, the most common treatment of VL is the systemic injection of sodium stibogluconate (SSG) or meglumine antimoniate formulations (antimonials). Antimonials are metalloid based (Sb), highly toxic, and have severe adverse side effects including pancreatitis and cardiac arrhythmia.[1] Additionally, many strains of *L. donovani* have become resistant to antimonial therapies, demanding the development of new chemotherapeutics for VL treatment. Miltefosine and the antifungal drug amphotericin B (AmpB) have also been used for *Leishmania* treatment. Unfortunately, miltefosine is not a preferred drug because it is teratogenic. While AmpB is highly effective in treating leishmaniasis, it is highly toxic and requires encapsulation in a liposomal formulation (AmBisome). In addition, AmpB is hypothesized to act directly on the *Leishmania* by binding to ergosterol found in the membrane of *Leishmania*[2] and by preventing entry of the promastigote into the macrophage. This direct activity can drive selective pressure in the development of drug resistance towards the treatment. In fact, researchers have already isolated a strain of *Leishmania* that demonstrates resistance to AmpB.[3] It has also been observed that certain *Leishmania* strains have an inherent broad spectrum resistance against drugs they have not previously encountered, illustrating that there would be some resistance to any proposed parasite specific therapy.[4]

One strategy to overcome drug resistance is through use of a host-directed (also called host-targeted) therapy that improves the host cell’s ability to clear infection. This removes the direct pressure on the pathogen and can mitigate further drug resistance. Targeting the host rather than the pathogen can also potentially treat drug resistant strains.[5] Furthermore, because many pathogens take advantage of similar pathways, there is a potential for developing therapies that target a broad-spectrum of pathogens. OSU-03012, also known as AR-12, is an orally active pyruvate dehydrogenase kinase isoenzyme 1 (PDK-1) and protein kinase B (AKT) signaling pathway inhibitor that reached Phase 1 clinical trials as a potential cancer therapeutic. During development, AR-12 was found to induce autophagy, which has been shown to disrupt the life-cycle of some intracellular pathogens.[6, 7] Additionally, it inhibits expression of Glucose Regulated Protein (GRP78), which has been shown to be induced in *Leishmania* infected macrophages.[8] As a host-directed therapy against VL, our work has shown that in vitro and in vivo, AR-12 can decrease the parasite burden of *L. donovani* in infected macrophages while having no direct effect on the promastigote.[9] Additionally, co-treatment with a conventional therapeutic, AmpB, resulted in a significant reduction in parasite burden compared to encapsulated OSU-03012 or AmpB alone, indicating the potential of AR-12 in sensitizing parasites to AmpB.[10]

In this work, we have screened a re-purposed library of 343 compounds derived from the FDA IND approved cancer drug AR-12 for host-directed anti-leishmanial activity. Additionally, we used a medium-throughput luminescence-based screening assay for detection of intracellular *Leishmania* amastigote viability. This method was validated against more conventional image-based analysis (Giemsa stain). The culmination of this work (**Figure 1**) identified four novel lead compounds that are significantly improved host-directed therapies compared to parental compound AR-12. Furthermore, proteomic analysis identified a proposed mechanism of action for the most promising lead compound being linked to lysozyme and its interaction with parasite.

**Figure 1.**
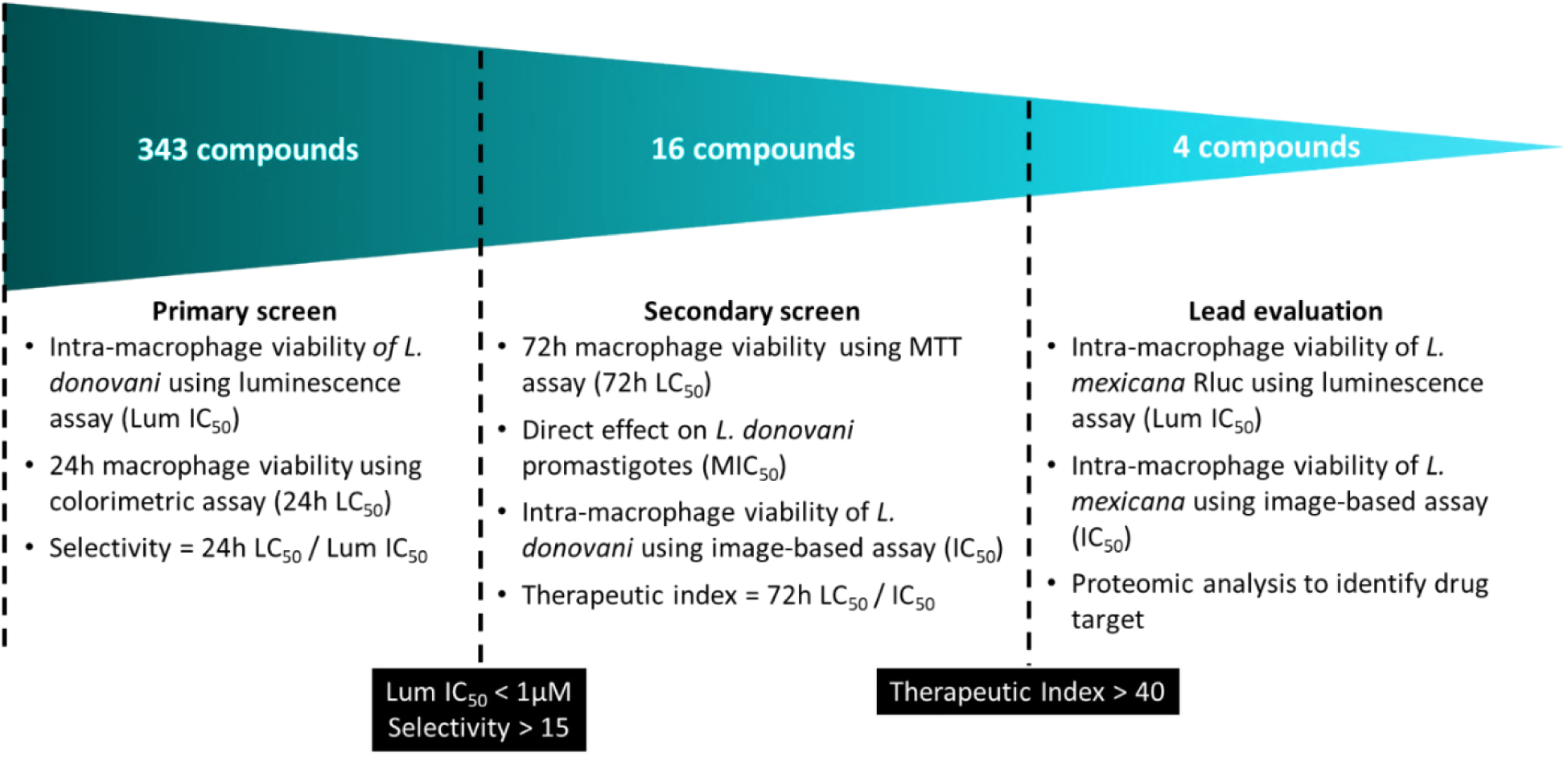
Process flow of screening methodology and lead compound selection. Concentration at which intracellular *Leishmania* burden is reduced by 50% in THP1 macrophages (Lum IC_50_) as identified by luminescence assay. Concentration where THP-1 macrophage cell viability is 50% (LC_50_) after incubation with compound as determined by MTT assay. Concentration at which intracellular *Leishmania* burden is reduced by 50% in bone marrow derived macrophages (IC_50_) as identified image-based Giemsa staining. Minimum inhibitory concentration (MIC) where extracellular *Leishmania* promastigote viability is reduced by 50% (MIC_50_) after 72-hour incubation with compound as measured by resazurin assay.

## RESULTS AND DISCUSSION

AR-12 (**Figure 2**) has been shown to have host-directed effects that decrease the parasite burden of *L. donovani* in infected macrophages.[9] AR-12 is based on an N1-aryl-3-trifluoromethyl pyrazole core structure, shown with variable R₁ (red) and R₂ (blue) positions. To explore structure–activity relationships, a focused library of 343 AR-12 analogs was synthesized with systematic substitutions at R₁ and R₂. The complete chemical structures of all compounds can be found in **Supplemental Table 1**. R₁ modifications included cyclohexyl, substituted benzene biphenyl derivatives. R₂ modifications encompass a chemically diverse set of glycinamide derivatives, 3-amino pyrrolidine, piperazine, N-benzyl pyrrolidine, and other elaborations on isosteres for the carboxyamide group of AR-12.

**Figure 2:**
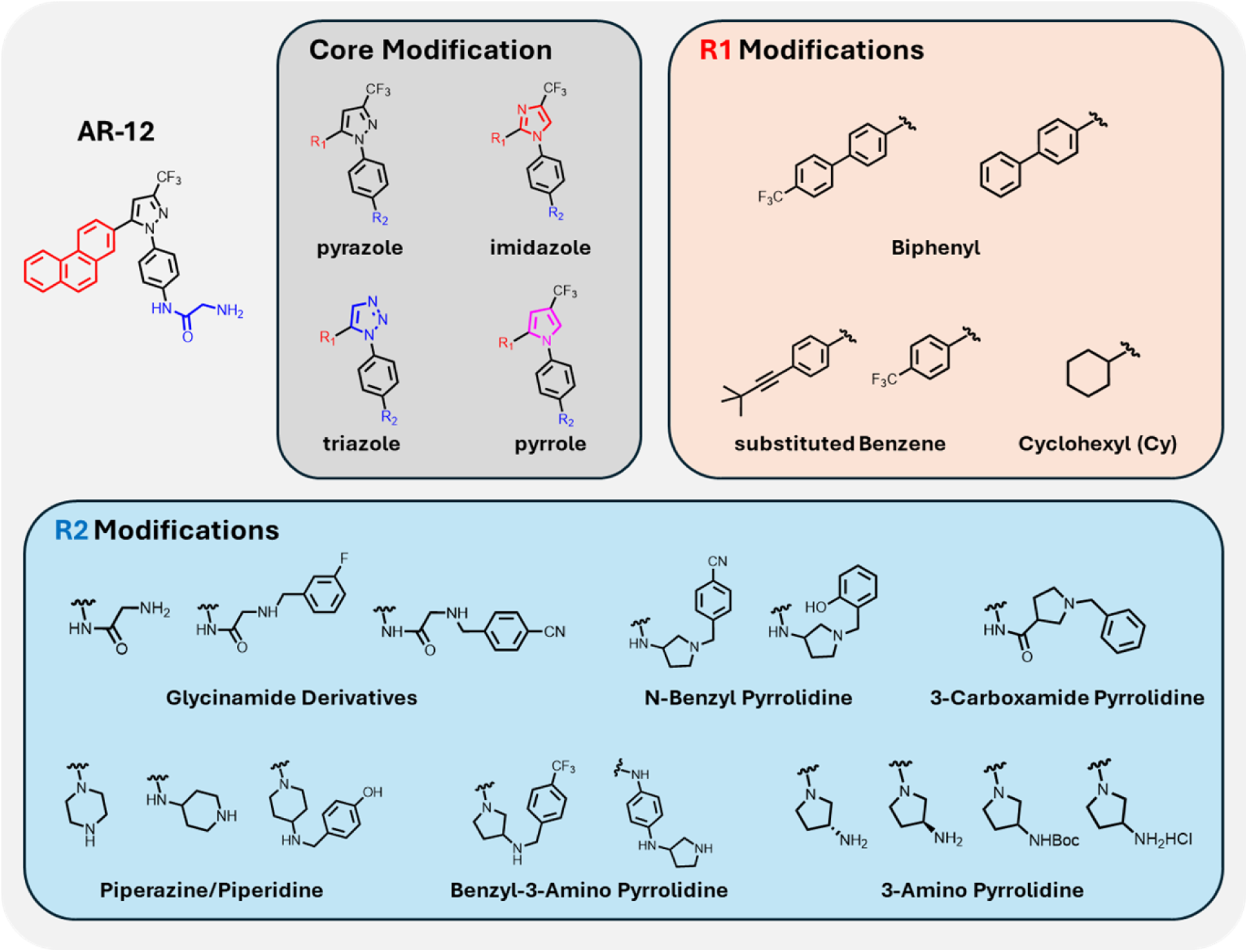
Schematic detailing workflow of structure activity relationship for chemical compounds derived from AR-12 with associated core, R1, and R2 modifications.

### Primary Screen

Primary screening of the 343 compounds was performed by medium throughput luminescence-based screening assay using THP-1 macrophages and luminescent *L. donovani* (**Supplemental Figure 1, Supplemental Table 2**). Additionally, the effect of the compounds on host cell viability was evaluated in uninfected THP-1 cells using a colorimetric assay. These two values were used to calculate selectivity of the compounds (24hr LC_50_ / Lum IC_50_). Comparing these compounds to AR-12 potency and toxicity to the host cell revealed 33 compounds both less cytotoxic and more potent (**Supplemental Figure 2**). These hits outline clear structure–activity trends. Biphenyl– and cyclohexyl-substituted pyrazoles emerged as favored scaffolds and select imidazole-core variants validated the conserved behavior of these motifs. Within R₂ substitutions, glycinamide derivatives (50, 158, 197, 354–355) consistently enhanced selectivity, while 3-amino pyrrolidines, particularly those decorated with polar benzyl groups (53, 91, 356, 362) increased potency, though sometimes at the expense of host tolerability. Importantly, modification of the pyrrolidine amine by protection or ionization (133, 134, 281) yielded solubility and selectivity advantages, suggesting an accessible synthetic handle within this series. Cyclohexyl R₁ analogs paired with piperidine or piperazine derivatives (324, 334, 336, 339, 370, 416–421) extended the structure-activity landscape, with alkoxypiperidines preserving activity while maintaining host viability. Changing the core-scaffold to imidazole series reinforced these trends, with compounds 154, 158, and 197 retaining potent intracellular activity. From these observations, sixteen compounds (**Table 1, Supplemental Figure 3-4**) were identified with either a high potency (Lum IC_50_< 1 µM) or a high selectivity (>15) and were selected for secondary screening.

**Table 1.** Hits identified by primary screen. Hits determined by potent host-directed activity (IC_50_ < 1 µM) or high selectivity (>15). Lum IC_50_ is the concentration at which intracellular *L. donovani* burden is reduced by 50% in THP1 macrophages as identified by luminescence assay. Concentration where THP-1 macrophage cell viability is 50% (LC_50_) after 24-hour incubation with compound as determined by MTT assay. Selectivity between host-directed effect and cytotoxicity, defined as 24h LC_50_ / Lum IC_50_. Parental compound AR-12 provided for reference.

|  | 24 hr viability of<br>THP-1<br>macrophages | Intracellular<br>*L. donovani* amastigotes | Selectivity |
| --- | --- | --- | --- |
| Compounds | 24h LC<sub>50</sub> (μM) | Lum IC<sub>50</sub> (μM) | 24h LC<sub>50</sub> /<br>Lum IC<sub>50</sub> |
| AR-12 | 13.3 | 3.4 | 3.9 |
| 23 | 5.8 | 0.4 | 14.8 |
| 25 | 28.9 | 0.7 | 43.8 |
| 44 | 7.4 | 0.9 | 8.1 |
| 50 | 23.5 | 1.5 | 16.0 |
| 53 | >50 | 0.4 | >112.6 |
| 86 | 21.9 | 1.4 | 16.0 |
| 91 | >50 | 1.4 | >36.1 |
| 129 | 6.3 | 0.1 | 46.7 |
| 130 | 4.2 | 0.2 | 23.9 |
| 133 | 29.0 | 0.9 | 32.5 |
| 134 | 22.6 | 0.5 | 42.1 |
| 158 | 16.0 | 0.2 | 81.0 |
| 197 | >50 | 2.4 | >20.6 |
| 319 | 6.4 | 0.8 | 8.1 |
| 354 | 22.0 | 0.5 | 44.9 |
| 408 | 25.4 | 1.5 | 17.0 |

### Secondary Screen

Additional screening of the selected 16 compounds was performed to confirm drug activity. Specifically, the effect of drugs on macrophage viability was evaluated over a longer range of time (72 hrs, **Supplemental Figure 5**). Furthermore, the direct effect of the compounds on extracellular *L. donovani* promastigotes was measured using resazurin assay (**Supplemental Figure 6**). Lastly, activity of compounds to reduce intracellular *L. donovani* was confirmed using Giemsa staining and image-based analysis in bone-marrow derived macrophages (BMDMs) was assessed (**Supplemental Figure 7**). The host-directed therapeutic index was calculated by the 72hr LC_50_ / IC_50_. This characterization is detailed in **Table 2** for all 16 compounds. Structure-activity analysis of the 16 candidates converged on four leads (53, 134, 197, and 354) that achieved therapeutic indices above 40. A strong preference for biphenyl substitution at R₁ was evident, appearing in three of the four leads, emphasizing its role as a privileged motif for potency and selectivity. Compound 53 exemplified how polar, electron-withdrawing groups can improve intracellular efficacy. In contrast, compound 197 demonstrated the successful transfer of these structural features to a new heteroaryl core. Cyclohexyl analog 354 confirmed that glycinamide R₂ groups can pair productively with alternative hydrophobic motifs, yielding a profile of balanced potency and host safety. Taken together, these results illustrate that optimal host-directed activity is achieved when hydrophobic R₁ scaffolds are complemented by solubilizing, electronically altered R₂ functionalities. These four leads were prioritized for tertiary screening in cutaneous strain, *L. mexicana,* to ensure broad host-directed activity.

**Table 2:** Further analysis of hit compounds. Concentration at which intracellular *L. donovani* burden is reduced by 50% in bone marrow derived macrophages (IC_50_) as identified image-based Giemsa staining. Minimum inhibitory concentration (MIC) where extracellular *L. donovani* promastigote viability is reduced by 50% (MIC_50_) after 72-hour incubation with compound as measured by resazurin assay. Concentration where THP-1 macrophage cell viability is 50% (LC_50_) after 72-hour incubation with compound as determined by MTT assay. Therapeutic index between host-directed effect and cytotoxicity, defined as LC_50_/IC_50_. Parental compound AR-12 provided for reference.

|  | Extracellular<br>*L. donovani*<br>promastigotes | Intracellular<br>*L. donovani*<br>amastigotes | 72 hr viability<br>of THP-1<br>macrophages | Therapeutic Index |
| --- | --- | --- | --- | --- |
| Compounds | MIC<sub>50</sub> (μM) | IC<sub>50</sub> (μM) | LC<sub>50</sub> (μM) | LC<sub>50</sub> / IC<sub>50</sub> |
| AR-12 | 2.2 | 2.2 | 8.6 | 4 |
| 23 | 4.0 | 0.9 | 6.9 | 8 |
| 25 | 2.1 | 2.5 | 6.3 | 2 |
| 44 | 1.9 | 1.0 | 14.5 | 14 |
| 50 | 14.6 | 9.1 | 42.7 | 5 |
| 53 | >200 | 1.1 | >300 | >273 |
| 86 | 3.2 | 0.6 | 8.2 | 14 |
| 91 | 56.3 | 2.4 | 53.4 | 22 |
| 129 | 1.7 | 0.4 | 6.2 | 17 |
| 130 | 2.1 | 1.6 | 6.4 | 4 |
| 133 | 13.0 | 1.7 | 33.9 | 20 |
| 134 | 5.7 | 0.7 | 32.0 | 46 |
| 158 | 4.6 | 0.6 | 16.5 | 29 |
| 197 | 6.4 | 1.1 | >300 | >273 |
| 319 | 0.8 | 3.4 | 7.2 | 2 |
| 354 | 22.1 | 0.4 | 20.2 | 51 |
| 408 | 0.9 | 2.6 | 25.1 | 10 |

### Tertiary Screen

The four lead compounds were then screened in *L. mexicana* using the same assays described for *L. donovani*. The ability of compounds to reduce intracellular *Leishmania* was evaluated by both luminescent and image-based assays. Additionally, the direct activity of compounds on extracellular *L. mexicana* promastigotes was measured using resazurin assay (**Figure 3**, **Table 3**). The ability of these four compounds to reduce intracellular burden was similar for both *L. mexicana* and *L. donovani* as measured by luminescent and image-based assays. However, the direct effect of compounds on promastigotes varied between the visceral and cutaneous strains. This suggests that the host-directed effect is the driving factor in intracellular pathogen clearance and is less susceptible to differences in *Leishmania* strains.

**Figure 3.**
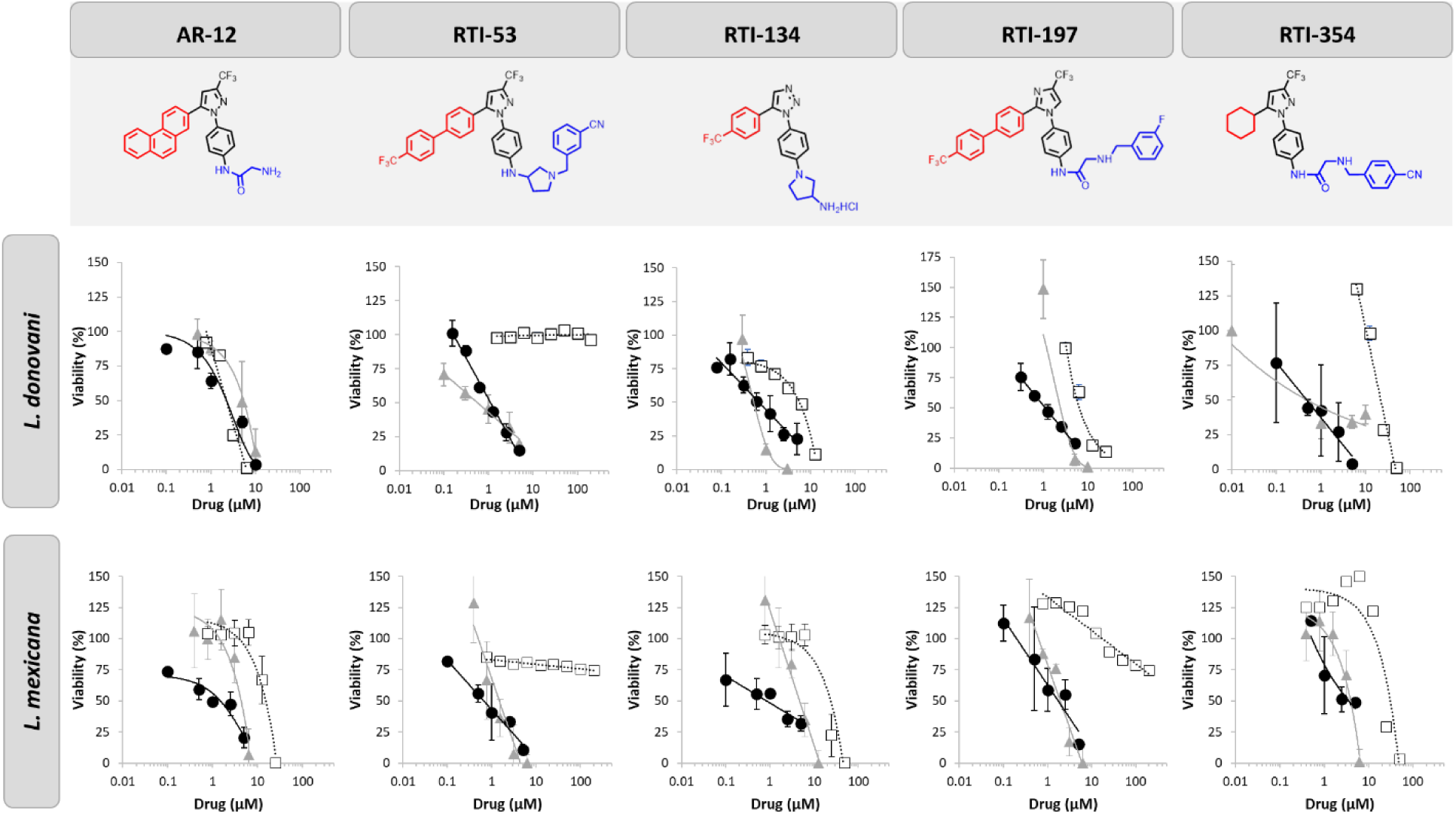
Lead compounds. Dose response of intracellular *Leishmania* spp. burden in THP-1 macrophages as identified by luminescence-based screen (gray triangle). Dose response of intracellular pathogen burden in bone marrow derived macrophages as identified image-based Giemsa staining (black circle). Dose response of extracellular promastigote viability after 72-hour incubation with compound as measured by resazurin assay (open square). Parental compound AR-12 provided for reference. Data is presented as mean ± standard deviation of biological triplicates.

**Table 3:**
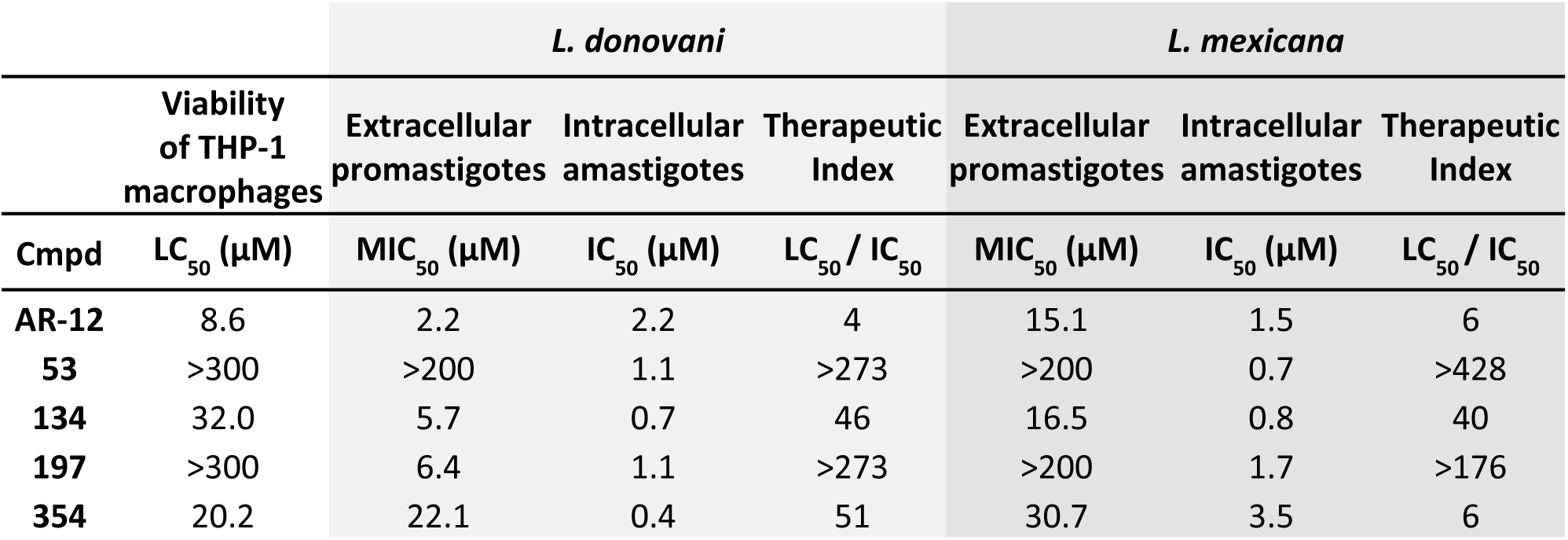
Comparison across cutaneous and visceral *Leishmania* strains. Concentration where THP-1 macrophage cell viability was 50% (LC_50_) after 72-hour incubation with compound as determined by MTT assay. Concentration at which intracellular *L. donovani* or *L. mexicana* burden was reduced by 50% in bone marrow derived macrophages (IC_50_) as identified image-based Giemsa staining. Minimum inhibitory concentration (MIC) where extracellular *L. donovani* or *L. mexicana* promastigote viability was reduced by 50% (MIC_50_) after 72-hour incubation with compound as measured by resazurin assay. Therapeutic index between host-directed effect and cytotoxicity, defined as LC_50_/IC_50_. Parental compound AR-12 provided for reference.

Tertiary screening revealed two compounds of interest: 53 and 197. Neither compound had any reduction in macrophage viability over a 72hr incubation up to 300µM (**Table 3**). However, both were able to decrease intracellular parasite burden by 50% with less than 2µM in both visceral and cutaneous *Leishmania* strains (**Table 3**, **Figure 3**). Interestingly, 53 had no direct effects on extracellular promastigote viability indicating that all anti-leishmanial effects are host-directed. While 197 does have some direct reduction in *L. donovani* extracellular promastigote viability, the host-directed effects are five-fold more significant and notably, 197 had no effect on *L. mexicana* extracellular promastigote viability.

### 197 Proteomic Analysis and Target Validation

Although the highest selectivity was noted with 53 in two strains, 197 displayed broad activity against the promastigote as well as host-directed activity, further the chemical ligation to 197 was more approachable than for 53. For these reasons we utilized two different proteomic strategies to identify the molecular target of 197. In one strategy, 197 was chemically modified (**Supplemental Figure 8A**) and conjugated to agarose beads using a 5-carbon spacer (termed ‘197-bead’). Chemical modification did not affect 197 anti-*Leishmania* activity (**Supplemental Figure 8B**). We also prepared a bead conjugated to butylamine to serve as a control for nonspecific interactions (termed ‘control-bead’). The beads were then incubated with *Leishmania* infected macrophage lysate. Afterwards, the beads were washed and boiled in sample buffer to release bound proteins which were then evaluated by LC-MS/MS. This analysis identified 841 human and 65 *Leishmania* enriched proteins on the 197-beads compared to control beads. These enriched proteins all had a Log2 fold-change >1 and p < 0.05 (**Supplemental Figure 9A)**.

In a second strategy, the thermal proteome profiling (TPP) approach was used to identify the proteins in *Leishmania* infected macrophage lysate that displayed a 197-induced change in thermal stability. The TPP approach is an attractive method by which to identify the direct (and indirect) targets of protein-ligands including small molecule drugs. The TPP experiment performed here effectively assayed over 1600 human proteins and over 1000 *Leishmania* proteins for binding to 197. A total of 123 human and 71 *Leishmania* proteins were identified with 197-induced changes in their thermal stability using selection criteria analogous to that used in the affinity pull-down experiment (i.e., a |z-score| > 1 and a p < 0.05) (**Supplemental Figure 9B**).

Cross-referencing the 841 significant human proteins from the affinity capture and 123 human proteins from TPP analysis revealed 26 overlapping proteins (**Figure 4 A-B**). Further evaluating strength of interaction, refining affinity capture proteins to Log2 fold-change > 4 and TPP proteins to |z-score| > 4 revealed one protein: lysozyme (**Figure 4 A-B**). The overlap between the 65 and 71 *Leishmania* proteins identified in affinity and TPP analysis, respectively, was also investigated revealing three overlapping proteins (**Supplemental Figure 9C-D**). Further evaluating strength of interaction, refining affinity capture proteins to Log2 fold-change > 4 and TPP proteins to |z-score| > 4 revealed 0 proteins (**Supplemental Figure 9E**). These results further support the idea that 197 is acting in a host-directed manner.

**Figure 4.**
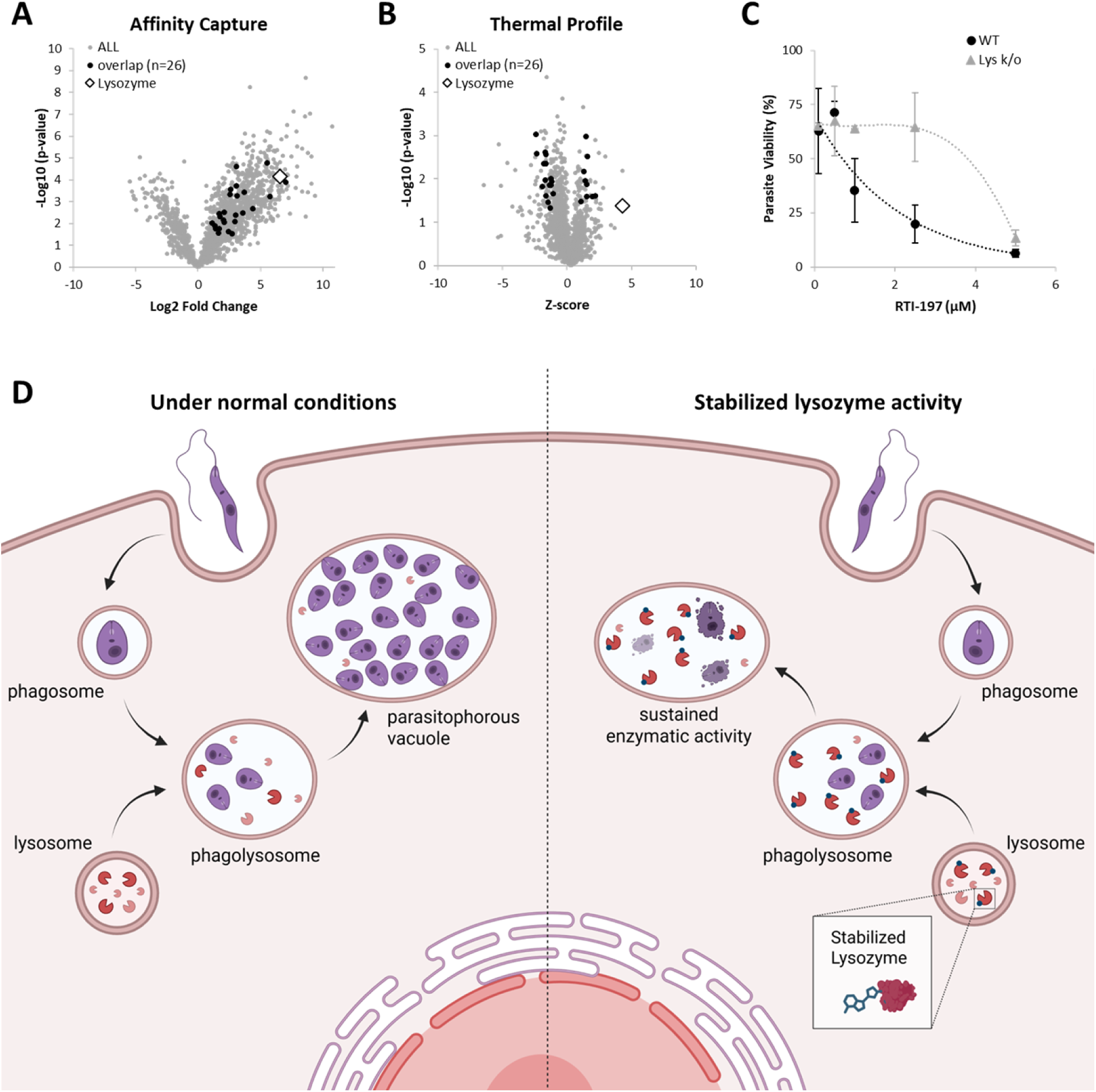
Proteomic Analysis. **A)** All human proteins (gray circle) identified by affinity capture using 197 functionalized bead plotted Log2(fold-change) and –Log10(p-value) over control bead. Proteins with a Log2FC > 1 and p < 0.05 overlapping with significant proteins identified by thermal profile analysis (|z-score| > 1 and p < 0.05) are shown with black circles. Lysozyme protein identified by white diamond. **B)** All human proteins (gray circle) identified by thermal profile analysis with 197. Proteins with a (|z-score| > 1 and p< 0.05 overlapping with significant proteins identified by affinity capture (Log2FC > 2 and p< 0.05) are shown with black circles. Lysozyme protein identified by white diamond. **C**) Dose response of 197 on intracellular *Leishmania* burden in bone marrow derived macrophages derived from wildtype C57BL/6 (black circle) or lysozyme knockout mice (gray triangle) as identified image-based Giemsa staining. Data is presented as mean ± standard deviation of biological triplicates. **D)** Schematic of proposed mechanism. Under normal conditions (left), *Leishmania* promastigote is internalized into a phagosome which matures into a phagolysosome by fusing with a lysosome. *Leishmania* amastigotes interfere with maturation, remodeling the phagolysosome into a parasitophorous vacuole permissive for parasite replication. With host-lysozyme stabilized by RTI-197 (right), phagolysosome maturation proceeds and enzymatic activity is maintained allowing host cell degradation of amastigotes.

To evaluate lysozyme as a mechanistic target for 197, the ability of 197 to eradicate parasitic burden from BMDM cells derived from both wild-type (WT) and lysozyme knockout (Lys K/O) mice were evaluated. Amphotericin B, which has a different mechanism of action, acting on membrane sterols to decrease permeability barrier to small metabolites, was utilized as a control.[11] Amphotericin B retained its anti-leishmanial activity in both WT and Lys K/O BMDMs (**Supplemental Figure 10**); however, 197 activity was notably decreased in Lys K/O BMDMs (**Figure 4C**). Specifically, the IC_50_ for 197 in Lys K/O BMDMs was more than five-fold greater than in WT BMDMs. This supports proteomic analysis that lysozyme is a target of 197 and plays a role in host-directed anti-*Leishmania* effect.

Lysozyme is a ∼14 kDa enzyme present in mucosal secretions and tissues of animals. It is also present in cytoplasmic granules of macrophages and the polymorphonuclear neutrophils and plays role in innate immunity. Lysozyme catalyzes the hydrolysis of 1,4-beta-linkages in peptidoglycan, which is a major component of gram-positive bacterial cell wall, resulting in bacterial lysis. Lysozyme has recently been explored as an alternative to antibiotics.[12] While lysozyme is primarily known for its antibacterial properties, Valigurova et al. has shown that lysosomal endocytosis plays a role in *Leishmania* host cell infection.[13] Most notably, Kumar et al. concluded that low lysozyme activity in patients may account for persistence of *Leishmania* parasites in VL infections.[14]

Under normal conditions, the *Leishmania* promastigote is internalized into a phagosome which matures into a phagolysosome by fusing with a lysosome. *Leishmania* amastigotes interfere with phagolysosome maturation, remodeling it into a parasitophorous vacuole permissive for parasite replication. TPP analysis of 197 had a strongly positive z-score of 4.32 for lysozyme. A positive z-score typically indicates increased protein stability often due to interactions such as ligand binding, post-translational modifications, or complex formation.[15, 16] This TPP result is consistent with the affinity capture finding of an increase in abundance of lysozyme (Log2 fold change = 6.57). These results both suggest that 197 helps stabilize the enzyme lysozyme. Mechanistically, this host-modification could help reduce intracellular infection by preventing *Leishmania*-induced remodeling of the phagolysosome into a parasitophorous vacuole. This prevents the formation of the permissive niche for parasitic replication and allows destruction of internalized amastigotes by the host cell (**Figure 4D**).

These findings not only advance our understanding of the molecular mechanisms underlying anti-leishmanial activity of our analogs, confirming host-directed and pathogen directed activity, but pave the way for the development of novel therapeutic strategies to combat this pervasive disease. Future research should focus on optimizing these lead compounds and exploring their potential in clinical settings to address the urgent need for effective leishmaniasis treatments.

## METHODS AND MATERIALS

### Synthetic Methods for Hit Compounds

Below is the synthesis for the 16 hit compounds, a more complete methods and characterization can be found at the end of the **Supplemental Information**.

**<u>RTI-23:</u>** A solution of **RTI-7** (150 mg, 0.29 mmol), 4-nitrobenzaldehyde (44 mg, 0.29 mmol) and 4A molecular sieves (150 mg) in anhydrous methanol (3 mL) and anhydrous tetrahydrofuran (1.5 mL) was stirred at room temperature for 18 hours. The reaction was cooled to 0° C and treated with sodium borohydride (22 mg, 0.58 mmol) was added and the reaction stirred for 4 hours at room temperature. The reaction was concentrated and the residue partitioned between saturated aqueous sodium bicarbonate solution and ethyl acetate. The combined organic layers were washed with brine, dried (Na_2_SO_4_), filtered and concentrated to yield 203.6 mg of a brown gel. The crude material was purified over silica gel using 0-10 % methanol from dichloromethane to yield 65 mg (36%) of **RTI-23** as an off-white solid. ^1^H-NMR (CDCl_3_) δ 7.69 (s, 4 H), 7.56 (d, 2 H, J = 9 Hz), 7.34 (d, 2 H, J = 6 Hz), 7.14 (dd, 4 H, J = 3 Hz, 9 Hz), 6.77 (s, 1 H), 6.66 (d, 2 H, J = 6 Hz), 6.47 (d, 2 H, J = 9 Hz), 3.76 (s, 2 H), 3.63-3.41 (m, 6 H), 3.39-3.29 (m, 1 H), 3.19-3.08 (m, 1 H), 2.31-2.19 (m, 1 H), 1.99-1.88 (m, 1 H). ESI-MS, calculated for C_34_H_27_F_6_N_5_O_2_ (MH)^+^ 622.6; observed 622.3.

**<u>RTI-25:</u>** A solution of **RTI-15** (4.90 g, 7.96 mmol) in dichloromethane (55 mL) was cooled to 0°C and treated with trifluoroacetic acid (5.9 mL, 79.4 mmol). The reaction warmed to room temperature and stirred for 18 hours. Upon completion, the mixture was concentrated and the residue was partitioned between ethyl acetate and 2 N aqueous NaOH. The aqueous layer was extracted with ethyl acetate and the combined organic layers were washed with brine, dried (Na_2_SO_4_), filtered and concentrated to yield a brown solid (**RTI-25**, 4.0 g, 97%) that required no further purification. ^1^H-NMR (CDCl_3_) δ 7.68 (dd, 4 H, J = 9 Hz), 7.56 (d, 2 H, J = 9 Hz), 7.36 (d, 2 H, J = 9 Hz), 7.14 (d, 2 H, J = 9 Hz), 6.78 (s, 1 H), 6.56 (d, 2 H, J = 9 Hz), 4.05-3.91 (m, 2 H), 3.21-3.08 (m, 2 H), 3.01-2.93 (m, 1 H), 2.91-2.85 (m, 1 H), 2.27-2.15 (m, 1 H). Anal. Calculated (with 0.8 mol of water) for C_27_H_22_F_6_N_4_; C, 61.08; H, 4.48; N, 10.55. Found: C, 61.23; H, 4.30; N, 10.43.

**<u>RTI-44:</u>** A solution of **28** (34 mg, 0.061 mmol) in dichloromethane (2 mL) was cooled to 0° C and treated with trifluoroacetic acid (0.2 mL, 2.69 mmol). The reaction warmed to room temperature and stirred for 15 hours. Upon completion, the mixture was concentrated and the residue was partitioned between ethyl acetate and 2 N aqueous NaOH. The aqueous layer was extracted with ethyl acetate and the combined organic layers were washed with brine, dried (Na_2_SO_4_), filtered and concentrated to yield 281 mg of **RTI-44** (>100%) of a yellow gel, which required no further purification. ^1^H-NMR (CD_3_OD) 7.67-7.61 (m, 2 H), 7.58-7.52 (m, 2 H), 7.30-7.17 (m, 4 H), 3.49-3.43 (m, 4 H), 3.37-3.32 (m, 4 H), 1.27 (s, 9 H).

**<u>RTI-50</u>:** Following the procedures for preparing **RTI-77** and **RTI-79**, **RTI-50** was isolated as an off-white solid (264 mg, 79% for the final step). ^1^H-NMR (CDCl_3_) δ 9.66 (br s, 1 H), 7.74 (d, 2 H, J = 9 Hz), 7.57 (d, 2 H, J = 9 Hz), 7.50 (dd, 4 H, J = 9 Hz), 7.43 (dd, 2 H, J = 6 Hz, 9 Hz), 7.36 (d, 1 H, J = 9 Hz), 7.32-7.24 (m, 3 H), 3.52 (s, 2 H). ESI-MS, calculated for C_24_H_19_F_3_N_4_O (MH)^+^ 437.4; observed 437.0. Anal. Calculated (with 0.2 mol water) for C_24_H_19_F_3_N_4_O; C, 65.50; H, 4.44; N, 12.73. Found: C, 66.04; H, 4.38; N, 12.83.

**<u>RTI-86</u>:** A mixture containing **11** (400 mg, 1.05 mmol), 1-(Phenylmethyl)-3-pyrrolidinecarboxylic acid (325 mg, 1.58 mmol), diisopropylethylamine (0.65 mL, 3.73 mmol) and propylphosphonic anhydride solution, 50 wt. % in ethyl acetate (1.9 mL, 3.19 mmol) in anhydrous tetrahydrofuran (38 mL) was sealed tightly and stirred at room temperature for 18 hours. The solvent was concentrated to 20% volume and the residue was partitioned between ethyl acetate and saturated aqueous NaHCO_3_. The organic layer was washed with brine, dried (Na_2_SO_4_), filtered and concentrated. The crude material was adsorbed onto silica gel and purified via ISCO using 0-10% methanol from dichloromethane to yield 513 mg (86%) of an off-white solid (**RTI-86**). ^1^H-NMR (CDCl_3_) δ 9.77 (s, 1 H), 7.57 (dd, 4 H, J = 6 Hz, 9 Hz), 7.48 (dd, 6 H, J = 6 Hz), 7.42 (d, 1 H, J = 6 Hz), 7.37-7.32 (m, 5 H), 7.29-7.20 (m, 4 H), 3.73 (dd, 2 H, J = 6 Hz, 12 Hz), 3.16 (dd, 2 H, J = 6 Hz, 9 Hz), 2.95 (dd, 1 H, J = 6 Hz, 9 Hz), 2.42-2.33 (m, 3 H), 2.12-2.05 (m, 1 H). LC-MS, calculated for C_34_H_29_F_3_N_4_O (MH)^+^ 567.6; observed 567.2. Anal. Calculated for C_34_H_29_F_3_N_4_O; C, 72.07; H, 5.15; N, 9.88. Found: C, 71.97; H, 5.25; N, 9.82.

**<u>RTI-53 [R = CH_2_-(3-cyanophenyl)]:</u>** Using 3-cyanobenzaldehyde, the product was isolated as a white solid in 58% yield (142 mg). ^1^H-NMR (CDCl_3_) δ 7.65 (dd, 5 H, J = 12 Hz), 7.55 (d, 2 H, J = 9 Hz), 7.43 (m, d, 1 H, J = 6 Hz), 7.35 (dd, 2 H, J = 9 Hz), 7.14 (dd, 2 H, J = 9 Hz), 6.77 (s, 1 H), 6.55 (d, 2 H, J = 9 Hz), 4.16-4.03 (m, 2 H), 3.65 (s, 2 H), 2.84-2.71 (m, 2 H), 2.59 (dd, 1 H, J = 9 Hz), 2.46-2.28 (m, 2 H), 1.76-1.65 (m, 1 H). ESI-MS, calculated for C_35_H_27_F_6_N_5_ (MH)^+^ 632.6; observed 632.6.

**<u>RTI-91 [R = CH_2_-(2-hydroxyphenyl)]:</u>** Using 2-hydroxybenzaldehyde, the product was isolated as an off-white solid in 48% yield (138.6 mg). ^1^H-NMR (CDCl_3_) δ 7.68 (dd, 4 H, J = 9 Hz), 7.56 (d, 2 H, J = 9 Hz), 7.35 (dd, 2 H, J = 9 Hz), 7.16 (dd, 3 H, J = 6 Hz, 9 Hz), 6.99 (d, 1 H, J = 9 Hz), 6.84-6.76 (m, 3 H), 6.53 (d, 2 H, J = 9 Hz), 4.06-3.97 (m, 2 H), 3.84 (s, 2 H), 2.97-2.85 (m, 2 H), 2.70 (dd, 1 H, J = 3 Hz, 6 Hz), 2.56 (dd, 1 H, J = 9 Hz), 2.42 (dd, 1 H, J = 6 Hz, 9 Hz), 1.78-1.72 (m, 1 H). ESI-MS, calculated for C_34_H_28_F_6_N_4_O (MH)^+^ 623.6; observed 623.8.

**<u>RTI-129 (5b: R-amino orientation):</u>** The product was isolated as a brown gel in 59 % yield (160 mg). ^1^H-NMR (CDCl_3_) δ 7.69 (s, 4 H), 7.56 (d, 2 H, J = 9 Hz), 7.35 (d, 2 H, J = 9 Hz), 7.17 (d, 2 H, J = 9 Hz), 6.78 (s, 1 H), 6.49 (d, 2 H, J = 9 Hz), 3.76 (dd, 1 H, J = 3 Hz, 6 Hz), 3.55-3.45 (m, 2 H), 3.38-3.30 (m, 1 H), 3.04 (dd, 1 H, J = 3 Hz, 6 Hz), 2.29-2.18 (m, 1 H), 1.88-1.78 (m, 1 H). ESI-MS, calculated for C_27_H_22_F_6_N_4_ (MH)^+^ 517.4; observed 517.6; Anal. Calculated for C_27_H_22_F_6_N_4_; C, 62.79; H, 4.29; N, 10.84. Found: C, 62.69; H, 4.37; N, 10.55; [α] = + 4.28 (c = 0.70/CHCl_3_).

**<u>RTI-130 (5c: S-amino orientation):</u>** The product was isolated as a tan solid in 82 % yield (335 mg). ^1^H-NMR (CDCl_3_) δ 7.66 (s, 4 H), 7.56 (d, 2 H, J = 9 Hz), 7.35 (d, 2 H, J = 9 Hz), 7.17 (d, 2 H, J = 9 Hz), 6.78 (s, 1 H), 6.49 (d, 2 H, J = 9 Hz), 3.75 (dd, 1 H, J = 6 Hz), 3.55-3.45 (m, 2 H), 3.38-3.30 (dd, 1 H, J = 6 Hz, 9 Hz), 3.04 (dd, 1 H, J = 3 Hz, 6 Hz), 2.29-2.18 (m, 1 H), 1.88-1.78 (m, 1 H). ESI-MS, calculated for C_27_H_22_F_6_N_4_ (MH)^+^ 517.4; observed 517.6; Anal. Calculated (with 0.2 mol water) for C_27_H_22_F_6_N_4_; C, 62.35; H, 4.34; N, 10.77. Found: C, 62.16; H, 4.37; N, 10.65; [α] = – 2.50 (c = 0.80/CHCl_3_).

**<u>RTI-133:</u>** The following were combined in a heavy-duty glass reactor: **20** (110 mg, 0.3 mmol), 3-N-Boc-aminopyrrolidine (72.4 mg, 0.39 mmol), BINAP (56 mg, 0.09 mmol), Pd_2_(dba)_3_ (27.5 mg, 0.03 mmol) and Cs_2_CO_3_ (127 mg, 0.39 mmol) in anhydrous toluene (3 mL) and nitrogen gas was bubbled into the mixture for two minutes. The reactor was then sealed with a Teflon cap and heated to 110° C for 15 hours. Upon cooling, the mixture was filtered through Celite and the filter pad was rinsed with ethyl acetate. The filtrate was washed with water and brine, dried (Na_2_SO_4_), filtered and concentrated. The crude material was purified over silica gel using 0-100% ethyl acetate from hexanes to yield 158.5 mg (>100%) of a yellow solid (**RTI-133**).

**<u>RTI-134:</u>** A solution of **RTI-133** (158.5 mg, 0.334 mmol) in dichloromethane (20 mL) was cooled to 0° C and treated with trifluoroacetic acid (0.5 mL, 6.73 mmol). The reaction warmed to room temperature and stirred for 15 hours. Upon completion, the mixture was concentrated and the residue was partitioned between ethyl acetate and 2 N aqueous NaOH. The aqueous layer was extracted with ethyl acetate and the combined organic layers were washed with brine, dried (Na_2_SO_4_), filtered and concentrated to yield 112.5 mg (90%) of a yellow solid, which required no further purification. This material was dissolved in diethyl ether and treated with 2 N HCl/diethyl ether, stirred at room temperature for 18 hours, filtered, washed with diethyl ether and dried to yield 104 mg (76%) of **RTI-134** as a white solid. ^1^H-NMR (CD_3_OD) δ 8.08 (s, 1 H), 7.63 (d, 2 H, J = 8.0 Hz), 7.48 (d, 2 H, J = 8.4 Hz), 7.21 (d, 2 H, J = 8.8 Hz), 6.71 (d, 2 H, J = 8.8 Hz), 4.09-4.01 (m, 1 H), 3.72-3.61 (m, 2 H), 3.49-3.42 (m, 2 H), 2.54-2.44 (m, 2 H). LC-MS, calculated for C_19_H_18_F_3_N_5_ (MH)^+^ 374.4; observed 374.2. Anal. Calculated (with 0.4 mol diethyl ether) for C_19_H_19_ClF_3_N_5_; C, 54.62; H, 5.45; N, 15.46. Found: C, 54.64; H, 5.05; N, 15.05.

**<u>RTI-158:</u>** A mixture containing **14** (50 mg, 0.132 mmol), Boc-glycine (69 mg, 0.394 mmol), diisopropylethylamine (0.15 mL, 0.792 mmol) and Propylphosphonic anhydride solution, 50 wt. % in ethyl acetate (0.25 mL, 0.394 mmol) in anhydrous tetrahydrofuran (20 mL) was sealed tightly and stirred at room temperature for 18 hours. The solvent was diluted with ethyl acetate (20 mL) and saturated aqueous NaHCO_3_ (25 mL). The organic layer was washed with brine, dried (Na_2_SO_4_), filtered and concentrated. The crude material was adsorbed onto silica gel and purified via ISCO using 10-50% ethyl acetate from hexanes to yield 50 mg (0.0933 mmol, 71%) of an off-white solid, which was dissolved in methanol (5 mL), cooled to 0° C and treated with 4.0 N hydrochloric acid in dioxane (1.2 mL, 4.8 mmol). The reaction warmed to room temperature and stirred for 18 hours. Upon completion, the mixture was concentrated and the residue was partitioned between dichloromethane and saturated aqueous NaHCO_3_. The aqueous layer was extracted with dichloromethane and the combined organic layers were washed with brine, dried (Na_2_SO_4_), filtered and concentrated. The crude material was adsorbed onto silica gel and purified via ISCO using 0-10% methanol from dichloromethane to yield 25 mg (63%) of an off-white solid (**RTI-158**). ^1^H-NMR (CDCl_3_) δ 9.60 (br s, 1 H), 7.68 (dd, 4 H, J = 8.4 Hz, 9.6 Hz), 7.49 (dd, 4 H, J = 2.0 Hz), 7.29-7.21 (m, 6 H), 3.50 (s, 2 H).

**<u>RTI-197:</u>** A mixture containing **RTI-79** (405 mg, 0.805 mmol), 3-fluorobenzyl bromide (0.11 mL, 0.897 mmol) and triethylamine (0.28 mL, 2.01 mmol) in dimethylformamide (7 mL) was stirred for 25 hours at room temperature. The reaction was poured into a saturated aqueous LiCl solution and extracted with ethyl ether. The organic layer was washed with brine, dried (Na_2_SO_4_), filtered and concentrated to obtain 523 mg of a yellow gel. The crude material was purified over silica gel using 0-5% methanol from dichloromethane to yield 212 mg (43%) of an off-white solid (**RTI-197**). ^1^H-NMR (CDCl_3_) δ 9.39 (s, 1 H), 7.71-7.64 (m, 6 H), 7.54-7.47 (m, 6 H), 7.37-7.23 (m, 3 H), 7.11-6.96 (m, 3 H), 3.88 (s, 2 H), 3.47 (s, 2 H). LC-MS, calculated for C_32_H_23_F_7_N_4_O (MH)^+^ 613.5; observed 613.2. Anal. Calculated for C_32_H_23_F_7_N_4_O; C, 62.74; H, 3.78; N, 9.14. Found: C, 62.46; H, 3.86; N, 9.09.

**<u>RTI-319:</u>** A solution of **23** (262 mg, 0.667 mmol), 4-hydroxybenzaldehyde (83 mg, 0.68 mmol) and 4A molecular sieves (300 mg) in anhydrous methanol (7.5 mL) and anhydrous tetrahydrofuran (3.5 mL), was stirred at room temperature for 18 hours. The reaction was cooled to 0° C and treated with sodium borohydride (51 mg, 1.33 mmol); the reaction stirred for 4 hours at room temperature. The reaction was concentrated and the residue partitioned between saturated aqueous sodium bicarbonate solution and ethyl acetate. The combined organic layers were washed with brine, dried (Na_2_SO_4_), filtered and concentrated to yield 390 mg of a yellow gel. The crude material was purified over silica gel using 0.5-5 % methanol from dichloromethane to yield 146 mg (44%) of **RTI-319** as a white solid. ^1^H-NMR (CDCl_3_) δ 7.30 (d, 2 H, J = 9 Hz), 7.17 (d, 2 H, J = 9 Hz), 6.93 (d, 2 H, J = 9 Hz), 6.72 (d, 2 H, J = 9 Hz), 6.55 (s, 1 H), 3.77 (s, 2 H), 3.72 (dd, 2 H, J = 12 Hz), 2.80 (dd, 2 H, J = 12 Hz), 2.74-2.65 (m, 2 H), 2.36 (br s, 1 H), 2.02 (d, 4 H, J = 12 Hz), 1.89-1.71 (m, 3 H), 1.62-1.22 (m, 7 H). ^13^C NMR (CDCl_3_, 75 MHz) δ 151.5, 129.4, 126.5, 115.6, 115.4, 54.0, 50.2, 47.9, 37.3, 33.0, 32.1, 26.2, 26.0; ESI-MS, calculated for C_28_H_33_F_3_N_4_O (M)^-^ 497.6; observed 497.2. Anal. Calculated for C_28_H_33_F_3_N_4_O; C, 67.45; H, 6.67; N, 11.23. Found: C, 67.19; H, 6.63; N, 11.11.

**<u>RTI-354:</u>** A mixture containing **RTI-341** (300 mg, 0.82 mmol), 4-cyanobenzyl bromide (164 mg, 0.835 mmol) and triethylamine (0.29 mL, 2.08 mmol) in dimethylformamide (7 mL) was stirred for 18 hours at room temperature. The reaction was poured into a saturated aqueous LiCl solution and extracted with ethyl ether. The organic layer was washed with brine, dried (Na_2_SO_4_), filtered and concentrated. The crude material was purified over silica gel using 1-5% methanol from dichloromethane to yield 212 mg (54%) of a white solid (**RTI-354**). ^1^H-NMR (CDCl_3_) δ 9.19 (s, 1 H), 7.68 (d, 2 H, J = 9 Hz), 7.45 (dd, 2 H, J = 6 Hz, 9 Hz), 6.60 (s, 1 H), 3.95 (s, 2 H), 3.46 (s, 2 H), 2.72 (dd, 1 H, J 3 Hz, 6 Hz), 2.07-1.99 (m, 2 H), 1.85-1.72 (m, 5 H), 1.52-1.25 (m, 6 H). ^13^C NMR (CDCl_3_, 75 MHz) δ 169.0, 158.4, 144.2, 137.9, 135.2, 132.6, 128.6, 126.4, 119.4, 111.6, 106.0, 53.5, 52.4, 37.3, 33.0, 26.2, 25.9; ESI-MS, calculated for C_26_H_26_F_3_N_5_O (MH)^+^ 482.5; observed 482.0. Anal. Calculated for C_26_H_26_F_3_N_5_O; C, 64.85; H, 5.44; N, 14.54. Found: C, 64.56; H, 5.51; N, 14.48.

**<u>RTI-408:</u>** A mixture containing **17** (100 mg, 0.269 mmol), tert-Butyl 4-oxopiperidine-1-carboxylate (160 mg, 0.809 mmol) and anhydrous sodium sulfate (catalytic amount) in acetic acid (5 mL) was stirred at room temperature for 2 hours. Sodium triacetoxyborohydride (360 mg, 1.61 mmol) was added and the reaction stirred at room temperature for 18 hours. The reaction was concentrated and the residue was partitioned between ethyl acetate and saturated aqueous NaHCO_3_. The crude material was adsorbed onto silica gel and purified via ISCO using 10-75% ethyl acetate from hexanes to yield 100 mg (0.181 mmol, 67%) of an off-white solid, which was dissolved in anhydrous dioxane (5 mL), cooled to 0° C and treated with 4.0 N hydrochloric acid in dioxane (2.25 mL, 9.03 mmol). The reaction warmed to room temperature and stirred for 18 hours. Upon completion, the mixture was concentrated and the residue was partitioned between dichloromethane and saturated aqueous NaHCO_3_. The aqueous layer was extracted with dichloromethane and the combined organic layers were washed with brine, dried (Na_2_SO_4_), filtered and concentrated to obtain 25 mg (30%) of an off-white solid (**RTI-408**). ^1^H-NMR (CDCl_3_) δ 7.56 (d, 2 H, J = 8.0 Hz), 7.51 (d, 2 H, J = 8.4 Hz), 7.41 (s, 1 H), 7.00 (dd, 2 H, J = 6.8 Hz, 8.8 Hz), 6.62 (dd, 2 H, J = 8.8 Hz, 9.6 Hz), 3.78-3.61 (m, 4 H), 3.27-3.17 (m, 2 H), 3.12-3.01 (m, 1 H), 2.36-2.23 (m, 2 H).

### Mammalian and Parasitic Cell Lines

Human monocytes, THP-1 (ATCC, TIB002) were used as host cells for *Leishmania* spp. infection and to assess drug cytotoxicity. The cells were cultured at 37C, 5% CO_2_ in RPMI medium (ATCC) supplemented with 10% FBS, 1% Penicillin/Streptomycin and 0.05 mM of β-mercaptoethanol. Cells were used in experiments up to passage 10.

Luminescent strains, *L. donovani* LV82 expressing firefly luciferase (a kind gift from Dr. Abhay Satoskar, Ohio State University) and *L. mexicana* (NR-51210, ATCC) expressing renilla luciferase were used to evaluate the effect of the compounds in intracellular and extracellular conditions[17]. Wild-type *L. donovani* LV82 (ATCC) were used for Giemsa staining based experiments. All parasites were cultured at 25°C and 5% CO_2_ in M199 media (Corning) supplemented with 10% FBS, 1% penicillin/streptomycin, and hemin (0.01 mg/mL), and used in experiments up to passage 15.

### Luminescent-based Evaluation of Intracellular Anti-Leishmanial Activity

THP-1 cells were seeded in a 96 well plate (25,000 cells/well) overnight then differentiated with 150 nM phorbol 12-myristate 13-acetate (PMA) over 72 hours (Supplemental Figure 1). The resulting macrophages were infected with *Leishmania* promastigotes at a multiplicity of infection of 1:10. Briefly, promastigotes of a known cell density were resuspended in RPMI media (with 10% FBS, 1% P/S and 0.05 mM β-mercaptoethanol) and incubated with adhered THP-1 cells over 18 hours. After infection, cells were washed three times with fresh media to remove extracellular promastigotes.

Infected macrophages were then treated with compounds solubilized in DMSO ranging from 0.1-10 µM in concentrations. After 72 hours of treatment, the viability of the amastigotes within macrophages was determined using Promega firefly luminescence assay (Cat E1500) or Pierce Renilla luciferase assay (Cat 16166) for *L. donovani* and *L. mexicana* strains, respectively. Briefly, media was removed, and cells were lysed with the assay lysis buffer. The luciferase substrate was then incubated with the lysed cells for 10 min at room temperature and luminescence of live *Leishmania* was measured in a white opaque 96 well plate using a Biotek plate reader. The IC_50_ value for each compound was obtained from the best fit curves obtained by plotting the relative luminescence units against the drug concentration.

### Effect of Compounds on Host Cell Viability

Effect of the compounds on THP-1 cell viability measured using thiazolyl blue tetrazolium bromide which measures cell metabolic activity (MTT). THP-1 cells were seeded in a 96 well plate (25,000 cells/well) overnight then differentiated with 150 nM phorbol 12-myristate 13-acetate (PMA) over 72 hours. Macrophages were then treated with compounds resuspended in RPMI media (1-50 µM) for 24 hours. Media was replaced with thiazolyl blue tetrazolium bromide solubilized in RPMI media (0.5 mg/mL) and incubated at 37°C for 2 hours. The reduced formazan crystals were solubilized with isopropyl alcohol and the absorbance of the resulting solution was measured at 560 nm with a background subtraction at 670 nm. The concentration required to reduce host cell viability by 50% (24hr LC_50_) value was measured from best fit curves plotting the relative absorbance values against drug concentration. This process was repeated with a 72-hr incubation and extended concentration range (1 – 300 µM) for select compounds.

### Image-based Evaluation of Intracellular Anti-Leishmanial Activity

Bone marrow derived macrophages (BMDMs) were isolated from BALB/c mice and cultured as previously described [18, 19]. Briefly, bone marrow was harvested from long bones of mice and the isolated cells were seeded at a concentration of 2 x 10^6^ cells/mL in petri dishes with RPMI media supplemented with 10% FBS, 1% Penn-Strep and 10% L929 conditioned media (LCM). Completely differentiated BMDMs are obtained after 7 days of culture with media change every 2 days. Harvested cells were seeded onto 10 mm glass cover slips at 5 x 10^5^ cells/ well of a 24 well plate in DMEM media (without LCM) and allowed to adhere overnight. BMDM cells are then infected overnight with LV82 *L. donovani* or *L. mexicana* (NR-51210) at a multiplicity of infection of 10. After infection, cells were washed three times with fresh media to remove extracellular promastigotes and treated with compounds for a 72-hour incubation. BMDM cells were then washed with phosphate buffered saline, fixed with ice-cold methanol, and stained with Giemsa (5% v/v in water). The cover slips with stained cells are mounted onto glass slides and imaged on EVOS XL (100X, Thermo Fisher Scientific). *Leishmania* amastigotes per 100 macrophages were determined in a blinded manner. The concentration required to reduce intracellular amastigote viability by 50% (IC_50_) value was measured from best fit curves plotting the normalized values against drug concentration.

### Effect of Compounds on Promastigote Viability

The effect of compounds on extracellular promastigotes was evaluated using a resazurin-based assay as described previously [10]. In short, late log phase *Leishmania* promastigotes were seeded in a 96 well plate at 1 x 10^5^ parasites per well and treated with drugs (0.5 – 200 µM) for 72 hours at 25°C. Ten microliters of resazurin (0.02% w/v) were added to the treated parasites to achieve a final concentration of 0.002% w/v and incubated for 24 hours. Viability of the promastigotes was assessed by fluorescence (excitation 544 nm, emission 590 nm, SpectraMax M2, Molecular Devices). The minimum inhibitory concentration required to reduce promastigote viability by 50% (MIC_50_) value was determined from best fit curves plotting the relative fluorescence values against drug concentration.

### Proteomic Analysis via Affinity Capture

To generate cell lysate, ten million THP-1 cells were plated overnight in a 10cm dish, then differentiated with 150 nM phorbol 12-myristate 13-acetate (PMA) for 72 hours. The resulting macrophages were infected with *L. mexicana* promastigotes at a multiplicity of infection of 1:10 and incubated for 18 hours. After infection, cells were washed three times with PBS to remove extracellular promastigotes. Cells were then collected by scraping and pelleted with centrifugation. The cell pellet was resuspended in cell buffer (50mM Tris-HCl, 150mM NaCl, 2mM EDTA, 0.5% triton-x-100) and subjected to five freeze-thaw cycles. Samples were centrifuged 14,000g x 10 minutes to remove cellular debris. Protein concentration of supernatant was determined by BCA assay and cell lysate was stored at –80°C until further use.

Functionalized agarose beads were made as follows. NHS activated agarose beads were washed three times to remove acetone and resuspended in PBS to create a slurry. 494 (197 with ligation chemical handle) was prepared in 0.3 mL at 25mM in DMSO and 1.7 mL PBS was added to 1 mL of agarose slurry for a final volume of 3 mL at a concentration of 2.5mM (10% v/v DMSO). Beads were incubated with 494 for 2 hrs at room temperature, then washed three times with PBS to remove unbound drug. Beads were then blocked with 1M Tris-HCl (pH 7.5) for 1 hr at room temperature, washed twice with buffer (100mM Tris-HCl, 300mM NaCl, 2mM EDTA, 0.5% NP-40) and resuspended in 1 mL of buffer. Control beads were generated by coupling 2.5mM butylamine to NHS-agarose beads in the same manner as above.

500 µg cell lysate was incubated with 200 µL slurry at a final volume of 1 mL for 1 hr. Unbound protein was removed through four 15-minute washes. Beads were then boiled in Laemli sample buffer and resolved in a polyacrylamide gel and stained with Coomassie. Gel lanes were then excised and destained. Samples were reduced with dithiothreitol, alkylated with iodoacetamide, then digested with trypsin overnight at 37°C. Peptides were extracted, acidified, desalted using C18 spin columns, and analyzed by LC-MS/MS using Thermo Easy nLC 1200-QExactive HF. Data was processed using Proteome Discoverer 2.5 against Uniprot human and *L. mexicana* databases. Only proteins with > 1 peptide were reported. Data was imported into Perseus for imputation and statistical analysis. Log2(fold-change) compared to butylamine bead was plotted vs –Log10(p-value) to identify outliers. Proteins hits were identified as those with Log2(fold-change) > 1 and p-value < 0.05.

### Proteomic Analysis via Thermal Proteome Profiling

To generate cell lysate, ten million THP-1 cells were plated overnight in a 10cm dish, then differentiated with 150 nM phorbol 12-myristate 13-acetate (PMA) for 72 hours. The resulting macrophages were infected with *L. mexicana* promastigotes at a multiplicity of infection of 10 and incubated for 18 hours. After infection, cells were washed three times with PBS to remove extracellular promastigotes. Cells were then collected by scraping and pelleted with centrifugation. The cell pellet was resuspended in phosphate buffered saline and subjected to five freeze-thaw cycles. Samples were centrifuged 14,000g x 10 minutes to remove cellular debris. Protein concentration of supernatant was determined by BCA assay and cell lysate was stored at –80°C until further use.

The cell lysate was divided into two portions. One portion was spiked with the 197 ligand in DMSO to generate the with ligand sample; and the other portion was spiked with DMSO to generate the without ligand sample. The final concentration of 147 in the with ligand sample was 100 µM, and both the without and with ligand samples contained 1% DMSO. The with and without samples were each subjected five replicate one-pot TPP analyses like that previously described.[20–22] In each replicate aliquots of the with and without samples were distributed into a series of 12 different samples before heating for 3 min at a temperature gradient ranging from 43– 65 °C with 2 °C intervals. After heat treatment, the samples are equilibrated at room temperature for 3 min before placing on ice. The with samples and the without samples in each biological replicate were combined to generate a single with and without ligand sample, respectively. The combined samples were centrifuged at 48000 rpm for 20 min using a TPA100.1 rotor and a Beckman Optima TL ultracentrifuge. The supernatants were transferred into 10 kDa MWCO centrifugal filter units and buffer exchanged to 8 M urea in 0.1 M Tris-HCl pH 8.5 before TCEP reduction and MMTS alkylation. The alkylated proteins were then digested with trypsin, and the peptides generated from the without and with samples from each of the five replicates were labeled with a TMT 10-Plex according to the manufacturer’s protocol. Ultimately, a C18 Macrospin column cleanup was performed on the combined TMT 10-plex sample, sample prior to LC-MS/MS analysis.

The LC-MS/MS analyses were performed using a nanoAcquity UPLC system (Waters) coupled to a Thermo Orbitrap Fusion Lumos mass spectrometer system. The dried peptide material generated from TPP analysis was reconstituted in 15 μL of 1% TFA, 2% acetonitrile in H2O, and a 1 μl aliquot was injected into the system. The peptides were first trapped on a Symmetry C18 20 mm×180 μm trapping column (5 μL/min at 99.9/0.1 water/acetonitrile, v/v). The analytical separation was performed using an Acquity 75 μm × 250 mm high strength silica (HSS) T3 C18 column with a 1.8 μm particle size (Waters); the column temperature was set to 55 °C. Peptide elution was performed using a 90 min linear gradient of 3–30 % ACN with 0.1 % formic acid at a flow rate of 400 nL/min. The MS data were collected using a top 20 data-dependent acquisition method which included MS1 at 120k and MS2 at 50k resolution. The MS1 AGC target was 4.0 × 105 ions with a max injection time of 50 ms. For MS2, the AGC target was 1.0 × 105 ions with a max injection time of 105 ms. The collision energy was set to 38 %, and the scan range was 375– 1500 m/z. The isolation window was 0.7 and the dynamic exclusion duration was 60 s. The peptide sample was subjected to three LC-MS/MS analyses.

Proteome Discoverer 2.2 (Thermo) was used to search the raw LC-MS/MS data against the mouse and *Leishmania* proteins in the 2017-06-07 release of the UniProt Knowledgebase. The raw LC MS/MS data were searched using fixed MMTS modification on cysteine; TMT 10-Plex labeling of lysine side chains and peptide N-termini; variable oxidation of methionine; variable deamidation of asparagine and glutamine; and variable acetylation of the protein N-terminus. Trypsin was set as the enzyme, and up to two missed cleavages were allowed. For peptide and protein quantification, reporter ion abundance was set as intensity, and the normalization mode and scaling mode were each set as none. All other settings were left as the default values. Only proteins/peptides with protein/peptide FDR confidence labeled as “high” (i.e., FDR < 0.01) and with no quantification channels being 0 were used for subsequent analyses. For each biological replicate, a normalization factor was calculated by dividing the ratio of the summed signal intensities recorded in the samples from each biological replicate by the summed signal intensities in the 126 TMT channel. For each identified protein, a ratio of the observed reporter ion intensities in the with sample to the without sample was generated for each biological replicate. The resulting ratio was divided by the normalization factor for each of the replicates. These normalized ratios (fold-change) were then *log*2-base transformed, averaged, and tested by a two-tailed student’s t-test comparing with a mean of 0. Proteins hits were defined as those with a |z-score| > 1 and p-value < 0.05.

### 197 Target Validation

B6.129P2-*Lyz2^tm1(cre)Ifo^*/J mice[23] (stock #004781, herein termed ‘Lys K/O’) were obtained from Jackson Laboratory. This strain has a nuclear-localized Cre recombinase inserted into the first coding ATG of the lysozyme 2 gene (Lyz2) which eliminates endogenous Lyz2 gene function. Wildtype C57BL6/J mice (stock #000664) were obtained as a control.

Bone marrow derived macrophages (BMDMs) were isolated from both mice and cultured as described above in “Image-based Evaluation of Intracellular Anti-Leishmanial Activity”. BMDMs were seeded onto glass cover slips, infected with LV82 *L. donovani* at a multiplicity of infection of 1:10. Cells were washed to remove extracellular promastigotes and treated with compounds (197, amphotericin B) for a 72-hour incubation. BMDM cells were then washed with phosphate buffered saline, fixed with ice-cold methanol, and stained with Giemsa (5% v/v in water). The cover slips with stained cells are mounted onto glass slides and imaged on EVOS XL (100X, Thermo Fisher Scientific). Leishmania amastigotes per 100 macrophages were determined in a blinded manner. The concentration required to reduce intracellular amastigote viability by 50% (IC_50_) value was measured from best fit curves plotting the normalized values against drug concentration.

## ACKNOWLEDGMENTS

The authors would like to thank Dr. Abhay Satoskar for providing the *Leishmania* samples essential for our research. This work was supported by grant R01AI125147. Also, we thank the UNC Metabolomics & Proteomics core for their contribution to the work.

## CONFLICT OF INTEREST

The authors declare that they have no conflicts of interest.

## AUTHOR CONTRIBUTIONS

Initial Manuscript writing was Gurysh, Zahid, Ainslie. Parasite and cell assays were performed by Gurysh, Zahid, Johnson, Varma, Woodring and Vath. Cell viability assays were performed in part by Pino. Figure generation and editing were performed by Hendricksen and Gurysh. Landavazo, Namjoshi, Wilson and Blough performed chemical synthesis and characterization. Quan and Fitzgerald performed thermal proteomic analysis. Bachelder, Ainslie, Gurysh, Johnson and Zahid contributed to experimental design. Supervision was performed by Bachelder, Blough and Ainslie. Funding was provided by Blough and Ainslie.

## SUPPLEMENTAL INFORMATION CAPTIONS

**Supplemental Table 1.**
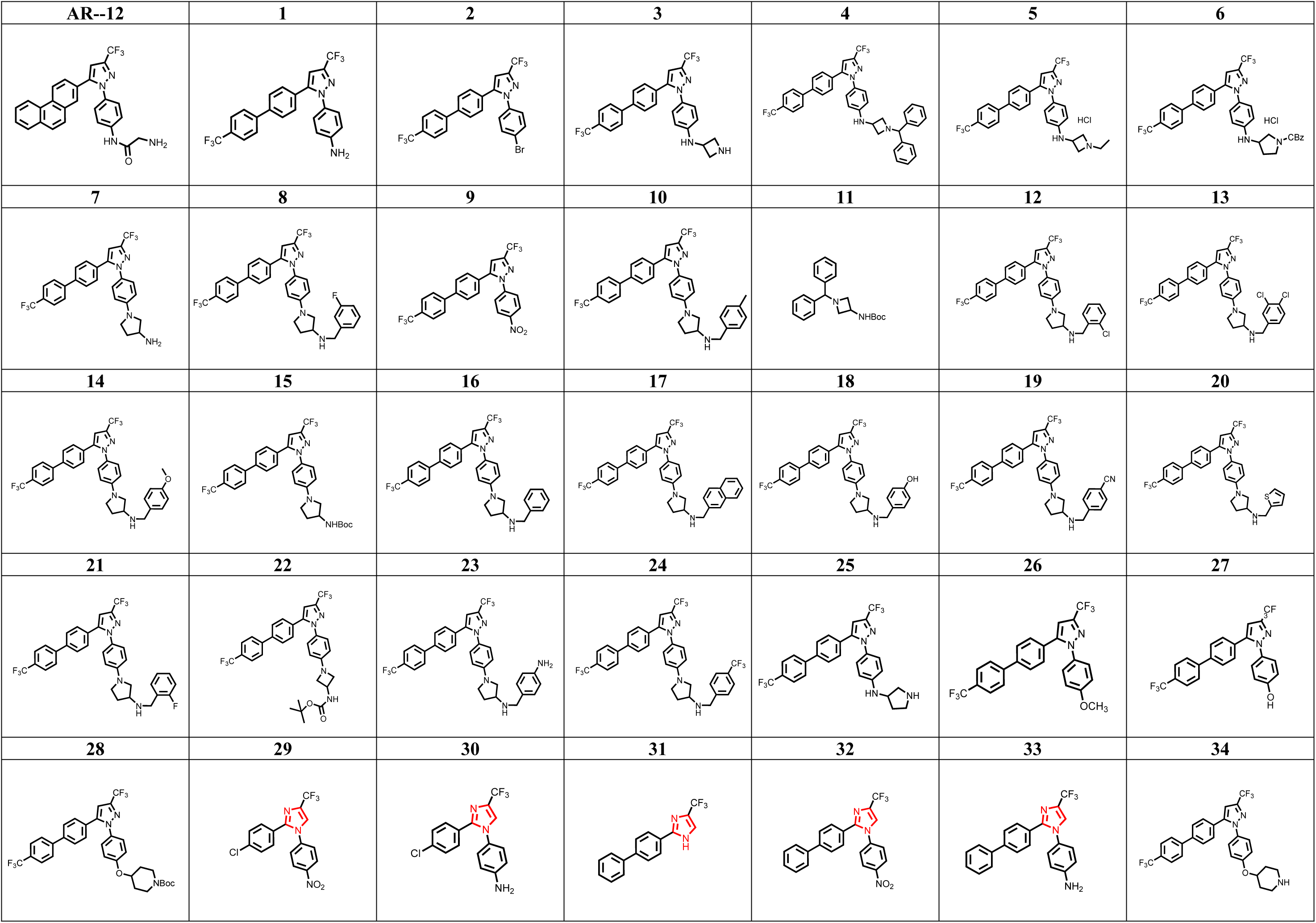
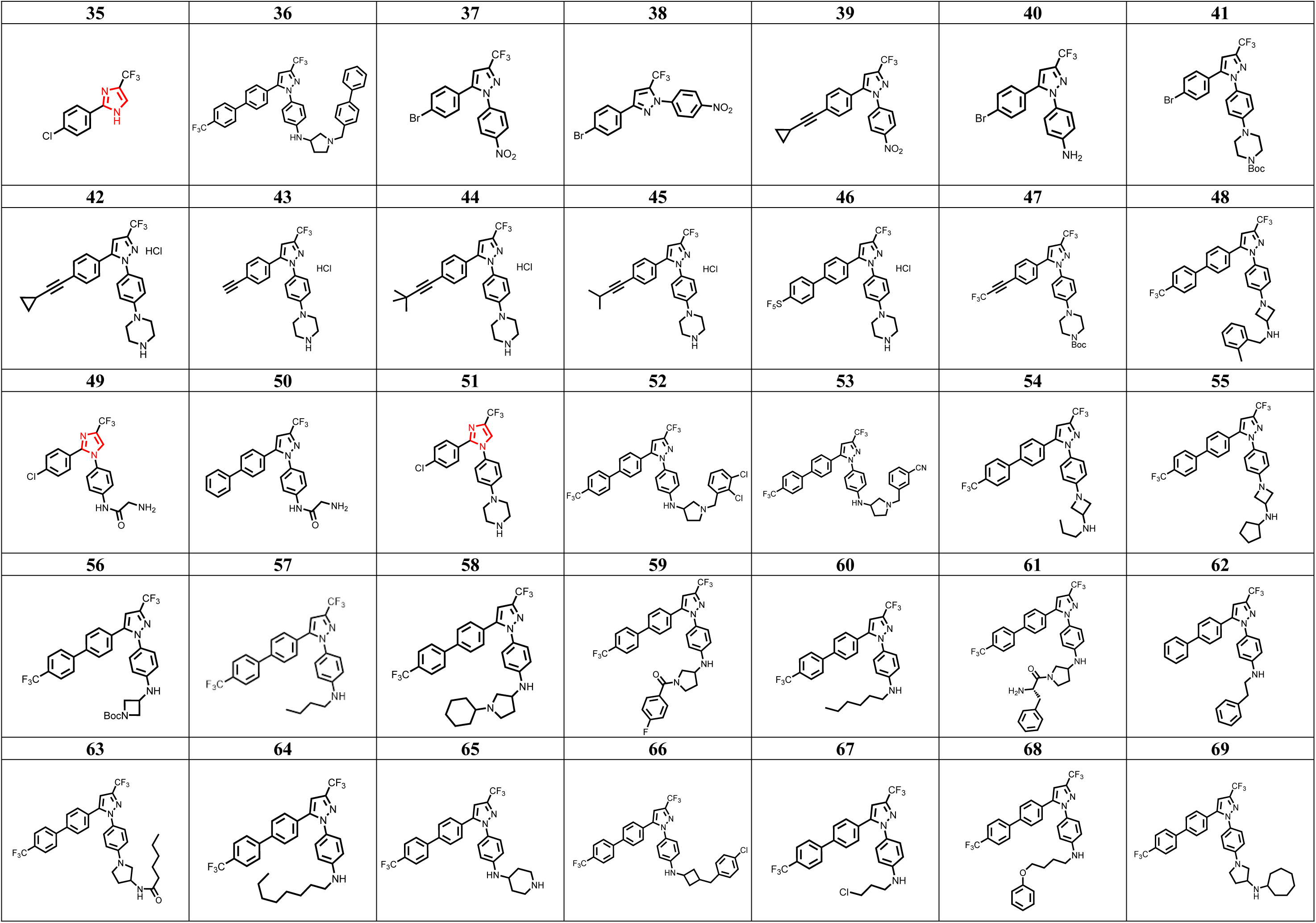
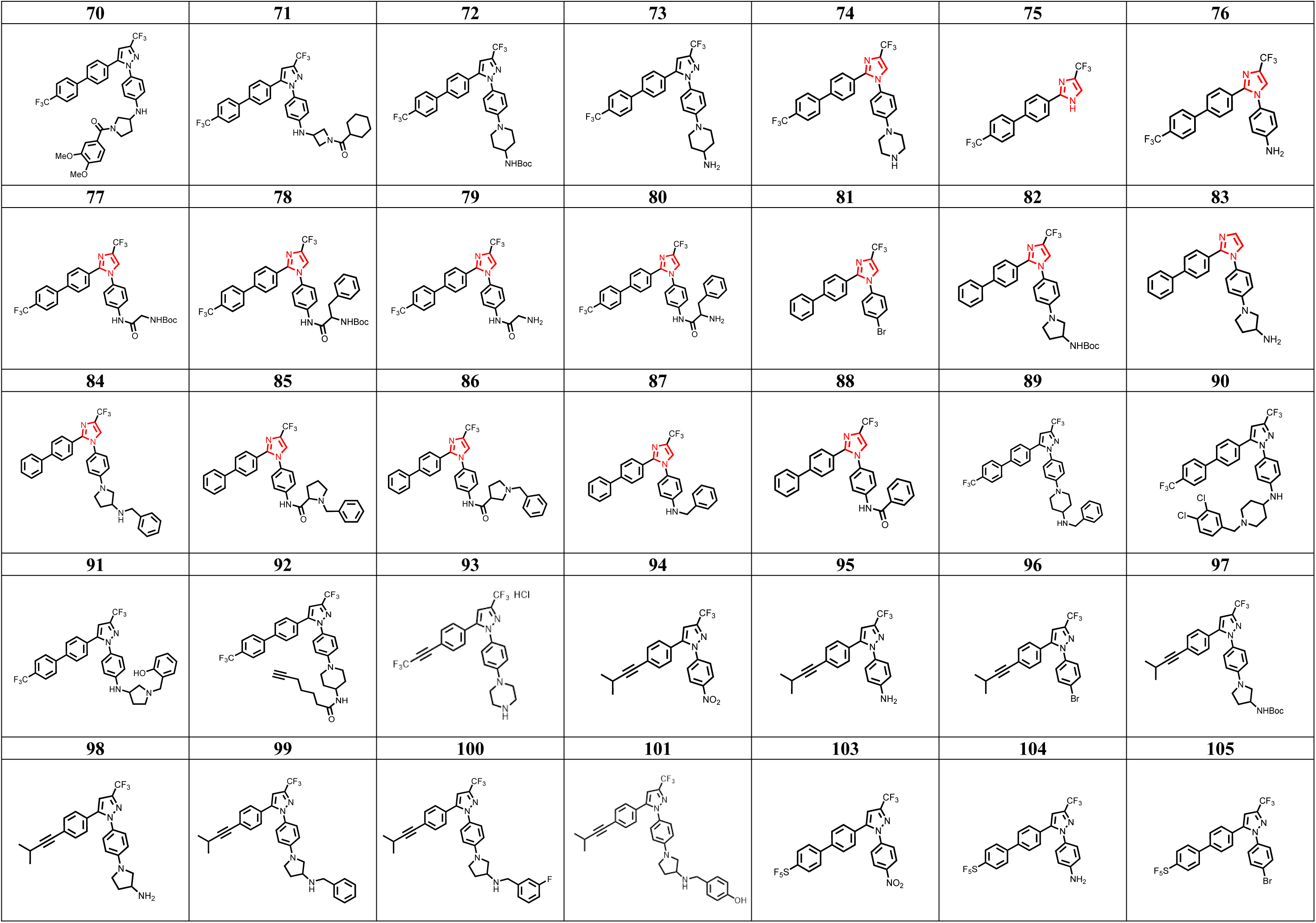
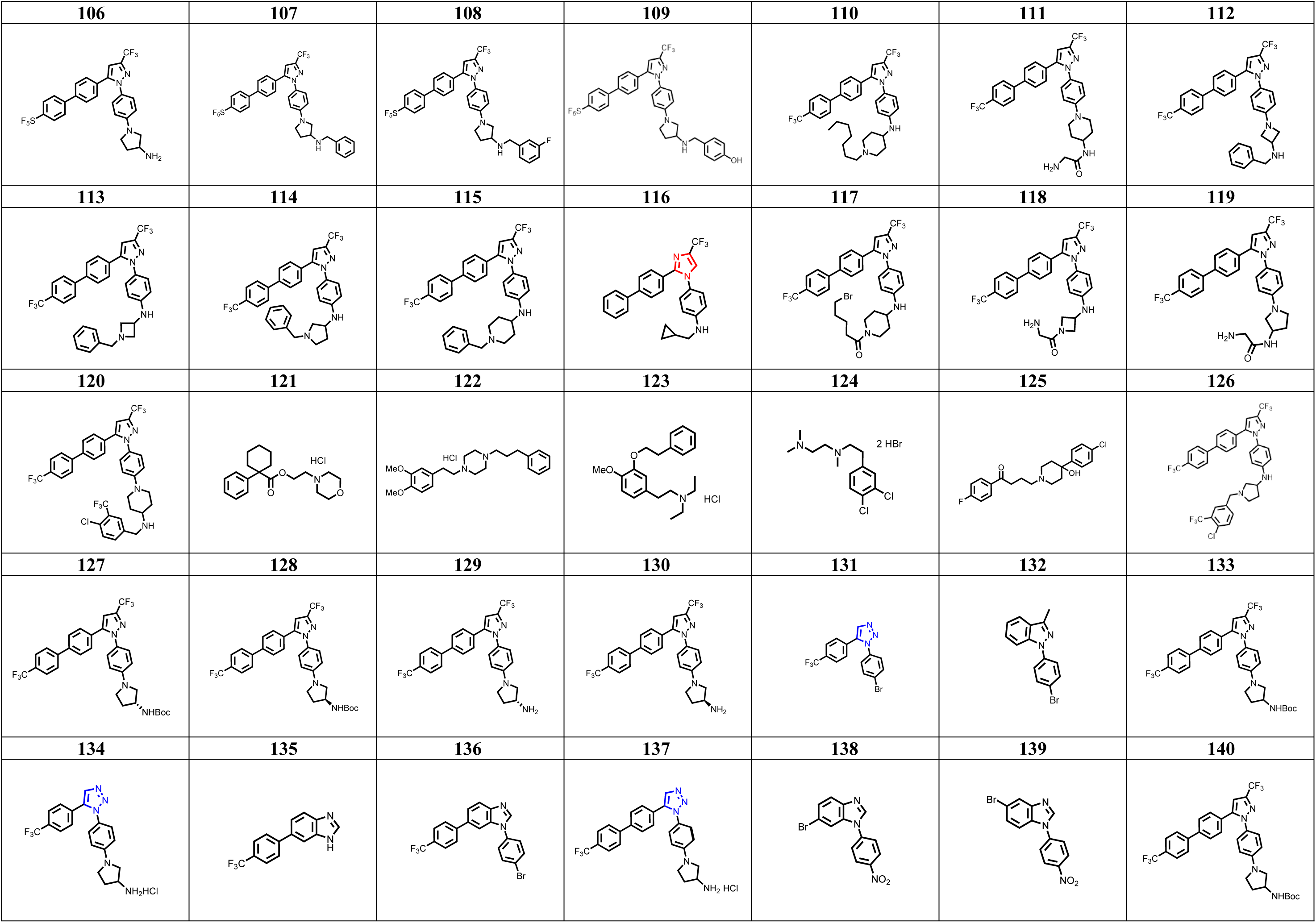
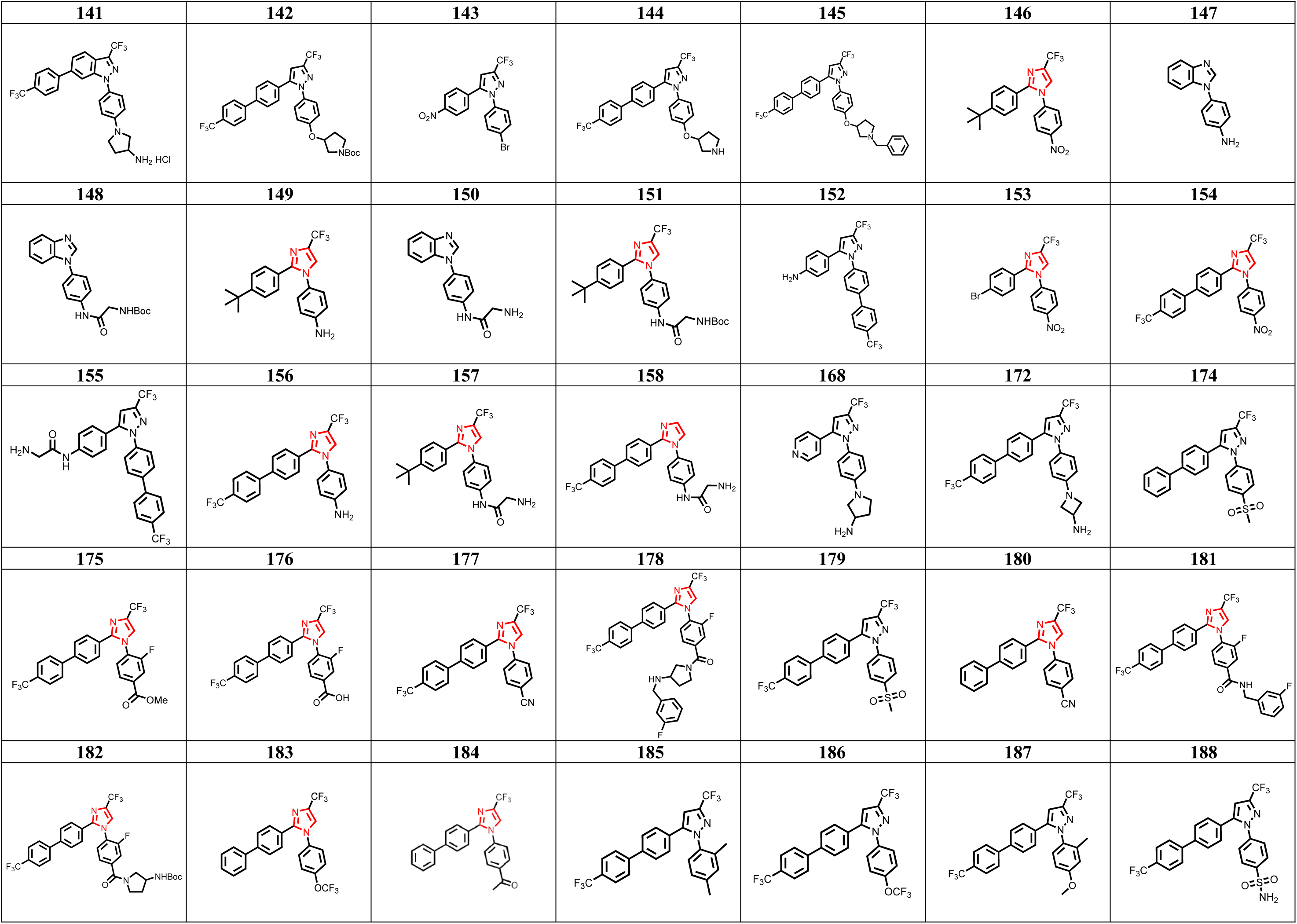
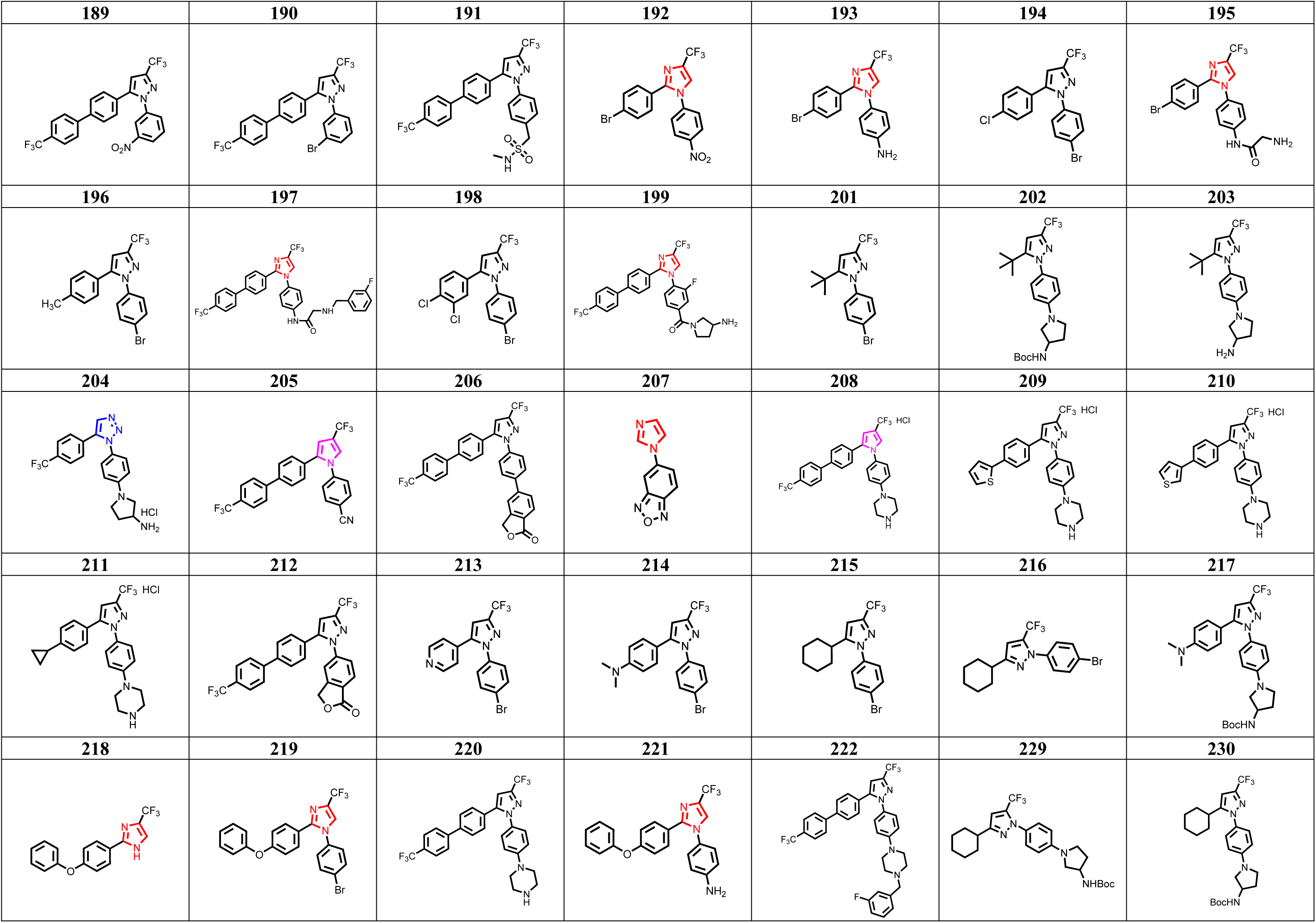
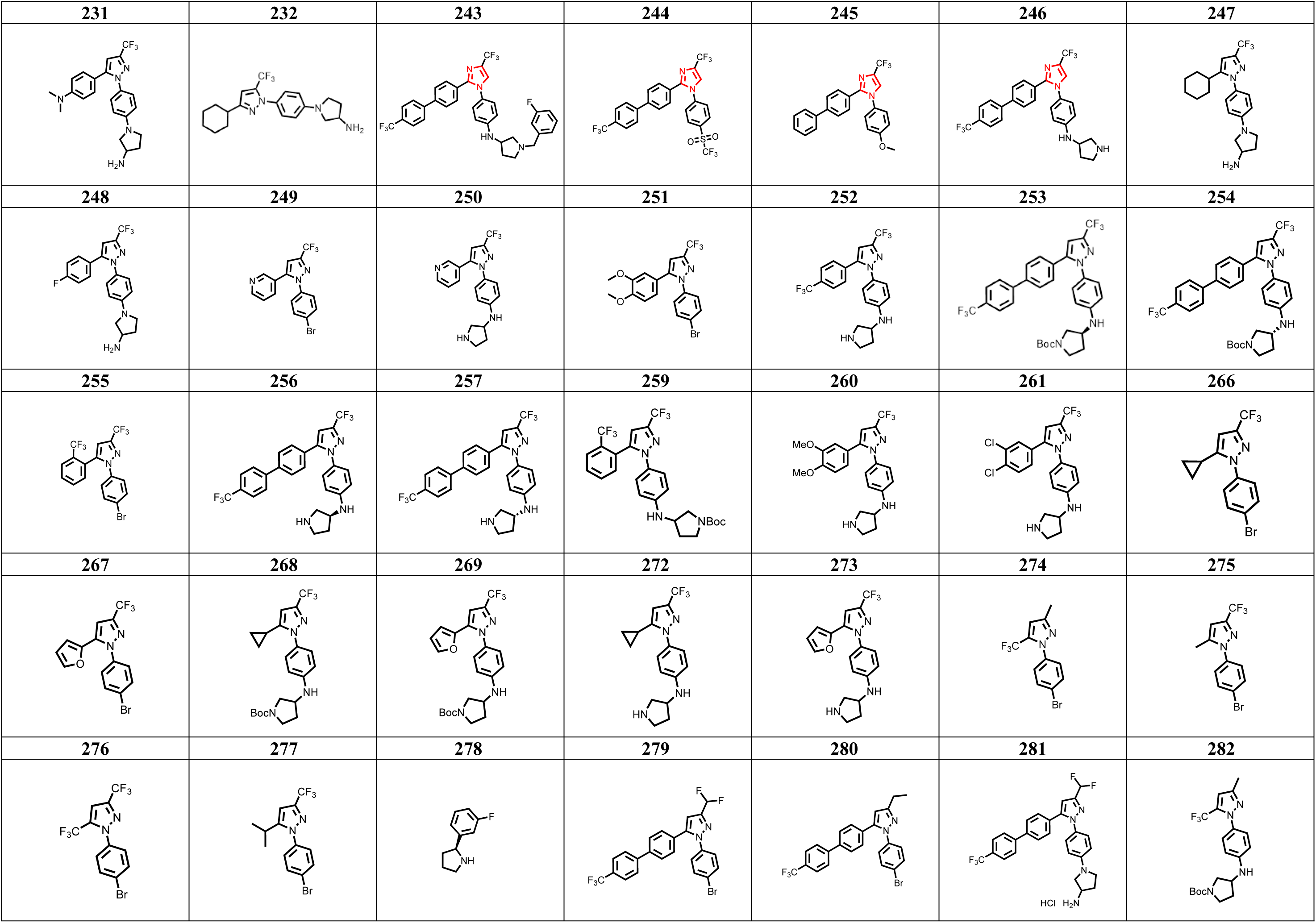
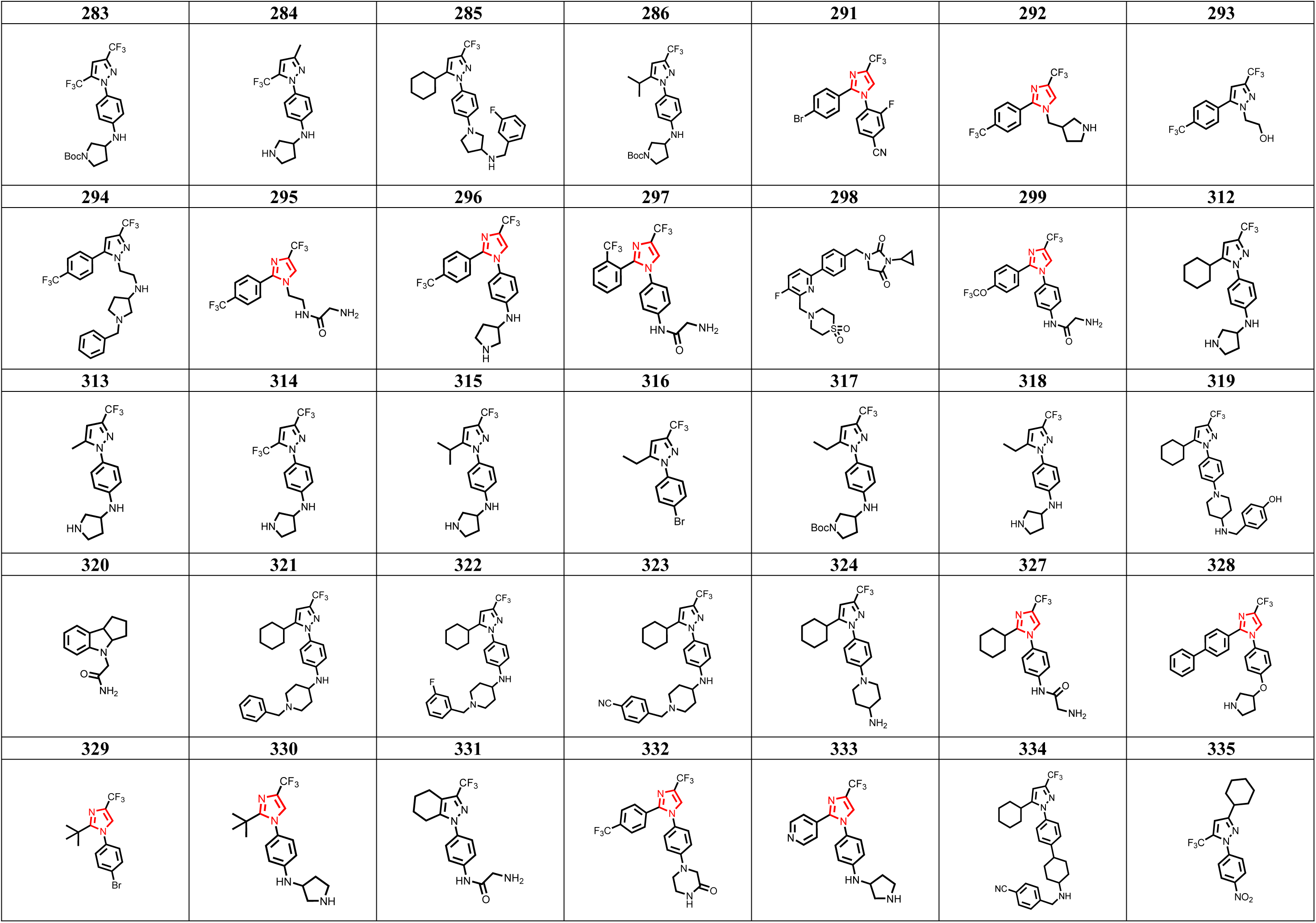
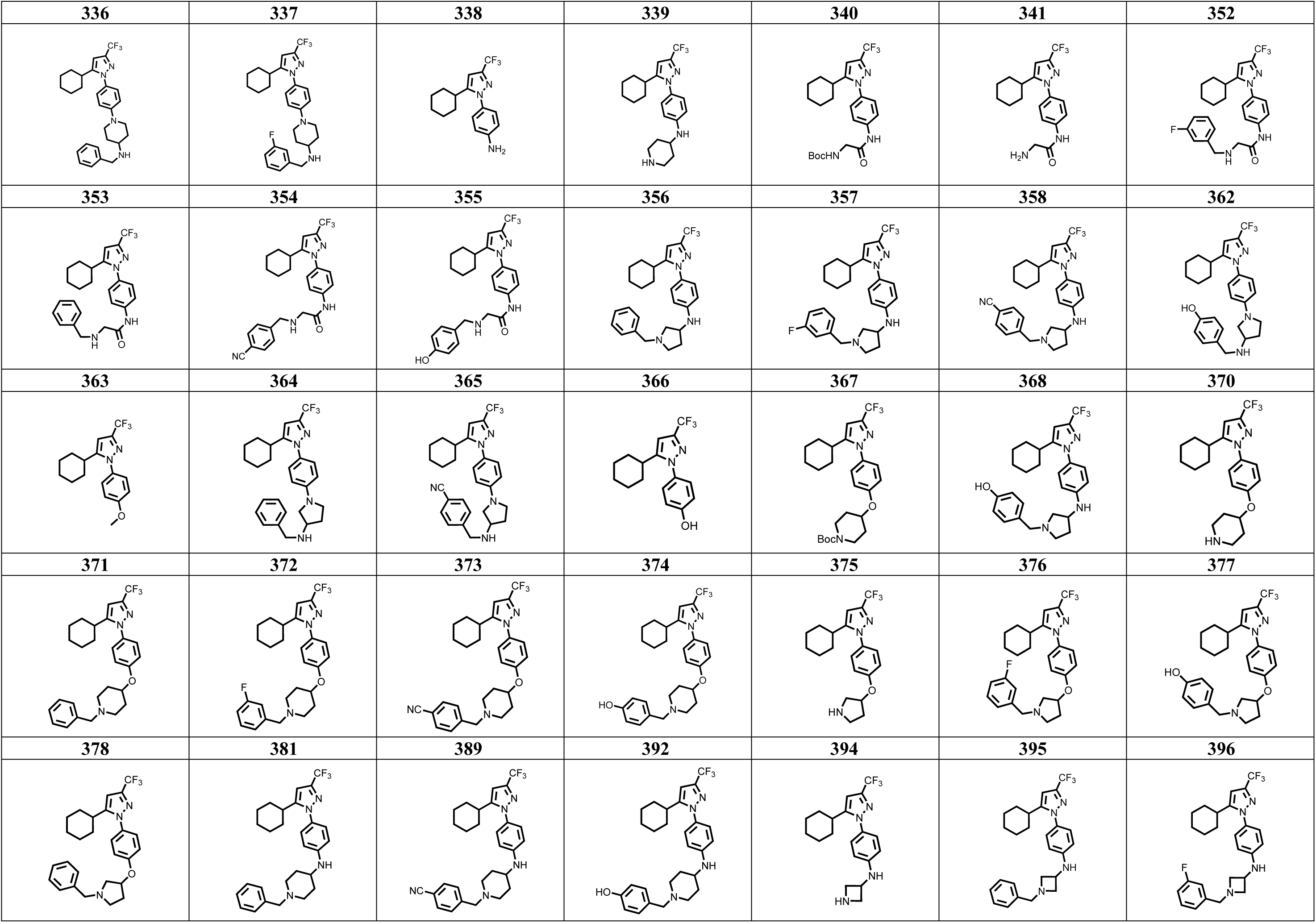
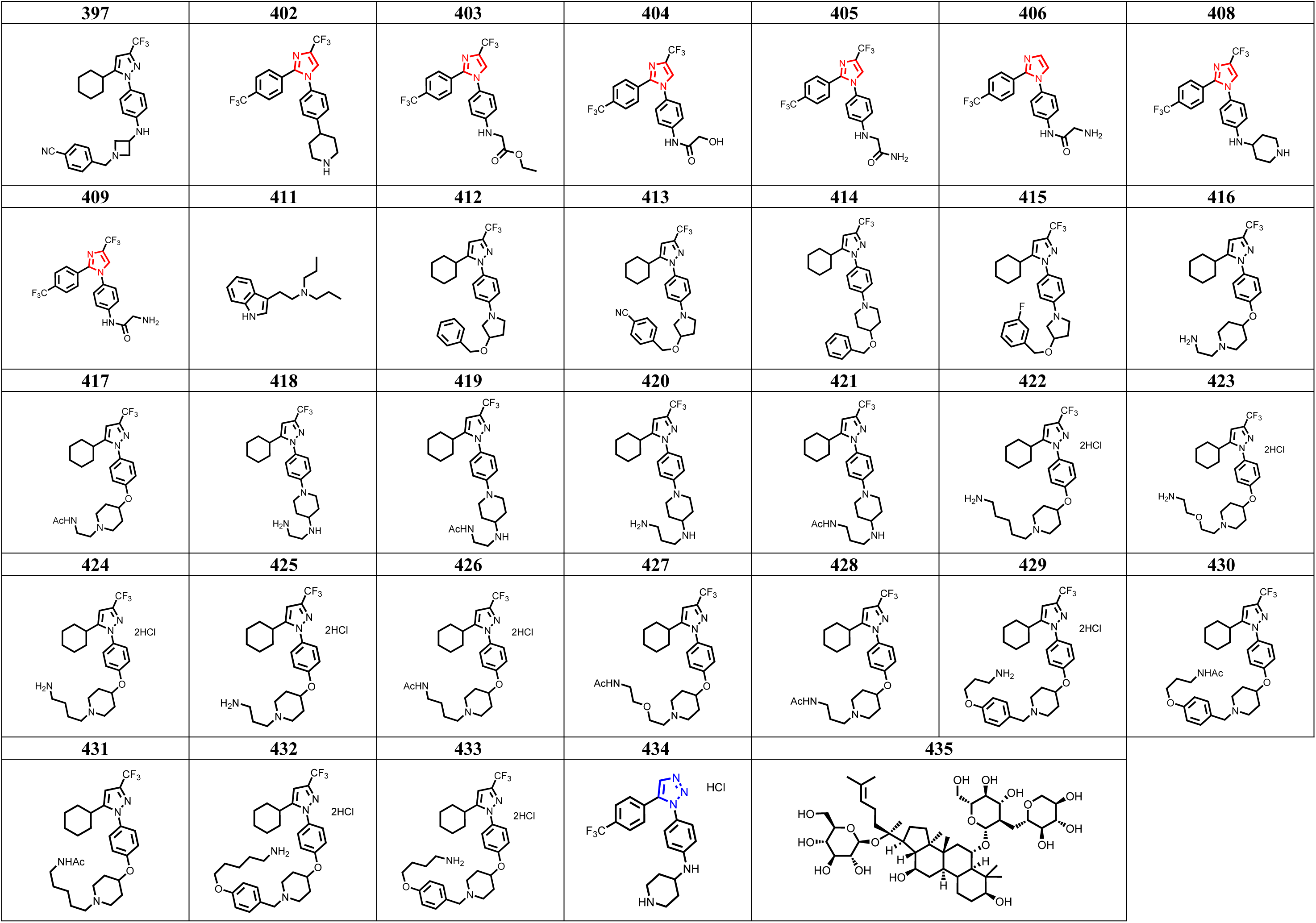
Chemical Structures of all compounds screened.

**Supplemental Figure 1.**
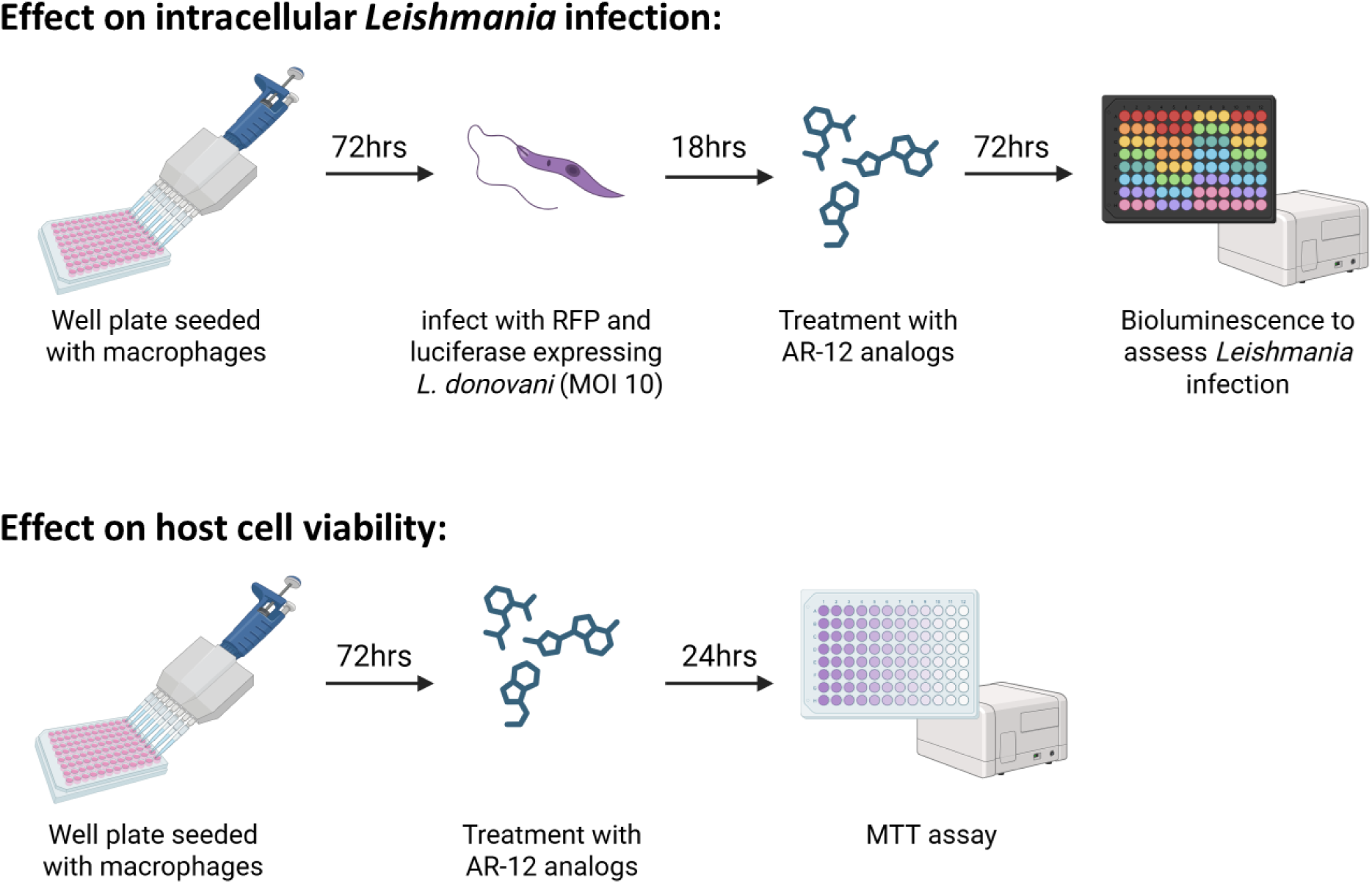
Initial Screening Approach. Schematic illustrating the medium-throughput luminescence-based assay to determine effect of compounds on intracellular *Leishmania* infection and parallel screening to determine effect of compounds on host cell viability.

**Supplemental Table 2.** Results of primary screen in all compounds. Concentration at which intracellular *Leishmania donovani* burden is reduced by 50% in THP1 macrophages (Lum IC_50_) as identified by luminescence assay. Concentration where THP1 macrophage cell viability is 50% (LC_50_) after 24-hour incubation with compound as determined by MTT assay. Selectivity between host-directed effect and cytotoxicity, defined as 24h LC_50_ / Lum IC_50_. Parental compound AR-12 provided for reference. Compounds highlighted in grey have higher selectivity than parental compound AR-12. Compounds highlighted in yellow were selected for secondary screening. ND = not determined.

| CMPD | 24h LC<sub>50</sub><br>(μM) | Lum<br>IC<sub>50</sub><br>(μM) | Selectivity |
| --- | --- | --- | --- |
| AR-12 | 13.3 | 3.4 | 3.9 |
| 1 | 19.1 | >10 | <1.9 |
| 2 | >50 | >10 | ND |
| 3 | 11.0 | 1.8 | 6.1 |
| 4 | >50 | >10 | ND |
| 5 | 13.3 | 2.3 | 5.8 |
| 6 | >50 | >10 | ND |
| 7 | 5.2 | 2.4 | 2.1 |
| 8 | >50 | >10 | ND |
| 9 | >50 | >10 | ND |
| 10 | >50 | >10 | ND |
| 11 | 1.4 | >10 | <0.1 |
| 12 | >50 | >10 | ND |
| 13 | >50 | >10 | ND |
| 14 | 50.0 | >10 | <5.0 |
| 15 | >50 | >10 | ND |
| 16 | >50 | >10 | ND |
| 17 | >50 | >10 | ND |
| 18 | 24.5 | 4.0 | 6.1 |
| 19 | >50 | >10 | ND |
| 20 | >50 | >10 | ND |
| 21 | >50 | >10 | ND |
| 22 | >50 | - | - |
| 23 | 5.8 | 0.4 | 14.8 |
| 24 | >50 | >10 | ND |
| 25 | 28.9 | 0.7 | 43.8 |
| 26 | 5.8 | >10 | <0.6 |
| 27 | >50 | >10 | ND |
| 28 | >50 | >10 | ND |
| 29 | >50 | >10 | ND |
| 30 | >50 | >10 | ND |
| 31 | >50 | >10 | ND |
| 32 | 6.3 | >10 | <0.6 |
| 33 | 1.5 | >10 | <0.2 |
| 34 | >50 | 6.0 | >8.3 |
| 35 | >50 | >10 | ND |
| 36 | >50 | >10 | ND |
| 37 | >50 | >10 | ND |
| 38 | >50 | >10 | ND |
| 39 | >50 | >10 | ND |
| 40 | >50 | >10 | ND |
| 41 | >50 | >10 | ND |
| 42 | 0.7 | 4.2 | 0.2 |
| 43 | 0.5 | 4.8 | 0.1 |
| 44 | 7.4 | 0.9 | 8.1 |
| 45 | 3.2 | 3.7 | 0.9 |
| 46 | 10.0 | >10 | <1.0 |
| 47 | >50 | >10 | ND |
| 48 | >50 | >10 | ND |
| 49 | 48.8 | >10 | <4.8 |
| 50 | 23.5 | 1.5 | 16.0 |
| 51 | 25.7 | 8.2 | 3.1 |
| 52 | >50 | >10 | ND |
| 53 | >50 | 0.4 | >112.6 |
| 54 | 9.9 | 2.4 | 4.1 |
| 55 | 29.7 | 8.5 | 3.5 |
| 56 | >50 | >10 | ND |
| 57 | >50 | >10 | ND |
| 58 | 2.6 | 2.2 | 1.2 |
| 59 | >50 | >10 | ND |
| 60 | >50 | >10 | ND |
| 61 | 6.1 | 2.3 | 2.7 |
| 62 | >50 | >10 | ND |
| 63 | >50 | >10 | ND |
| 64 | >50 | >10 | ND |
| 65 | 4.0 | 2.3 | 1.7 |
| 66 | >50 | >10 | ND |
| 67 | >50 | >10 | ND |
| 68 | 34.7 | >10 | <3.5 |
| 69 | 12.4 | 5.5 | 2.3 |
| 70 | >50 | >10 | ND |
| 71 | 8.4 | >10 | <0.8 |
| 72 | >50 | >10 | ND |
| 73 | 4.8 | 2.7 | 1.8 |
| 74 | 6.5 | 2.8 | 2.3 |
| 75 | 48.8 | >10 | <4.9 |
| 76 | 33.2 | >10 | <3.3 |
| 77 | >50 | >10 | ND |
| 78 | >50 | >10 | ND |
| 79 | 6.3 | 7.1 | 0.9 |
| 80 | 8.2 | 3.0 | 2.7 |
| 81 | >50 | >10 | ND |
| 82 | >50 | >10 | ND |
| 83 | 6.2 | 1.8 | 3.5 |
| 84 | >50 | 6.5 | >7.7 |
| 85 | >50 | >10 | ND |
| 86 | 21.9 | 1.4 | 16.0 |
| 87 | >50 | 6.7 | >7.4 |
| 88 | >50 | >10 | ND |
| 89 | 8.0 | 4.0 | 2.0 |
| 90 | >50 | >10 | ND |
| 91 | >50 | 1.4 | >36.1 |
| 92 | 14.7 | >10 | <1.5 |
| 93 | 40.0 | >10 | <4.0 |
| 94 | 29.0 | >10 | <2.9 |
| 95 | >50 | >10 | ND |
| 96 | 37.5 | >10 | <3.8 |
| 97 | >50 | >10 | ND |
| 98 | 19.5 | >10 | <1.9 |
| 99 | 29.2 | >10 | <2.9 |
| 100 | >50 | >10 | ND |
| 101 | 0.5 | 1.1 | 0.4 |
| 102 | - | - | - |
| 103 | >50 | >10 | ND |
| 104 | >50 | >10 | ND |
| 105 | >50 | >10 | ND |
| 106 | 6.0 | 8.1 | 0.7 |
| 107 | >50 | >10 | ND |
| 108 | >50 | >10 | ND |
| 109 | 9.4 | 3.3 | 2.8 |
| 110 | 8.5 | 4.0 | 2.1 |
| 111 | 5.2 | 2.9 | 1.8 |
| 112 | 6.3 | >10 | <0.6 |
| 113 | 19.0 | >10 | <1.9 |
| 114 | 13.6 | >10 | <1.4 |
| 115 | 13.2 | 7.4 | 1.8 |
| 116 | >50 | 9.9 | >5.0 |
| 117 | >50 | 6.1 | >8.2 |
| 118 | 6.2 | 2.6 | 2.4 |
| 119 | 37.2 | >10 | <3.7 |
| 120 | >50 | >10 | ND |
| 126 | >50 | >10 | ND |
| 127 | >50 | >10 | ND |
| 128 | >50 | >10 | ND |
| 129 | 6.3 | 0.1 | 46.7 |
| 130 | 4.2 | 0.2 | 23.9 |
| 131 | 37.2 | >10 | <3.7 |
| 132 | >50 | >10 | ND |
| 133 | 29.0 | 0.9 | 32.5 |
| 134 | 22.6 | 0.5 | 42.1 |
| 135 | >50 | >10 | ND |
| 136 | >50 | >10 | ND |
| 137 | 7.3 | 4.4 | 1.7 |
| 138 | >50 | >10 | ND |
| 139 | >50 | >10 | ND |
| 140 | >50 | >10 | ND |
| 141 | 7.7 | 3.6 | 2.1 |
| 142 | >50 | >10 | ND |
| 143 | >50 | >10 | ND |
| 144 | 5.4 | >10 | <0.5 |
| 145 | >50 | >10 | ND |
| 146 | >50 | 4.4 | >11.4 |
| 147 | >50 | >10 | ND |
| 148 | >50 | >10 | ND |
| 149 | 28.8 | 9.1 | 3.2 |
| 150 | >50 | >10 | ND |
| 151 | 31.8 | >10 | <3.2 |
| 152 | 43.6 | >10 | <4.4 |
| 153 | >50 | >10 | ND |
| 154 | 25.5 | 2.9 | 8.9 |
| 155 | 6.1 | 4.0 | 1.5 |
| 156 | 20.8 | 5.8 | 3.6 |
| 157 | 27.6 | >10 | <2.8 |
| 158 | 16.0 | 0.2 | 81.0 |
| 168 | >50 | 10.0 | >5.0 |
| 172 | >50 | >10 | ND |
| 174 | >50 | >10 | ND |
| 175 | >50 | >10 | ND |
| 176 | >50 | >10 | ND |
| 177 | >50 | >10 | ND |
| 178 | >50 | >10 | ND |
| 179 | 45.0 | >10 | <4.5 |
| 180 | >50 | >10 | ND |
| 181 | 28.7 | >10 | <2.9 |
| 182 | 18.0 | 5.9 | 3.0 |
| 183 | >50 | >10 | ND |
| 184 | >50 | 5.0 | >10 |
| 185 | >50 | >10 | ND |
| 186 | >50 | >10 | ND |
| 187 | >50 | >10 | ND |
| 188 | 9.0 | 3.1 | 2.9 |
| 189 | >50 | >10 | ND |
| 190 | >50 | >10 | ND |
| 191 | 12.0 | >10 | <1.2 |
| 192 | >50 | >10 | ND |
| 193 | >50 | >10 | ND |
| 194 | >50 | >10 | ND |
| 195 | 38.6 | >10 | <3.9 |
| 196 | >50 | >10 | ND |
| 197 | >50 | 2.4 | >20.6 |
| 198 | >50 | >10 | ND |
| 199 | 9.4 | 1.3 | 7.2 |
| 201 | >50 | >10 | ND |
| 202 | >50 | 8.5 | >5.9 |
| 203 | 11.8 | 2.9 | 4.1 |
| 204 | 37.2 | >10 | <3.7 |
| 205 | >50 | >10 | ND |
| 206 | >50 | >10 | ND |
| 207 | >50 | >10 | ND |
| 208 | 31.0 | 5.7 | 5.4 |
| 209 | 10.2 | 3.0 | 3.4 |
| 210 | 10.5 | 2.5 | 4.3 |
| 211 | 13.2 | 2.1 | 6.3 |
| 212 | >50 | >10 | ND |
| 213 | >50 | >10 | ND |
| 214 | >50 | >10 | ND |
| 215 | >50 | >10 | ND |
| 216 | >50 | >10 | ND |
| 217 | >50 | >10 | ND |
| 218 | >50 | >10 | ND |
| 219 | >50 | >10 | ND |
| 220 | 15.4 | 1.3 | 11.8 |
| 221 | >50 | >10 | ND |
| 222 | >50 | >10 | ND |
| 229 | 5.0 | 3.4 | 1.5 |
| 230 | 9.5 | >10 | <1.0 |
| 231 | 10.8 | 1.8 | 6.2 |
| 232 | 14.4 | 2.6 | 5.6 |
| 243 | >50 | >10 | ND |
| 244 | >50 | >10 | ND |
| 245 | >50 | >10 | ND |
| 246 | 5.5 | 3.8 | 1.4 |
| 247 | 7.6 | 3.3 | 2.3 |
| 248 | 14.4 | 2.6 | 5.6 |
| 249 | 50.0 | >10 | <5.0 |
| 250 | >50 | >10 | ND |
| 251 | >50 | >10 | ND |
| 252 | 14.4 | 2.3 | 6.2 |
| 253 | >50 | >10 | ND |
| 254 | 50.0 | >10 | <5.0 |
| 255 | >50 | >10 | ND |
| 256 | 6.2 | 2.5 | 2.5 |
| 257 | 5.0 | 2.5 | 2.0 |
| 259 | 23.6 | >10 | <2.4 |
| 260 | 13.9 | 4.3 | 3.2 |
| 261 | 22.4 | 3.1 | 7.3 |
| 266 | >50 | >10 | ND |
| 267 | >50 | >10 | ND |
| 268 | >50 | >10 | ND |
| 269 | >50 | >10 | ND |
| 272 | 17.2 | 5.6 | 3.1 |
| 273 | 15.6 | 4.9 | 3.2 |
| 274 | >50 | >10 | ND |
| 275 | >50 | >10 | ND |
| 276 | 32.2 | >10 | <0.6 |
| 277 | >50 | >10 | ND |
| 278 | >50 | >10 | ND |
| 279 | >50 | >10 | ND |
| 280 | >50 | >10 | ND |
| 281 | 14.3 | 2.0 | 7.2 |
| 282 | >50 | >10 | ND |
| 283 | >50 | >10 | ND |
| 284 | >50 | >10 | ND |
| 285 | >50 | >10 | ND |
| 286 | >50 | >10 | ND |
| 291 | >50 | >10 | ND |
| 292 | >50 | >10 | ND |
| 293 | >50 | >10 | ND |
| 294 | 28.2 | >10 | <2.8 |
| 295 | >50 | >10 | ND |
| 296 | 12.9 | 3.0 | 4.3 |
| 297 | >50 | >10 | ND |
| 298 | >50 | >10 | ND |
| 299 | 31.6 | >10 | <3.2 |
| 312 | 25.5 | 4.8 | 5.3 |
| 313 | >50 | >10 | ND |
| 314 | 21.9 | 2.7 | 8.0 |
| 315 | 25.1 | 3.3 | 7.6 |
| 316 | >50 | >10 | ND |
| 317 | >50 | >10 | ND |
| 318 | 27.0 | 1.8 | 14.8 |
| 319 | 6.4 | 0.8 | 8.1 |
| 320 | >50 | >10 | ND |
| 321 | 23.9 | 5.1 | 4.7 |
| 322 | >50 | >10 | ND |
| 323 | 28.3 | 8.9 | 3.2 |
| 324 | 27.5 | 1.9 | 14.2 |
| 327 | >50 | >10 | ND |
| 328 | 11.1 | 2.6 | 4.3 |
| 329 | >50 | >10 | ND |
| 330 | >50 | 6.2 | >8.1 |
| 331 | 47.0 | 9.9 | 4.7 |
| 332 | 31.2 | 8.0 | 3.9 |
| 333 | >50 | >10 | ND |
| 334 | 24.5 | 2.5 | 9.7 |
| 335 | >50 | >10 | ND |
| 336 | 15.9 | 2.4 | 6.6 |
| 337 | 32.1 | 3.5 | 9.3 |
| 338 | >50 | >10 | ND |
| 339 | 18.2 | 1.7 | 10.4 |
| 340 | >50 | >10 | ND |
| 341 | 26.5 | 4.0 | 6.6 |
| 352 | 29.3 | >10 | <2.9 |
| 353 | 17.5 | 4.2 | 4.2 |
| 354 | 22.0 | 0.5 | 44.9 |
| 355 | 21.2 | 2.0 | 10.5 |
| 356 | 15.5 | 2.0 | 7.6 |
| 357 | 13.1 | 1.2 | 10.7 |
| 358 | 13.2 | >10 | <1.3 |
| 362 | 20.0 | 2.2 | 9.0 |
| 363 | >50 | >10 | ND |
| 364 | 31.1 | 6.8 | 4.5 |
| 365 | 32.9 | >10 | <3.1 |
| 366 | 30.2 | >10 | <3.3 |
| 367 | >50 | >10 | ND |
| 368 | 12.1 | 1.6 | 7.6 |
| 370 | 19.5 | 2.6 | 7.5 |
| 371 | >50 | 8.9 | >5.6 |
| 372 | >50 | >10 | ND |
| 373 | >50 | 7.0 | >7.2 |
| 374 | 8.2 | 3.1 | 2.6 |
| 375 | 12.3 | 6.3 | 1.9 |
| 376 | >50 | >10 | ND |
| 377 | 14.6 | 8.3 | 1.8 |
| 378 | >50 | >10 | ND |
| 381 | 12.4 | 6.2 | 2.0 |
| 389 | >50 | >10 | ND |
| 392 | 25.9 | 4.7 | 5.5 |
| 394 | 30.9 | 5.5 | 5.6 |
| 395 | 37.4 | 7.6 | 4.9 |
| 396 | 23.3 | 6.1 | 3.8 |
| 397 | 27.6 | 8.1 | 3.4 |
| 402 | 26.9 | >10 | <3.0 |
| 403 | >50 | >10 | ND |
| 404 | >50 | >10 | ND |
| 405 | >50 | >10 | ND |
| 406 | >50 | >10 | ND |
| 407 | >50 | >10 | ND |
| 408 | 25.4 | 1.5 | 17.0 |
| 409 | 35.2 | 8.8 | 4.0 |
| 411 | >50 | >10 | ND |
| 412 | >50 | >10 | ND |
| 413 | >50 | >10 | ND |
| 414 | >50 | >10 | ND |
| 415 | >50 | >10 | ND |
| 416 | 18.2 | 1.3 | 14.2 |
| 417 | 24.4 | 3.0 | 8.1 |
| 418 | 12.6 | 3.0 | 4.2 |
| 419 | 21.6 | 2.7 | 7.9 |
| 420 | 7.0 | 1.0 | 6.7 |
| 421 | 19.0 | 1.5 | 12.5 |
| 422 | 6.1 | 2.5 | 2.4 |
| 423 | 3.4 | 6.0 | 0.6 |
| 424 | 6.5 | 2.4 | 2.7 |
| 425 | 5.5 | 2.2 | 2.5 |
| 426 | 31.9 | 4.4 | 7.3 |
| 427 | 29.5 | 6.1 | 4.8 |
| 428 | 11.4 | 8.9 | 1.3 |
| 429 | 12.1 | 2.4 | 5.1 |
| 430 | 16.5 | >10 | <1.6 |
| 431 | 22.4 | 4.8 | 4.7 |
| 432 | 5.6 | 2.9 | 2.0 |
| 433 | 27.0 | 3.8 | 7.1 |
| 434 | 24.6 | >10 | <2.5 |
| 435 | 26.9 | >10 | <2.7 |

**Supplemental Figure 2.**
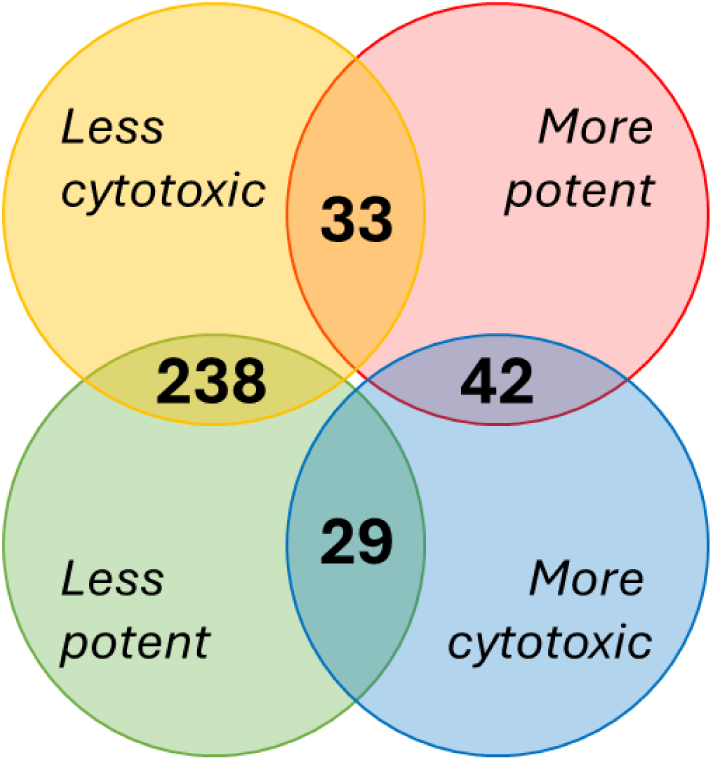
Venn diagram demonstrating compound potency against intracellular *L. donovani* (Lum IC_50_) and cytotoxicity against THP-1 host cell (24 hr LC_50_) relative to parental compound AR-12.

**Supplemental Figure 3.**
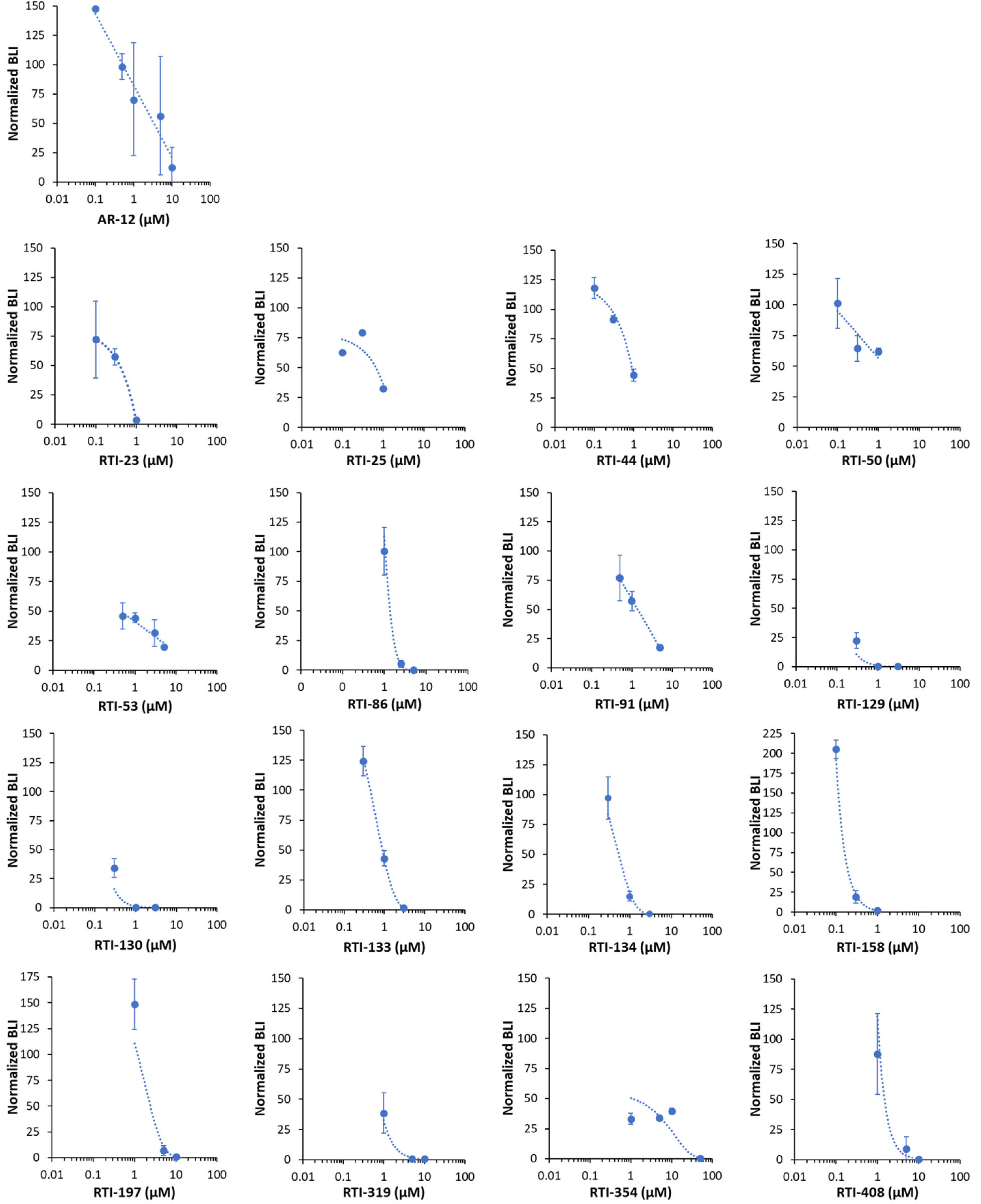
Luminescent activity of intracellular *L. donovani* infected THP1 macrophage cell after 72-hour incubation with compounds.

**Supplemental Figure 4.**
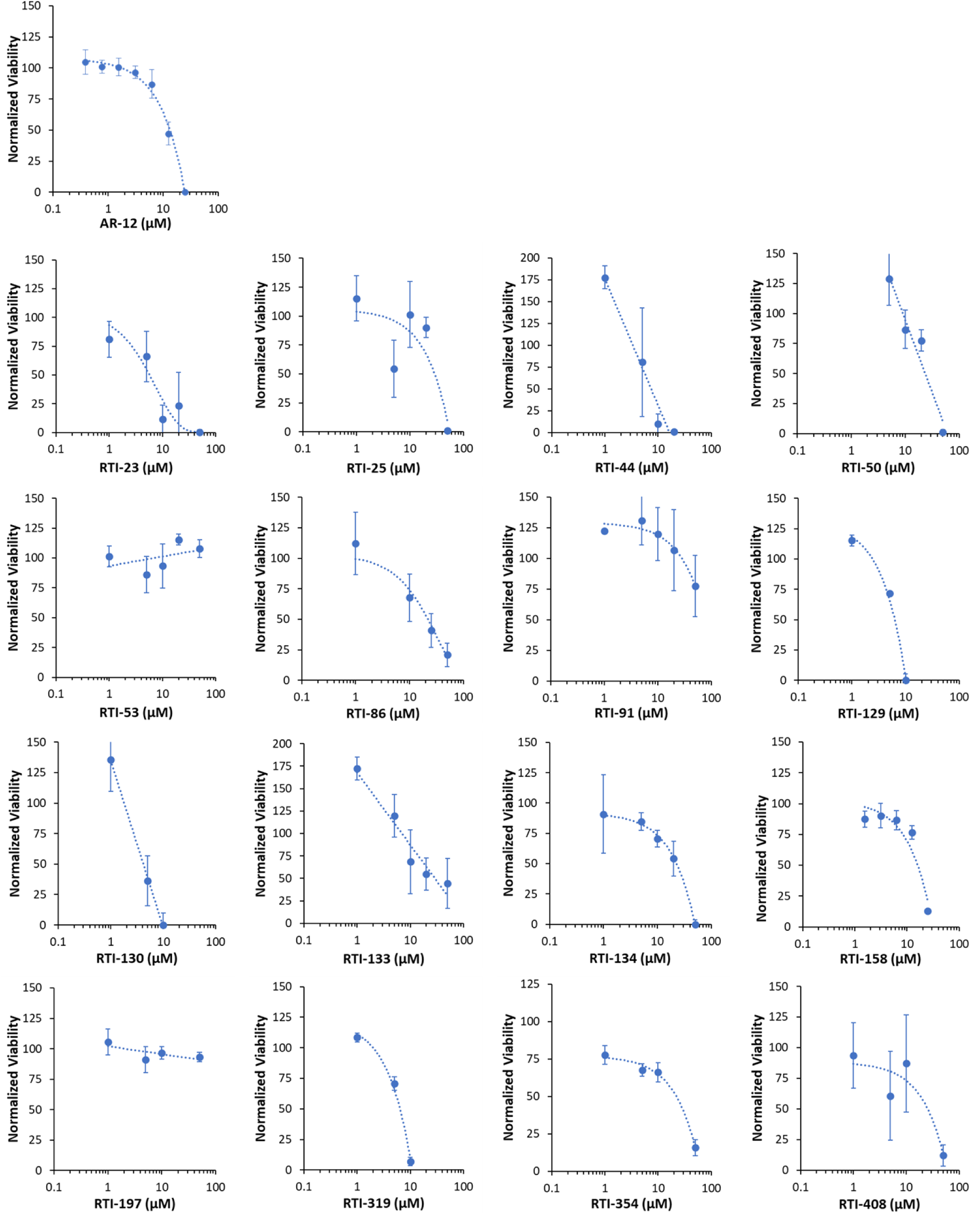
Graphs of THP1 macrophage cell viability after 24-hour incubation with compounds as determined by MTT assay.

**Supplemental Figure 5.**
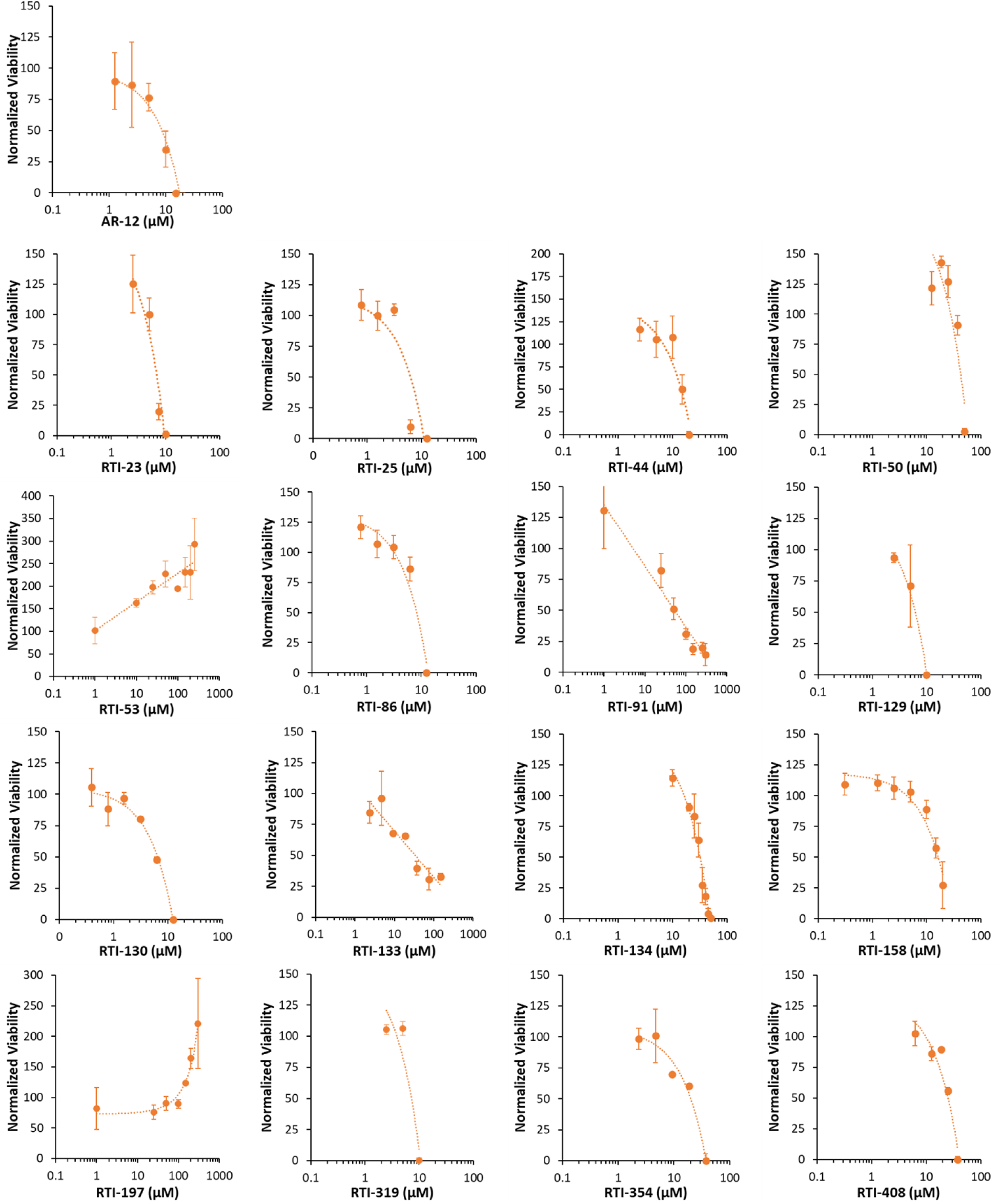
Graphs of THP1 macrophage cell viability after 72-hour incubation with compounds as determined by MTT assay.

**Supplemental Figure 6.**
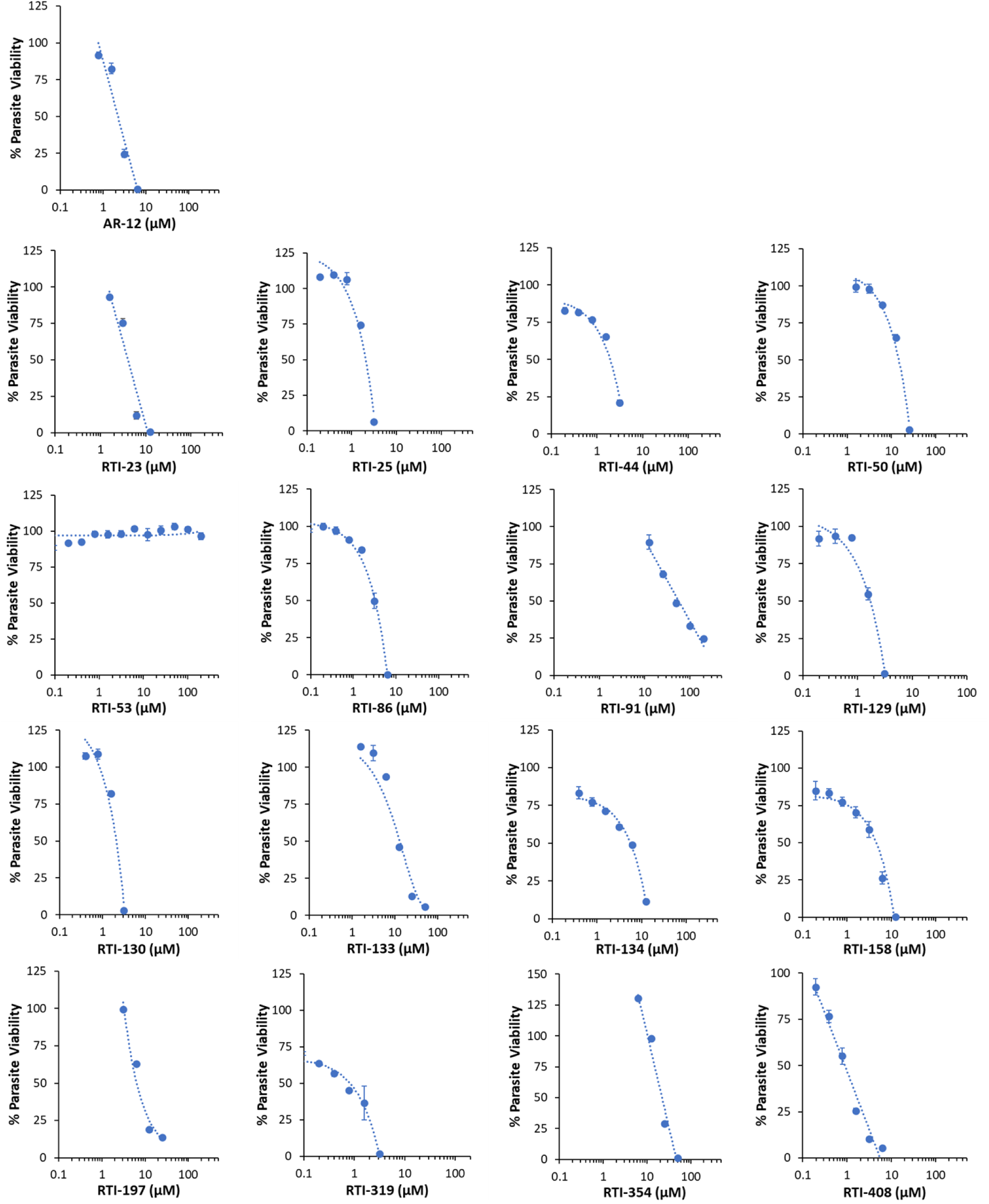
Dose response of extracellular *Leishmania* promastigote viability after 72-hour incubation with compound as measured by resazurin assay.

**Supplemental Figure 7.**
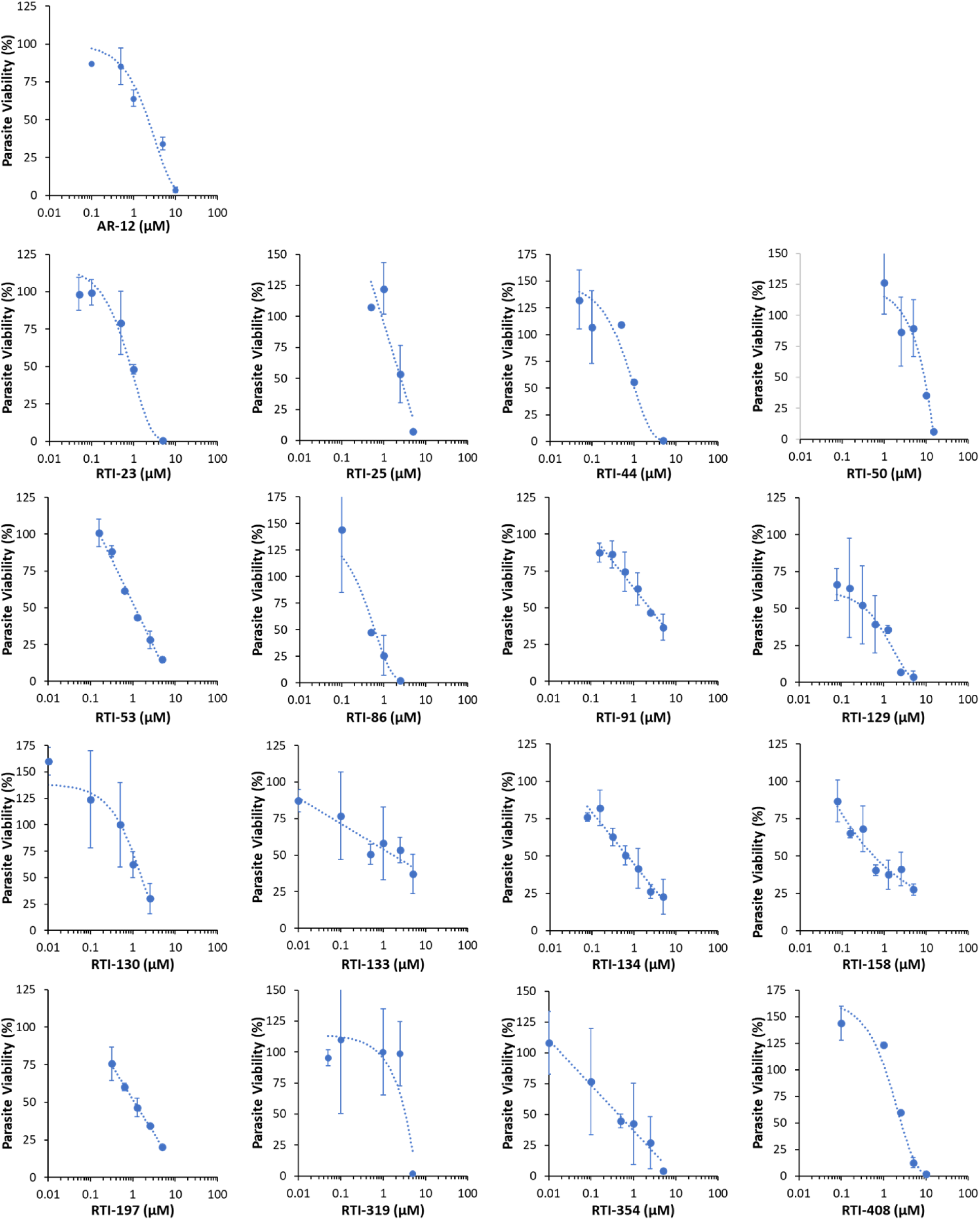
Dose response of intracellular *Leishmania donovani* burden in bone marrow derived macrophages after 72-hour incubation with compound as identified image-based giemsa staining.

**Supplemental Figure 8.**
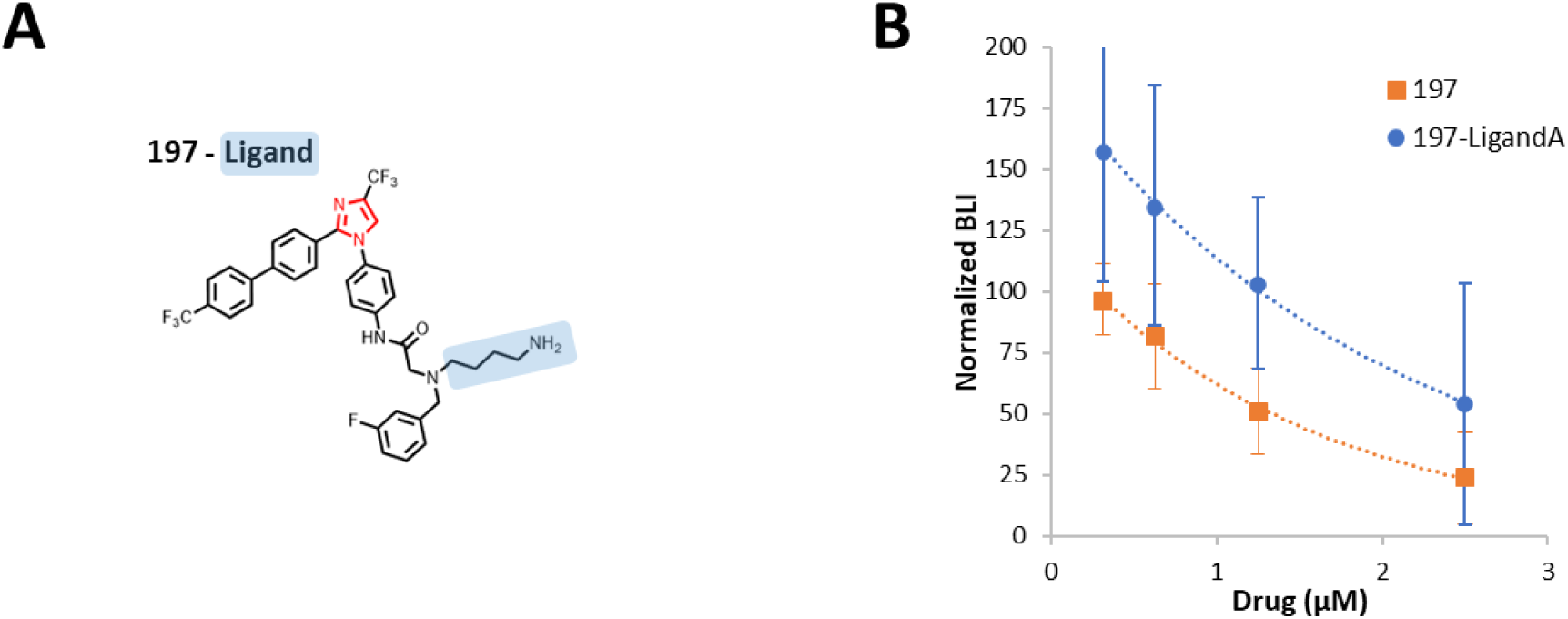
**A**) Chemical structure of 197 chemically modified for conjugation to agarose bead for affinity capture proteomic analysis. **B)** Luminescent activity of intracellular *L. donovani* infected THP1 macrophage cell after 72-hour incubation with 197 compared to chemically modified 197.

**Supplemental Figure 9.**
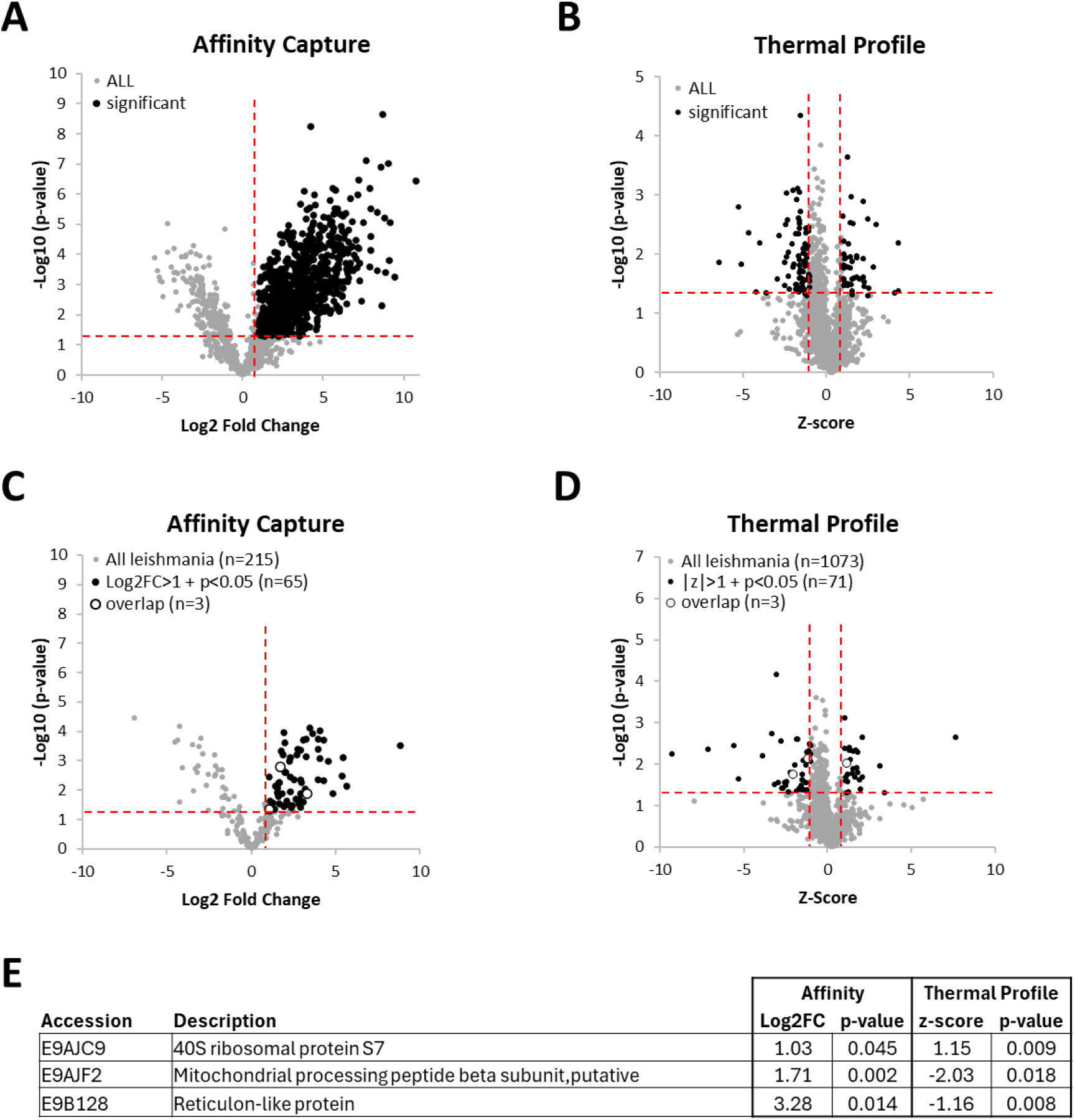
**A**) All human proteins (gray circle) identified by affinity capture using 197 functionalized bead plotted Log2(fold-change) and –Log10(p-value) over control bead. Significant proteins Log2FC > 1 and p< 0.05 are shown with black circles. **B)** All human proteins (gray circle) identified by thermal profile analysis with 197 plotted z-score and –Log10(p-value). Significant proteins with a |z-score| > 1 and p< 0.05 are shown with black circles. **C)** All *Leishmania* proteins (gray circle) identified by affinity capture using 197 functionalized bead plotted Log2(fold-change) and –Log10(p-value) over control bead. Significant proteins Log2FC > 1 and p< 0.05 are shown with black circles. Proteins with a (|z-score| > 1 and p< 0.05 overlapping with significant proteins identified by affinity capture (Log2FC > 2 and p< 0.05) are shown with open circles. **D)** All *Leishmania* proteins (gray circle) identified by thermal profile analysis with 197 plotted z-score and –Log10(p-value). Significant proteins with a |z-score| > 1 and p< 0.05 are shown with black circles. Significant proteins Log2FC > 1 and p< 0.05 are shown with black circles. Proteins with a (|z-score| > 1 and p< 0.05 overlapping with significant proteins identified by affinity capture (Log2FC > 2 and p< 0.05) are shown with open circles. **E)** Values for three *Leishmania* proteins overlapping between two proteomic approaches.

**Supplemental Figure 10.**
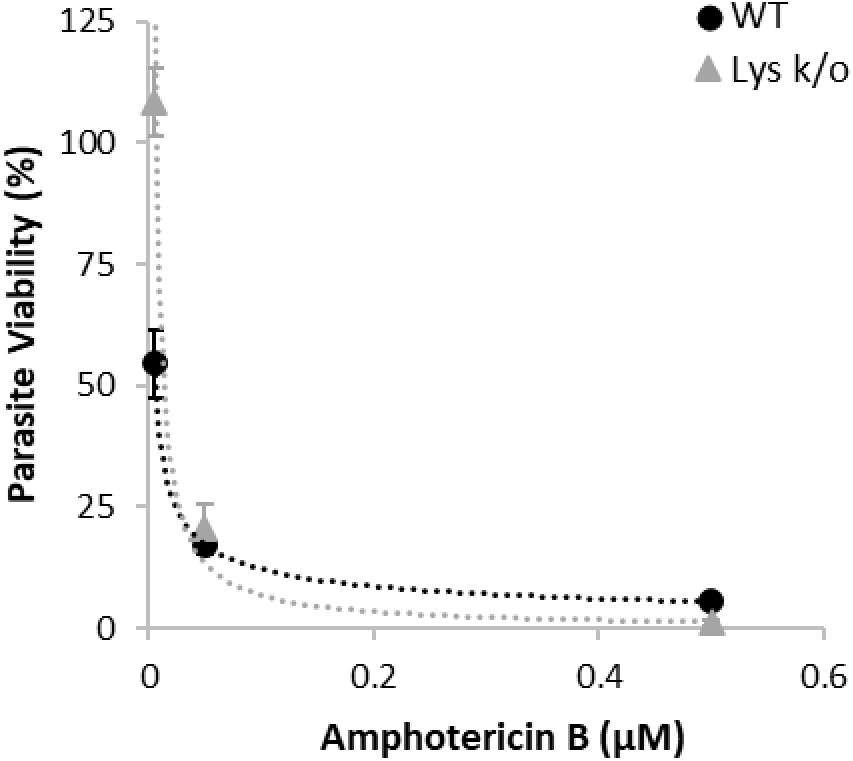
Dose response of amphotericin B on intracellular *Leishmania* burden in bone marrow derived macrophages derived from wildtype C57BL/6 (WT, black circle) or lysozyme knockout mice (Lys K/O, gray triangle) as identified image-based Giemsa staining. Data is presented as mean ± standard deviation of biological triplicates.

## Supplemental Information

## Synthesis and Characterization of Selected Analogs

Seven compounds were commercially available and purchased for this project. **RTI-121** (PRE-084 hydrochloride), **RTI-122** (SA4503 dihydrochloride), **RTI-123** (NE 100 hydrochloride), and **RTI-124** (BD 1047 dihydrobromide) were purchased from Tocris Bioscience, Inc. **RTI-125** (haloperidol) was acquired from Alfa Aesar. **RTI-278** ((S)-2-(3-fluorophenyl)pyrrolidine d-Tartrate) was sourced from AstaTech Inc. Lastly, **RTI-435** (notoginsenoside R1) was purchased from Sigma-Aldrich.

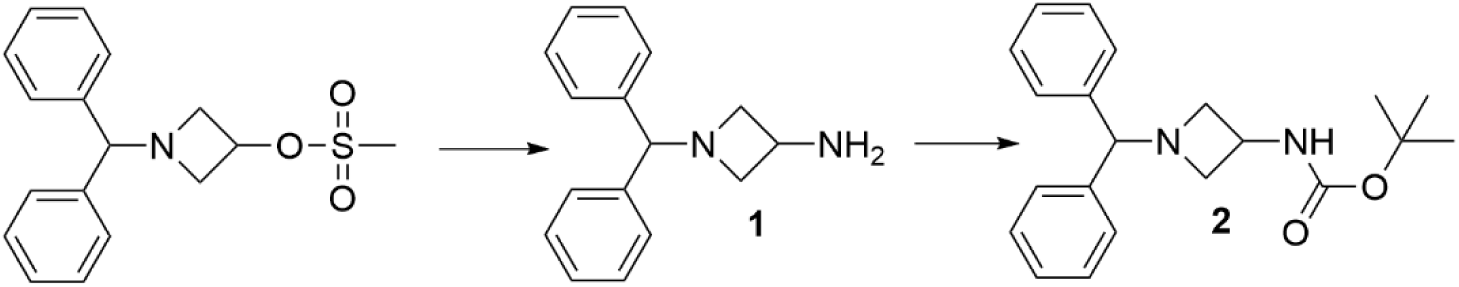

**<u>RTI-11</u>: In** a glass pressure reactor was combined 1-Benzhydrylazetidin-3-yl methanesulfonate (5.07 g, 15.9 mmol) and NH_4_OH (19.5 mL, 288.6 mmol) in 2-propanol (30 mL). The mixture was sealed with a Teflon cap and heated to 70° C for 3 hours. The reaction mixture was quenched with saturated aqueous NaHCO_3_ and extracted with ethyl acetate. The organic layer was washed with brine, dried (Na_2_SO_4_), filtered and concentrated to yield 3.76 g (100%) of **1** as an off-white gel, which required no further purification. ^1^H-NMR (CDCl_3_) δ 7.40-7.27 (m, 4 H), 7.24-7.15 (m, 6 H), 4.27 (s, 1 H), 3.67-3.50 (m, 3 H), 2.73-2.64 (m, 2 H).

A mixture containing **1** (3.76 g, 15.8 mmol) in tetrahydrofuran (75 mL) and 5% aqueous Na_2_CO_3_ (90 mL) was cooled to 0° C. A solution of di-tert-butyl decarbonate (4.48 g, 20.5 mmol) in tetrahydrofuran (15 mL) was added slowly and the reaction stirred at room temperature for 18 hours. Upon completion, the solvents were removed *in vacuo* and the residue was extracted with ethyl acetate. The organic layer was washed with brine, dried (MgSO_4_), filtered and concentrated to yield 6.03 g of a white solid. The crude material was stirred in hexanes and the resulting solid was filtered, washed with hexanes and dried to obtain 3.91 g (73%) of **2** (**RTI-11**) as a white solid. ^1^H-NMR (CDCl_3_) δ 7.39 (m, 4 H), 7.29-7.15 (m, 6 H), 4.86 (br s, 1 H), 4.32-4.27 (m, 2 H), 3.52 (t, 2 H, J = 9.0 Hz), 2.87-2.79 (m, 2 H), 1.42 (s, 9 H).

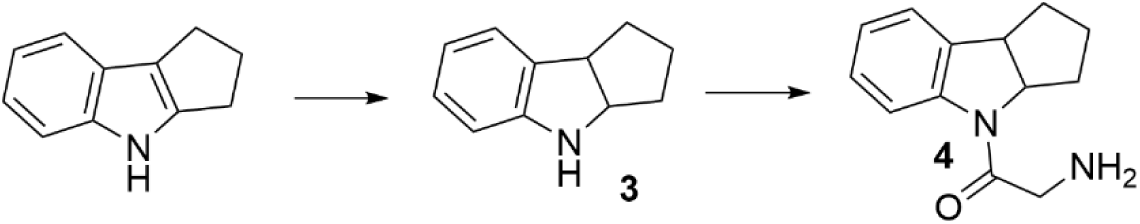

**<u>RTI-320:</u>** 1,2,3,4-Tetrahydrocyclopent[b]indole (4.0 g, 25.4 mmol) and 10% Palladium on Carbon (500 mg) were suspended in ethanol (50 mL) and concentrated hydrochloride acid (2.7 mL), charged with hydrogen gas and agitated on a Parr Apparatus at room temperature for 18 hours. The mixture was filtered through Celite and the pad was washed with methanol. The filtrate was concentrated, diluted with 1 N hydrochloric acid and extracted with diethyl ether. The aqueous layer was neutralized to pH of 8 with 2 N aqueous NaOH and extracted with dichloromethane. The organic layer was washed with brine, dried (MgSO_4_), filtered and concentrated to yield 3.05 g (75%) of **3** as a brown oil, which required no further purification. ^1^H-NMR (CDCl_3_) δ 7.02 (d, 1 H, J = 6.0 Hz), 6.98 (d, 1 H, J = 6.0 Hz), 6.67 (dd, 1 H, J = 3.0 Hz, 6.0 Hz), 6.52 (d, 1 H, J = 9.0 Hz), 4.36 (dd, 1 H, J = 3.0 Hz, 6.0 Hz), 3.77 (t, 2 H, J = 9.0 Hz), 2.02-1.51 (m, 6 H).

A mixture containing **3** (3.05 g, 19.15 mmol), 2-chloroacetamide (3.71 g, 39.6 mmol) and diisopropylethylamine (10.3 mL, 59.1 mmol) in N, N-dimethylformamide (8 mL), in a glass pressure reactor, was sealed with a Teflon cap and heated to 100° C for 18 hours. The mixture was diluted with water and extracted with ethyl acetate. The organic layer was washed with brine, dried (MgSO_4_), filtered and concentrated. The crude material was adsorbed onto Celite and purified over silica gel using 0-50% hexane/ethyl acetate yielding 3.74 g (86%) of **4** (**RTI-320**) as a yellow solid. ^1^H-NMR (CDCl_3_) δ 7.07 (dd, 2 H, J = 6.0 Hz), 6.74 (dd, 1 H, J = 6.0 Hz, 9.0 Hz), 6.50 (br s, 1 H), 6.34 (d, 1 H, J = 9.0 Hz), 5.49 (br s, 1 H), 4.20-4.05 (m, 2 H), 3.78 (dd, 2 H, J = 9.0 Hz), 2.04-1.49 (m, 6 H). LC-MS, calculated for C_13_H_16_N_2_O (MH)^+^ 217.3; observed 217.0. Anal. Calculated for C_13_H_16_N_2_O: C, 72.19; H, 7.45; N, 12.95. Found: C, 71.93; H, 7.43; N, 12.90.

### Novel Pyrazole Scaffolds (Type A: Standard Structure with N-substituted Anilines)

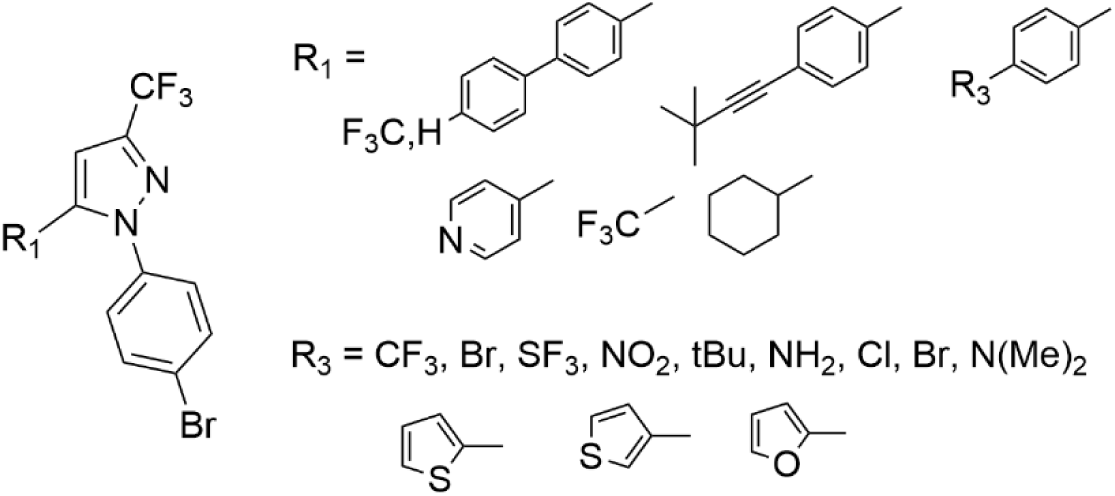

Examples:

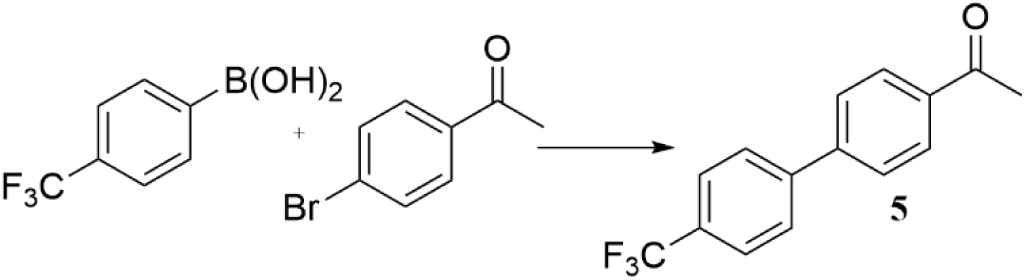

**<u>5</u>**: A mixture of 4-Trifluoromethylphenylboronic acid (7.84 g, 41.3 mmol), 4-Bromoacetophenone (8.25 g, 41.4 mmol), Palladium acetate (186 mg, 0.82 mmol), K_2_CO_3_ (17.1 g, 123.8 mmol) and Tetrabutylammonium bromide (17.3 g, 53.6 mmol) in THF (40 mL) and nitrogen gas was bubbled into the mixture for two minutes. Water (410 mL) was added and the reaction was heated to 60 C for 2 hours. Upon cooling to room temperature, the mixture was extracted with ethyl acetate. The organic material was washed with water and brine, dried (MgSO_4_), filtered and concentrated to yield a quantitative yield (11.2 g) of a copper-colored solid (**5**), which was pure enough for further synthesis. ^1^H-NMR (CDCl_3_) δ 8.05 (d, 2 H, J = 6 Hz), 7.79-7.69 (m, 6 H), 2.68 (s, 3 H).

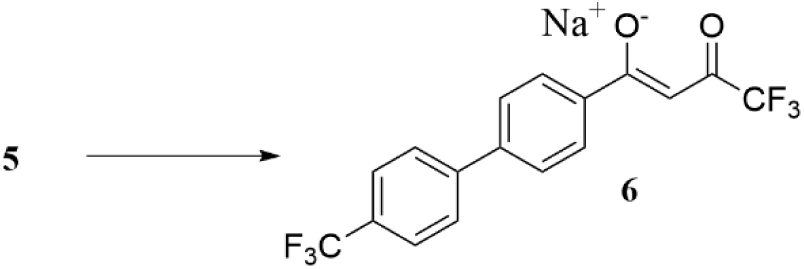

**<u>6</u>**: Into an oven dried flask was introduced sodium hydride (60% wt./mineral oil, 6.54 g, 272.4 mmol) and stirred in anhydrous THF (50 mL) for 5 minutes at room temperature. Ethyl trifluoroacetate (21.7 mL, 182.4 mmol) was added dropwise and this mixture stirred at room temperature for 10 minutes. A solution of **5** (24 g, 90.8 mmol) in anhydrous THF (85 mL) was added dropwise and the reaction mixture was refluxed for 3 hours. The reaction was concentrated and the residue was partitioned between ethyl acetate and water. The aqueous layer was extracted with ethyl acetate and the combined organics were washed with brine, dried (Na_2_SO_4_) and concentrated to yield a quantitative yield of a yellow solid (**6**), which was pure enough for the next step. ^1^H-NMR (CDCl_3_) δ 7.70 (d, 2 H, J = 6 Hz), 7.63 (d, 2 H, J = 9 Hz), 7.50 (d, 2 H, J = 9 Hz), 7.36 (d, 2 H, J = 9 Hz), 6.08 (s, 1 H).

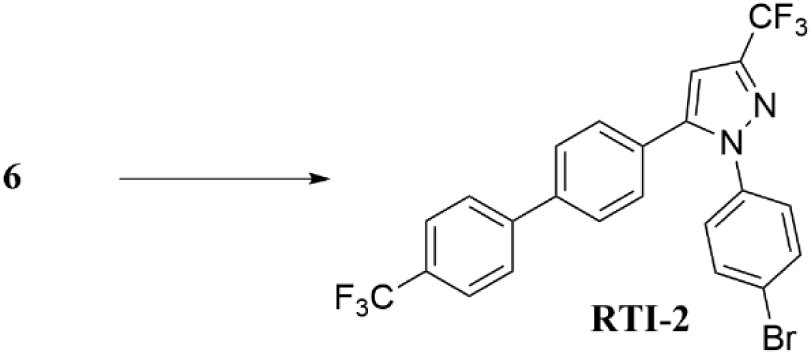

**<u>RTI-2:</u>** A mixture containing **6** (20 g, 52.3 mmol) and 4-bromohydrazine hydrochloride (16.4 g, 73.4 mmol) in ethanol (650 mL) was refluxed for 18 hours. The solvent was concentrated and the residue was partitioned between ethyl acetate and saturated aqueous NaHCO_3_. The aqueous layer was extracted with ethyl acetate and the combined organics were washed with brine, dried (MgSO_4_), filtered and concentrated. The crude material was purified over silica gel using 1-5% ethyl acetate from hexanes to yield 18.7 g of a yellow solid that contained both pyrazole isomers. The solid was crystallized from DCM/hexanes to yield 6.9 g (26%) of pure **RTI-2** as a yellow solid. ^1^H-NMR (CDCl_3_) δ 7.71 (dd, 4 H, J = 9 Hz), 7.60 (d, 2 H, J = 6 Hz), 7.53 (d, 2 H, J = 9 Hz), 7.31 (d, 2 H, J = 6 Hz), 7.25 (d, 2 H, J = 9 Hz), 6.81 (s, 1 H).

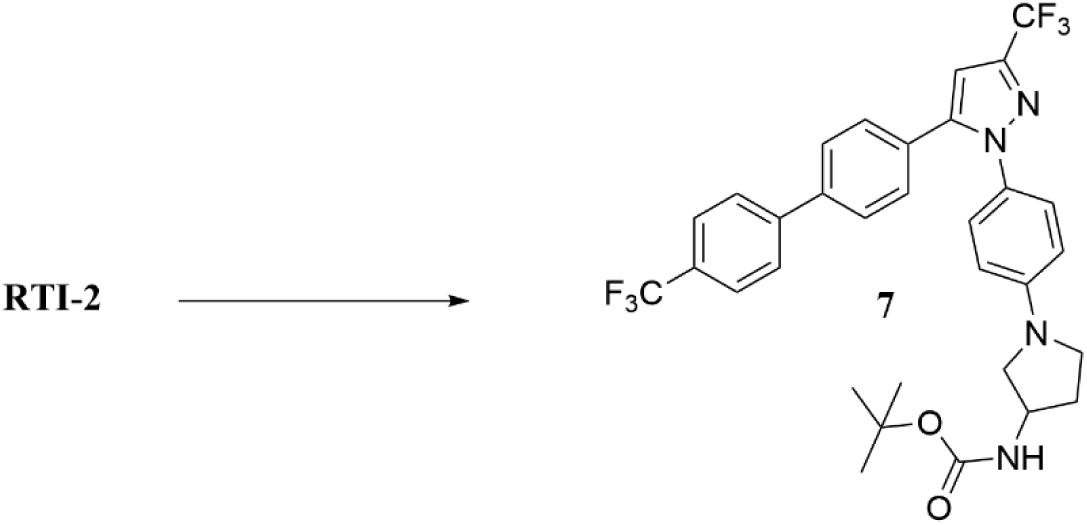

**<u>7</u>:** The following were combined in a heavy-duty glass reactor: **RTI-2** (4.65 g, 9.10 mmol), 3-N-Boc-aminopyrrolidine (2.72 g, 14.5 mmol), BINAP (1.7 g, 2.73 mmol), Pd_2_(dba)_3_ (1.08 g, 1.18 mmol) and Cs_2_CO_3_ (4.75 g, 14.5 mmol) in anhydrous toluene (95 mL) and nitrogen gas was bubbled into the mixture for two minutes. The reactor was then sealed with a Teflon cap and heated to 110° C for 18 hours. Upon cooling, the mixture was filtered through Celite and the filter pad was rinsed with ethyl acetate. The filtrate was washed with water and brine, dried (Na_2_SO_4_), filtered and concentrated. The crude material was purified over silica gel using 0-10% ethyl acetate from hexanes to yield 4.6 g (80%) of a yellow solid (**7**). ^1^H-NMR (CDCl_3_) δ 7.71 (dd, 4 H, J = 9 Hz), 7.56 (d, 2 H, J = 6 Hz), 7.35 (d, 2 H, J = 9 Hz), 7.18 (d, 2 H, J = 9 Hz), 6.78 (s, 1 H), 6.49 (d, 2 H, J = 9 Hz), 4.78-4.68 (m, 1 H), 4.42-4.33 (m, 1 H), 3.59 (dd, 1 H, J = 3 Hz, 6 Hz), 3.47-3.32 (m, 2 H), 3.17 (dd, 1 H, J = 3 Hz, 6 Hz), 2.33-2.26 (m, 1 H), 2.05-1.94 (m, 1 H), 1.46 (s, 9 H).

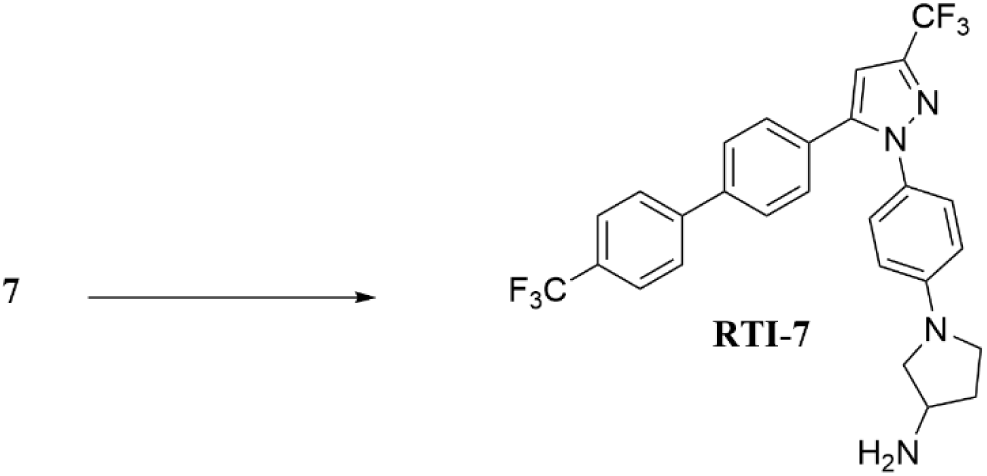

**<u>RTI-7:</u>** A solution of **7** (1.68 g, 2.72 mmol) in dichloromethane (30 mL) was cooled to 0° C and treated with trifluoroacetic acid (2.0 mL, 26.9 mmol). The reaction warmed to room temperature and stirred for 18 hours. Upon completion, the mixture was concentrated and the residue was partitioned between ethyl acetate and 2 N aqueous NaOH. The aqueous layer was extracted with ethyl acetate and the combined organic layers were washed with brine, dried (Na_2_SO_4_), filtered and concentrated to yield a tan solid (**RTI-7**, 1.34 g, 94%) that required no further purification. ^1^H-NMR (CDCl_3_) δ 7.66 (s, 4 H), 7.56 (d, 2 H, J = 9 Hz), 7.35 (d, 2 H, J = 9 Hz), 7.17 (d, 2 H, J = 9 Hz), 6.78 (s, 1 H), 6.49 (d, 2 H, J = 9 Hz), 3.75 (dd, 1 H, J = 6 Hz), 3.55-3.45 (m, 2 H), 3.35 (dd, 1 H, J = 6 Hz), 3.04 (dd, 1 H, J = 3 Hz, 6 Hz), 2.29-2.18 (m, 1 H), 1.88-1.78 (m, 1 H). ESI-MS, calculated for C_27_H_22_F_6_N_4_ (MH)^+^ 517.4; observed 517.6; Anal. Calculated for C_27_H_22_F_6_N_4_; C, 62.79; H, 4.29; N, 10.84. Found: C, 62.72; H, 4.29; N, 10.59.

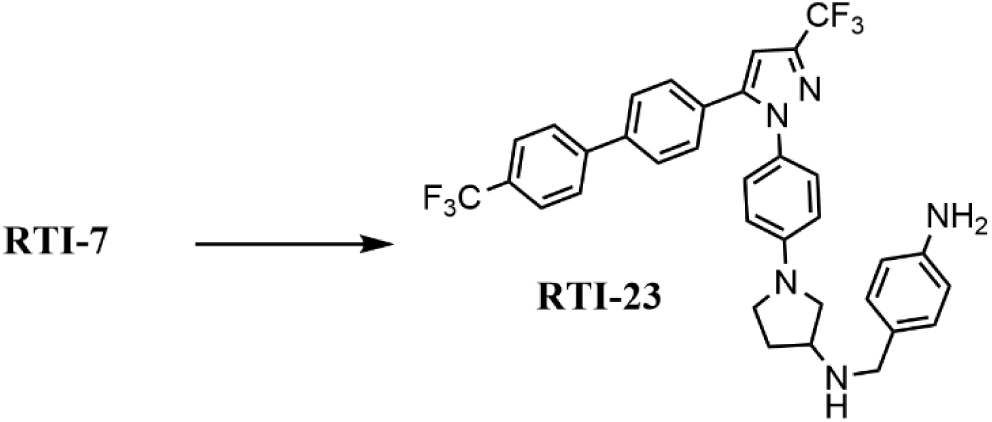

### Other Selected Analogs

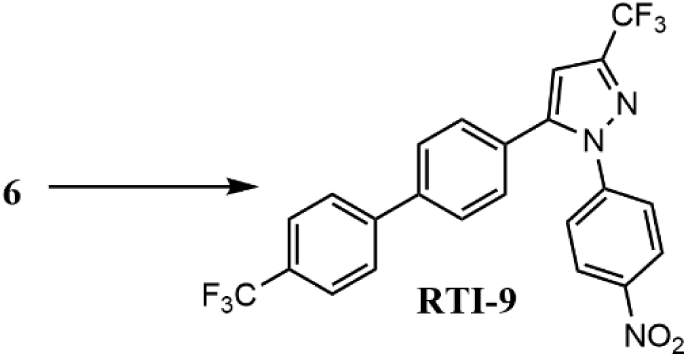

**<u>RTI-9:</u> A** mixture containing **6** (19.16 g, 50.1 mmol) and 4-nitrohydrazine hydrochloride (12.35 g, 65.1 mmol) in ethanol (600 mL) was refluxed for 18 hours. The solvent was concentrated and the residue was partitioned between ethyl acetate and saturated aqueous NaHCO_3_. The aqueous layer was extracted with ethyl acetate and the combined organics were washed with brine, dried (MgSO_4_), filtered and concentrated. The crude material was purified over silica gel using 1-5% ethyl acetate from hexanes to yield 10.58 g (44%) of pure **RTI-9** as a yellow solid. ^1^H-NMR (CDCl_3_) δ 8.27 (d, 2 H, J = 6 Hz), 7.75-7.69 (m, 4 H), 7.64 (d, 2 H, J = 9 Hz), 7.57 (d, 2 H, J = 6 Hz), 7.36 (d, 2 H, J = 9 Hz), 6.85 (s, 1 H). ESI-MS, calculated for C_23_H_13_F_6_N_3_O_2_ (MH)^+^ 477.4; observed 477.8; Anal. Calculated for C_23_H_13_F_6_N_3_O_2_; C, 57.87; H, 2.74; N, 8.80. Found: C, 57.61; H, 2.92; N, 8.71.

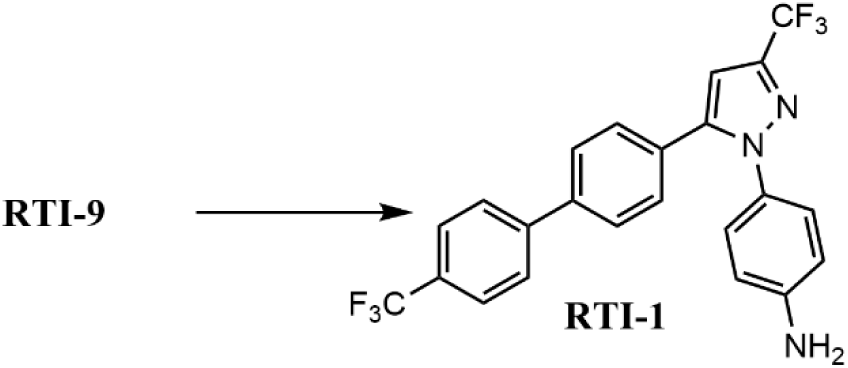

**<u>RTI-1:</u> A** mixture containing **RTI-9** (8.93 g, 18.7 mmol), Tin (II) chloride (12.4 g, 65.5 mmol) and concentrated hydrochloric acid (28 mL, 336 mmol) in ethanol (125 mL) was heated to 50 °C for 2.5 hours. The solvent was concentrated and the residue was partitioned between ethyl acetate and 2 N NaOH. The aqueous layer was extracted with ethyl acetate and the combined organics were washed with water and brine, dried (MgSO_4_), filtered and concentrated to yield 8.15 g (97 %) of **RTI-1** as a tan solid, which was used without any further purification. ^1^H-NMR (CDCl_3_) δ 7.68 (dd, 4 H, J = 9 Hz), 7.56 (d, 2 H, J = 9 Hz), 7.35 (d, 2 H, J = 9 Hz), 7.13 (d, 2 H, J = 9 Hz), 6.78 (s, 1 H), 6.66 (d, 2 H, J = 9 Hz), 3.82 (br s, 2 H).

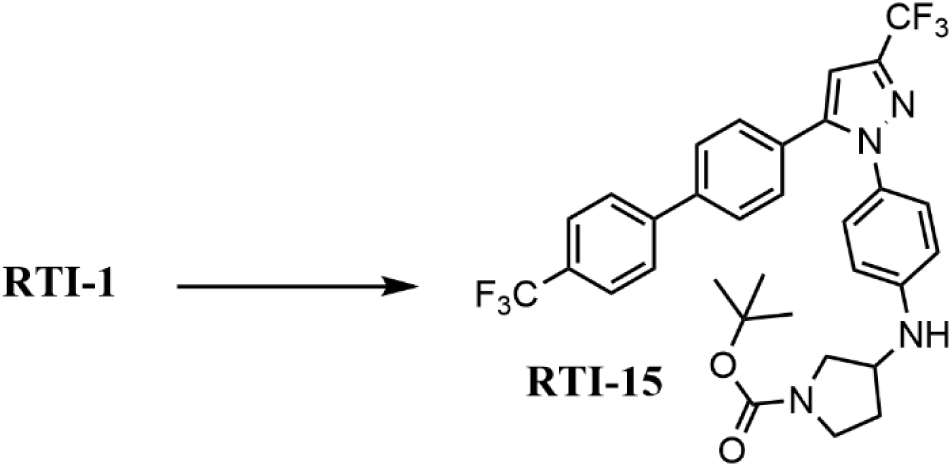

**<u>RTI-15:</u>** A mixture of **RTI-1** (6 g, 13.4 mmol) and N-Boc-3-pyrrolidinone (3.23 g, 17.4 mmol) in glacial acetic acid (110 mL) was treated with anhydrous sodium sulfate (11.43 g, 80.4 mmol); the mixture was then cooled to 0° C. Sodium triacetoxyborohydride (5.97 g, 28.2 mmol) was added and the reaction was stirred at room temperature for 18 hours. Upon completion, the reaction was concentrated and the residue was partitioned between ethyl acetate and saturated aqueous sodium bicarbonate. The aqueous layer was extracted with ethyl acetate. The organic layer was washed with brine, dried (Na_2_SO_4_), filtered and concentrated to yield 10.58 g of a black gel. The crude material was purified over silica gel using 0-20 % ethyl acetate from hexanes to yield 5.18 g (63 %) of **RTI-15** as a tan solid. ^1^H-NMR (CDCl_3_) δ 7.69 (s, 4 H), 7.60 (d, 2 H, J = 9 Hz), 7.36 (d, 2 H, J = 9 Hz), 7.16 (d, 2 H, J = 9 Hz), 6.78 (s, 1 H), 6.57 (d, 2 H, J = 9 Hz), 4.10-4.00 (m, 1 H), 3.98-3.90 (m, 1 H), 3.78-3.65 (m, 1 H), 3.56-3.45 (m, 2 H), 3.37-3.19 (m 1 H), 2.26-2.18 (m,1 H), 1.97-1.86 (m, 1 H), 1.46 (s, 9 H). ESI-MS, calculated for C_32_H_30_F_6_N_4_O_2_ (M+Na)^+^ 639.6; observed 639.5.

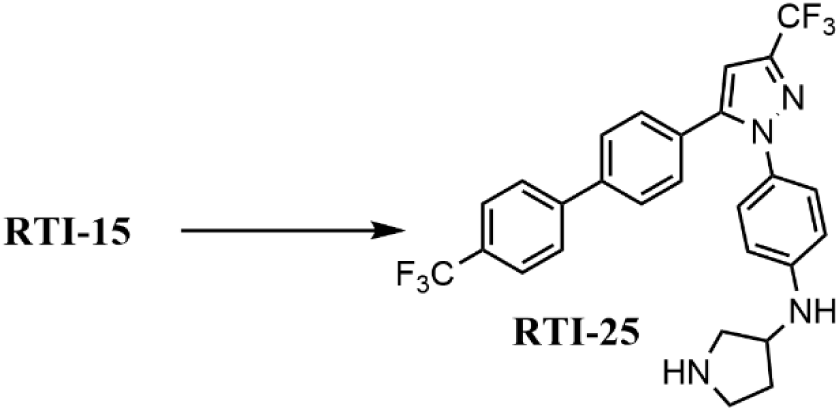

**<u>RTI-25:</u>** A solution of **RTI-15** (4.90 g, 7.96 mmol) in dichloromethane (55 mL) was cooled to 0° C and treated with trifluoroacetic acid (5.9 mL, 79.4 mmol). The reaction warmed to room temperature and stirred for 18 hours. Upon completion, the mixture was concentrated and the residue was partitioned between ethyl acetate and 2 N aqueous NaOH. The aqueous layer was extracted with ethyl acetate and the combined organic layers were washed with brine, dried (Na_2_SO_4_), filtered and concentrated to yield a brown solid (**RTI-25**, 4.0 g, 97%) that required no further purification. ^1^H-NMR (CDCl_3_) δ 7.68 (dd, 4 H, J = 9 Hz), 7.56 (d, 2 H, J = 9 Hz), 7.36 (d, 2 H, J = 9 Hz), 7.14 (d, 2 H, J = 9 Hz), 6.78 (s, 1 H), 6.56 (d, 2 H, J = 9 Hz), 4.05-3.91 (m, 2 H), 3.21-3.08 (m, 2 H), 3.01-2.93 (m, 1 H), 2.91-2.85 (m, 1 H), 2.27-2.15 (m, 1 H). Anal. Calculated (with 0.8 mol of water) for C_27_H_22_F_6_N_4_; C, 61.08; H, 4.48; N, 10.55. Found: C, 61.23; H, 4.30; N, 10.43.

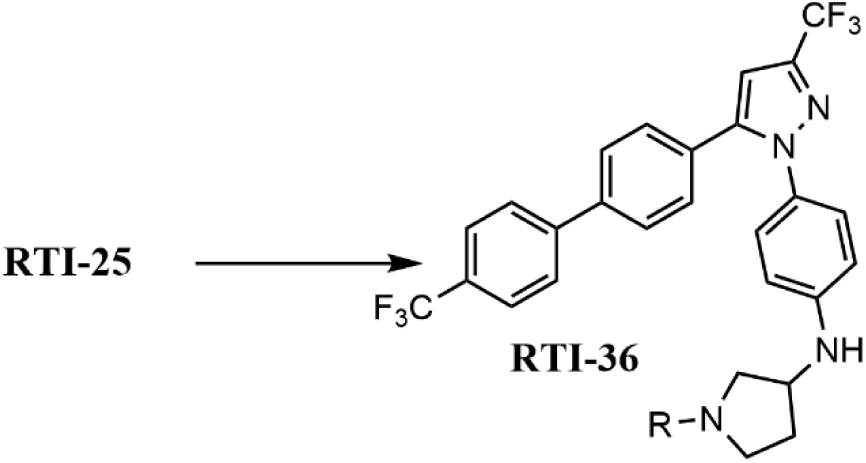

**<u>RTI-36 [R = CH_2_-(4-biphenyl)]:</u>** A solution of **RTI-25** (150 mg, 0.29 mmol), biphenyl 4-carboxaldehyde (58 mg, 0.32 mmol) and 4A molecular sieves (150 mg) in anhydrous 1,2-dichloroethane (4 mL) was stirred at room temperature for 5 hours. Sodium triacetoxyborohydride (136 mg, 0.64 mmol) was added and the reaction stirred for 18 hours at room temperature. The reaction was quenched with saturated aqueous sodium bicarbonate solution and extracted with ethyl acetate and the combined organic layers were washed with brine, dried (Na_2_SO_4_), filtered and concentrated to yield 176 mg of a brown solid. The crude material was purified over silica gel using 0-40 % ethyl acetate from hexanes to yield 85.7 mg (43%) of **RTI-36** as a white solid. ^1^H-NMR (CDCl_3_) δ 7.68 (dd, 4 H, J = 9 Hz), 7.60-7.53 (m, 6 H), 7.46-7.33 (m, 7 H), 7.12 (d, 2 H, J = 6 Hz), 6.77 (s, 1 H), 6.54 (d, 2 H, J = 9 Hz), 4.11 (dd, 1 H, J = 6 Hz), 4.01 (br s, 1 H), 3.68 (s, 2 H), 2.86-2.75 (m, 2 H), 2.62 (d, 1 H, J = 3 Hz, 6 Hz), 2.48 (dd, 1 H, J = 6 Hz, 9 Hz), 2.34 (dd, 1 H, J = 6 Hz, 9 Hz), 1.77-1.65 (m, 1 H). ESI-MS, calculated for C_40_H_32_F_6_N_4_ (MH)^+^ 683.7; observed 683.7.

**<u>RTI-53 [R = CH_2_-(3-cyanophenyl)]:</u> Using** 3-cyanobenzaldehyde, the product was isolated as a white solid in 58% yield (142 mg). ^1^H-NMR (CDCl_3_) δ 7.65 (dd, 5 H, J = 12 Hz), 7.55 (d, 2 H, J = 9 Hz), 7.43 (m, d, 1 H, J = 6 Hz), 7.35 (dd, 2 H, J = 9 Hz), 7.14 (dd, 2 H, J = 9 Hz), 6.77 (s, 1 H), 6.55 (d, 2 H, J = 9 Hz), 4.16-4.03 (m, 2 H), 3.65 (s, 2 H), 2.84-2.71 (m, 2 H), 2.59 (dd, 1 H, J = 9 Hz), 2.46-2.28 (m, 2 H), 1.76-1.65 (m, 1 H). ESI-MS, calculated for C_35_H_27_F_6_N_5_ (MH)^+^ 632.6; observed 632.6.

**<u>RTI-91 [R = CH_2_-(2-hydroxyphenyl)]:</u> Using** 2-hydroxybenzaldehyde, the product was isolated as an off-white solid in 48% yield (138.6 mg). ^1^H-NMR (CDCl_3_) δ 7.68 (dd, 4 H, J = 9 Hz), 7.56 (d, 2 H, J = 9 Hz), 7.35 (dd, 2 H, J = 9 Hz), 7.16 (dd, 3 H, J = 6 Hz, 9 Hz), 6.99 (d, 1 H, J = 9 Hz), 6.84-6.76 (m, 3 H), 6.53 (d, 2 H, J = 9 Hz), 4.06-3.97 (m, 2 H), 3.84 (s, 2 H), 2.97-2.85 (m, 2 H), 2.70 (dd, 1 H, J = 3 Hz, 6 Hz), 2.56 (dd, 1 H, J = 9 Hz), 2.42 (dd, 1 H, J = 6 Hz, 9 Hz), 1.78-1.72 (m, 1 H). ESI-MS, calculated for C_34_H_28_F_6_N_4_O (MH)^+^ 623.6; observed 623.8.

### Preparation of Novel Pyrazole Scaffolds (Type B: Left-Ring Substitution with N-substituted Anilines)

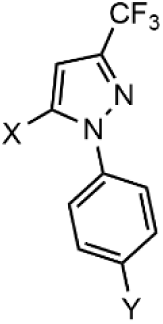

Following procedures described above, a few examples include:

**<u>RTI-247 (X = Cyclohexyl, Y = 3-Aminopyrrolidinyl):</u>** Using 1-Cyclohexylethanone in the procedure for **RTI-7**, **RTI-247** was isolated as a tan oil in 65% yield (125 mg). ^1^H-NMR (CDCl_3_) δ 7.26 (d, 2 H, J = 9 Hz), 6.54 (d, 2 H, J = 9 Hz), 6.53 (s, 1 H), 3.81-3.71 (m, 1 H), 3.61-3.49 (m, 2 H), 3.05 (dd, 1 H, J = 3 Hz, 6 Hz), 2.79-2.67 (m, 1 H), 2.32-2.21 (m, 1 H), 2.08-2.00 (m, 2 H), 1.91-1.71 (m, 4 H), 1.60-1.22 (m, 5 H). ESI-MS, calculated for C_20_H_25_F_3_N_4_ (MH)^+^ 379.4; observed 379.4; Anal. Calculated for C_20_H_25_F_3_N_4_; C, 63.47; H, 6.65; N, 14.80. Found: C, 63.31; H, 6.50; N, 14.57.

**<u>RTI-272 (X = Cyclopropyl, Y *=* 3-Aminopyrrolidinyl)</u>**: **RTI-272** was isolated as a tan solid (139.6 mg, 24%, over 2 steps). ^1^H-NMR (CDCl_3_) δ 7.34 (d, 2 H, J = 6 Hz), 6.64 (d, 2 H, J = 6 Hz), 6.14 (s, 1 H), 4.07-3.94 (m, 2 H), 3.19-3.07 (m, 2 H), 3.00-2.85 (m, 2 H), 2.28-2.14 (m, 1 H), 1.79-1.68 (m, 3 H), 1.02-0.95 (m, 2 H), 0.78-0.73 (m, 2 H). NMR (CDCl_3_, 75 MHz) δ 147.6, 129.2, 126.6, 112.9, 99.2, 54.0, 53.9, 45.8, 33.7, 8.8, 7.4. ESI-MS, calculated for C_17_H_19_F_3_N_4_ (MH)^+^ 337.4; observed 337.4. Anal. Calculated for C_17_H_19_F_3_N_4_; C, 60.70; H, 5.69; N, 16.65. Found: C, 60.49; H, 5.80; N, 16.50.

**<u>RTI-273 (X = 2-Furanyl, Y *=* 3-Aminopyrrolidinyl)</u>**: **RTI-273** was isolated as a tan solid (382 mg, 37%, over 2 steps). ^1^H-NMR (CDCl_3_) δ 7.42 (d, 2 H, J = 3 Hz), 7.20 (d, 2 H, J = 6 Hz), 6.87 (s, 1 H), 6.63 (d, 2 H, J = 6 Hz), 6.33 (d, 1 H, J = 3 Hz), 5.92 (d, 1 H, J = 3 Hz), 4.14-3.95 (m, 2 H), 3.22-3.08 (m, 2 H), 3.01-2.86 (m, 2 H), 2.29-2.19 (m, 1 H), 1.76-1.67 (m, 1 H). NMR (CDCl_3_, 75 MHz) δ 148.3, 143.6, 142.8, 136.4, 129.1, 127.4, 119.4, 112.9, 111.3, 109.3, 102.8, 53.9, 53.8, 45.8, 33.6. ESI-MS, calculated for C_18_H_17_F_3_N_4_O (MH)^+^ 363.3; observed 363.4. Anal. Calculated for C_18_H_17_F_3_N_4_O; C, 59.75; H, 4.73; N, 15.48. Found: C, 59.36; H, 4.84; N, 15.52.

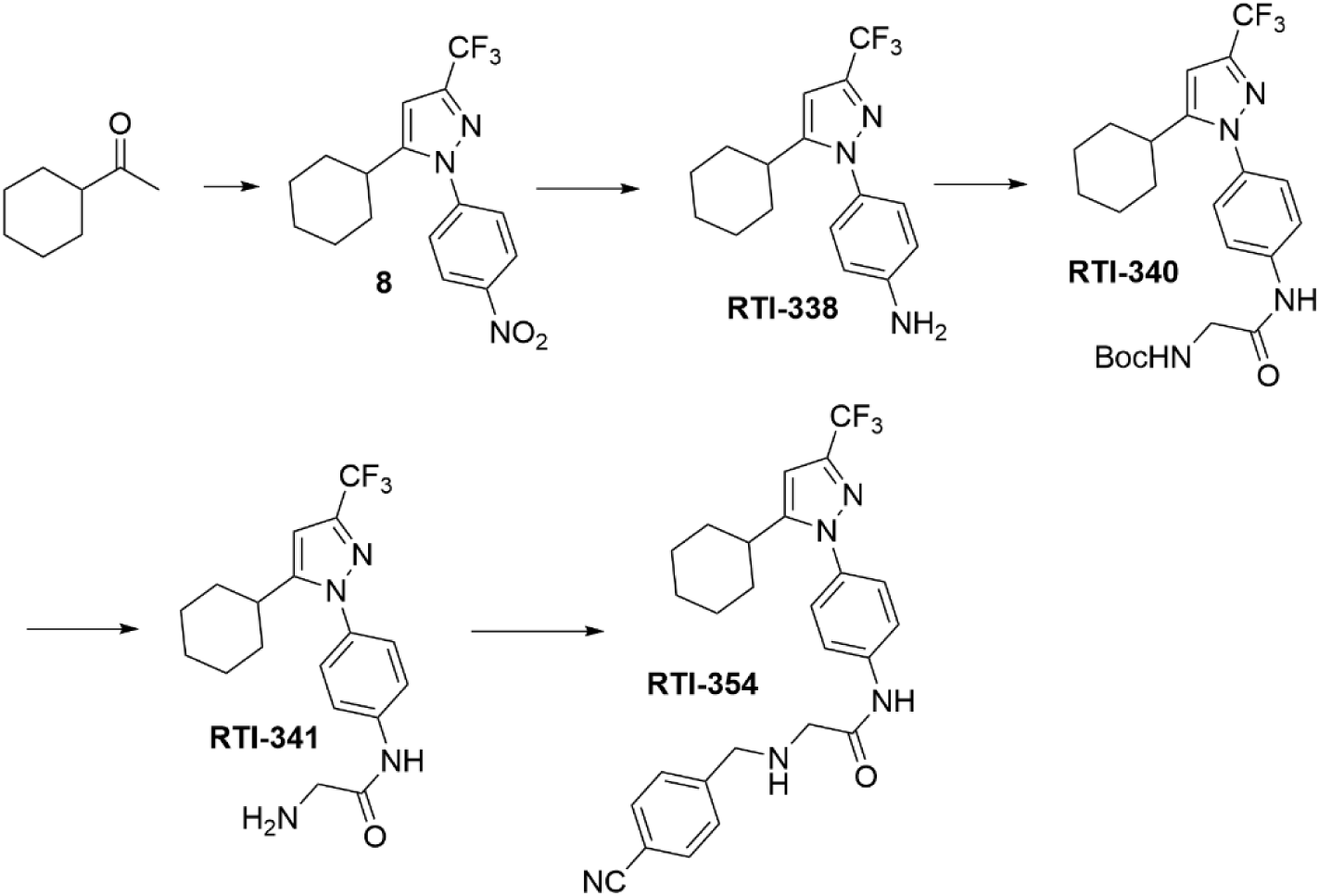

**<u>8</u>:** Into an oven dried flask was introduced sodium hydride (60% wt./mineral oil, 9.52 g, 237.6 mmol) and stirred in anhydrous THF (140 mL) for 5 minutes at room temperature. Ethyl trifluoroacetate (18.9 mL, 158.8 mmol) was added dropwise and this mixture stirred at room temperature for 10 minutes. A solution of cyclohexyl methyl ketone (10 g, 79.2 mmol) in anhydrous THF (40 mL) was added dropwise and the reaction mixture was refluxed for 3 hours. The reaction was concentrated and the residue was partitioned between ethyl acetate and water. The aqueous layer was extracted with ethyl acetate and the combined organics were washed with brine, dried (Na_2_SO_4_) and concentrated to yield a quantitative yield of a yellow oil, which was used without further purification. ^1^H-NMR (CD_3_OD) δ 5.24 (s, 1 H), 2.94-2.84 (m, 1 H), 1.82-1.66 (m, 4 H), 1.40-1.19 (m, 6 H). Using this material, the procedure for **RTI-2** was followed, substituting 4-nitrohydrazine hydrochloride. During pyrazole formation, two different pyrazoles may form. The product (**8**, the less polar isomer, with approximate R_f_ = 0.7) was isolated as a yellow oil in 59% yield (15.9 g). ^1^H-NMR (CDCl_3_) δ 8.36 (d, 2 H, J = 9 Hz), 7.73 (d, 2 H, J = 9 Hz), 6.72 (s, 1 H), 2.77-2.69 (m, 1 H), 1.86-1.73 (m, 4 H), 1.59-1.22 (m, 6 H).

**<u>RTI-338:</u> A** mixture containing **8** (21.8 mmol), stannous chloride (14.5 g, 76.3 mmol) and concentrated HCl (33 mL, 396 mmol) in ethanol (145 mL) was heated to 50° C for 2.5 hours. Upon cooling to room temperature, the solvent was concentrated and the residue was diluted with ethyl acetate and 2 N NaOH (300 mL) was stirred in the mixture stirred at room temperature for 1h. The aqueous layer was extracted with ethyl acetate and the combined organics were washed with water and brine, dried (MgSO_4_), filtered and concentrated. The crude material was adsorbed onto silica gel and purified via ISCO using 1-20% ethyl acetate from hexanes to yield 3.54 g (52%) of a white solid (**RTI-338**). ^1^H-NMR (CDCl_3_) δ 7.22 (d, 2 H, J = 9 Hz), 6.70 (d, 2 H, J = 9 Hz), 6.54 (s, 1 H), 3.83 (br s, 2 H), 2.74-2.65 (m, 1 H), 2.05-1.96 (m, 2 H), 1.84-1.71 (m, 2 H), 1.51-1.31 (m, 6 H). ^13^C NMR (CDCl_3_, 75 MHz) δ 147.1, 127.0, 114.7, 105.2, 60.3, 37.3, 33.0, 26.2, 26.0, 14.1; ESI-MS, calculated for C_16_H_18_F_3_N_3_ (MH)^+^ 310.3; observed 310.0. Anal. Calculated for C_16_H_18_F_3_N_3_; C, 62.12; H, 5.86; N, 13.58. Found: C, 62.06; H, 5.98; N, 13.44.

**<u>RTI-340:</u>** A mixture containing **RTI-338** (3.03 g, 9.79 mmol), Boc-glycine (3.77 g, 21.5 mmol), diisopropylethylamine (6.0 mL, 34.4 mmol), 4-N,N-dimethylaminopyridine (180 mg, 1.47 mmol) and N-(3-dimethylaminopropyl)-n-ethylcarbodiimide hydrochloride (4.13 g, 21.5 mmol) in tetrahydrofuran (90 mL) was stirred at room temperature for 18 hours. The solvent was concentrated and the residue was partitioned between ethyl acetate and 1 N HCl. The aqueous layer was extracted with ethyl acetate and the combined organics were washed with saturated aqueous NaHCO_3_, water and brine, dried (Na_2_SO_4_), filtered and concentrated. The crude material was adsorbed onto silica gel and purified via ISCO using 3-20% ethyl acetate from hexanes to yield 3.60 g (72%) of a white solid (**RTI-340**). ^1^H-NMR (CDCl_3_) δ 8.35 (br s, 1 H), 7.64 (d, 2 H, J = 9 Hz), 7.43 (d, 2 H, J = 9 Hz), 6.59 (s, 1 H), 5.22 (br s, 1 H), 3.94 (d, 2 H, J = 6 Hz), 2.71 (dd, 1 H, J = 3 Hz, 9 Hz), 2.07-2.00 (m, 2 H), 1.84-1.62 (m, 3 H), 1.49 (s, 9 H), 1.44-1.24 (m, 5 H). ^13^C NMR (CDCl_3_, 75 MHz) δ 126.3, 119.8, 37.3, 33.0, 28.2, 26.2, 26.0; ESI-MS, calculated for C_23_H_29_F_3_N_4_O_3_ (MH)^+^ 467.5; observed 467.0. Anal. Calculated for C_23_H_29_F_3_N_4_O_3_; C, 59.21; H, 6.26; N, 12.01. Found: C, 59.12; H, 6.26; N, 12.16.

**<u>RTI-341:</u>** A solution of **RTI-340** (3.36 g, 7.84 mmol) in dichloromethane (80 mL) was cooled to 0° C and treated with trifluoroacetic acid (5.9 mL, 79.4 mmol). The reaction warmed to room temperature and stirred for 18 hours. Upon completion, the mixture was concentrated and the residue was partitioned between ethyl acetate and 2 N aqueous NaOH. The aqueous layer was extracted with ethyl acetate and the combined organic layers were washed with brine, dried (Na_2_SO_4_), filtered and concentrated to yield a tan solid. The crude material was purified over silica gel using 0-5% methanol from dichloromethane to yield 2.60 g (90%) of a white solid. Some of the crude material (400 mg) was adsorbed onto silica gel and purified via ISCO using 0-20% methanol from dichloromethane to yield 372 mg (93%) of a white solid (**RTI-341**). ^1^H-NMR (CDCl_3_) δ 9.59 (br s, 1 H), 7.73 (d, 2 H, J = 9 Hz), 7.44 (d, 2 H, J = 9 Hz), 6.59 (s, 1 H), 3.50 (s, 2 H), 2.72 (dd, 1 H, J = 3 Hz, 9 Hz), 2.08-1.96 (m, 2 H), 1.88-1.71 (m, 3 H), 1.52-1.33 (m, 5 H). ^13^C NMR (CDCl_3_, 75 MHz) δ 170.7, 158.6, 138.2, 136.5, 126.3, 119.3, 105.9, 45.1, 37.3, 33.0, 28.2, 26.2, 26.0; ESI-MS, calculated for C_18_H_21_F_3_N_4_O (MH)^+^ 367.3; observed 367.0. Anal. Calculated for C_18_H_21_F_3_N_4_O; C, 59.00; H, 5.77; N, 15.29. Found: C, 58.97; H, 5.81; N, 15.17.

### Examples of Imidazole Structure Synthesis

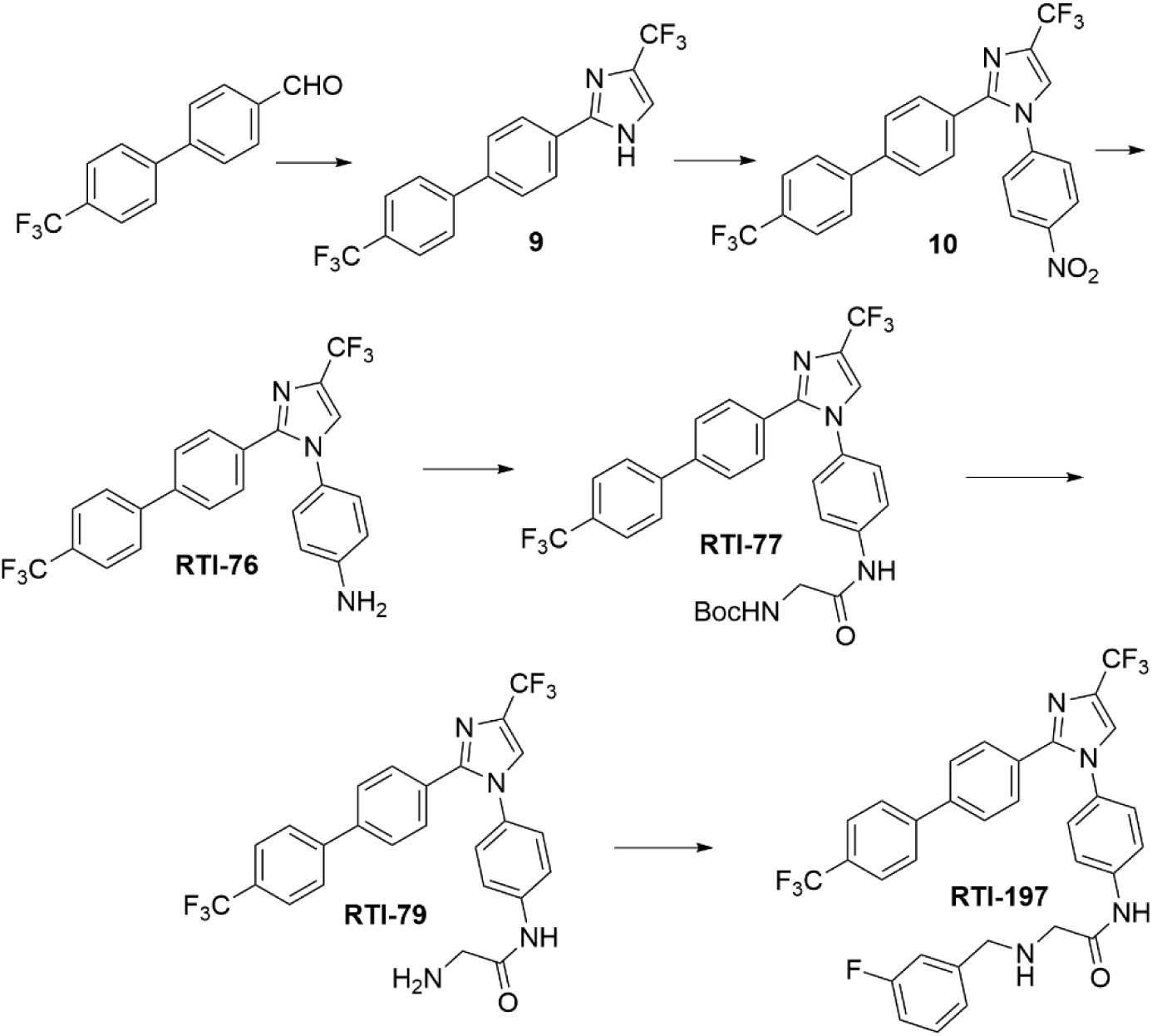

**<u>9</u>:** A mixture containing 1,1-Dibromo-3,3,3-trifluoroacetone (3.3 g, 12.0 mmol) and sodium acetate (1.32 g, 16.0 mmol) in water (42 mL) was heated to 95° C for 1 hour, then cooled to room temperature. 4’-Trifluoromethylbiphenyl-4-carboxaldehyde (2.0 g, 8.0 mmol) in methanol (102 mL) was cooled to 0° C and treated with dropwise addition of ammonium hydroxide (49 mL, 725.2 mmol). This mixture stirred for 10 minutes at 0° C and then the first mixture was added dropwise; the combined reaction slowly warmed to room temperature and stirred for 18 hours. The resulting solids were filtered, washed with water and dried and the filtrate was concentrated. The dried solids and filtrate residue were partitioned between dichloromethane and water. The organic layer was washed with brine, dried (Na_2_SO_4_) and concentrated. The crude material was adsorbed onto silica gel and purified via ISCO using 0-30% ethyl acetate from hexanes to yield 727 mg (25%) of a yellow solid (**9**). ^1^H-NMR (CDCl_3_) δ 8.00-7.95 (m, 1 H), 7.75-7.68 (m, 7 H), 7.64-7.57 (m, 2 H); LC-MS, calculated for C_17_H_10_F_6_N_2_ (MH)^+^ 357.2; observed 357.0.

**<u>10:</u>** A mixture containing **9** (727 mg, 2.04 mmol), 4-fluoronitrobenzene (864 mg, 6.12 mmol) and K_2_CO_3_ (1.0 g, 7.23 mmol) in N, N-dimethylformamide (21 mL) was heated to 90° C for 21 hours. Upon cooling to room temperature, the reaction was poured into saturated aqueous LiCl solution and extracted with ethyl acetate. The combined organics were washed with water (5 x) and brine, dried (MgSO_4_), filtered and concentrated. The crude material was adsorbed onto silica gel and purified via ISCO using 1-20% ethyl acetate from hexanes to yield 465 mg (48%) of an off-white solid (**10**). ^1^H-NMR (CDCl_3_) δ 8.36 (d, 2 H, J = 9 Hz), 7.70 (dd, 4 H, J = 6 Hz, 9 Hz), 7.57 (d, 3 H, J = 6 Hz), 7.49 (d, 4 H, J = 9 Hz); ESI-MS, calculated for C_23_H_13_F_6_N_3_O_2_ (MH)^+^ 478.3; observed 478.0.

**<u>RTI-76:</u>** A mixture containing **10** (465 mg, 0.974 mmol), stannous chloride dihydrate (770 mg, 3.41 mmol) and concentrated HCl (1.5 mL, 18.0 mmol) in ethanol (10 mL) was heated to 50° C for 3.5 hours. Upon cooling to room temperature, the solvent was concentrated and the residue was diluted with ethyl acetate and 2 N NaOH. The aqueous layer was extracted with ethyl acetate and the combined organics were washed with water and brine, dried (Na_2_SO_4_), filtered and concentrated to yield 537 mg (> 100%) of a yellow gel (**RTI-76**), which required no further purification. ^1^H-NMR (CDCl_3_) δ 7.69 (dd, 4 H, J = 6 Hz, 9 Hz), 7.52 (dd, 4 H, J = 6 Hz, 9 Hz), 7.44 (s, 1 H), 7.07 (dd, 2 H, J = 9 Hz), 6.71 (d, 2 H, J = 9 Hz), 3.91 (br s, 2 H).

**<u>RTI-77:</u>** A mixture containing **RTI-76** (0.974 mmol), Boc-glycine (260 mg, 1.46 mmol), diisopropylethylamine (0.6 mL, 3.44 mmol) and Propylphosphonic anhydride solution, 50 wt. % in ethyl acetate (1.8 mL, 3.02 mmol) in anhydrous tetrahydrofuran (35 mL) was sealed tightly and stirred at room temperature for 68 hours. The solvent was concentrated to 20% volume and the residue was partitioned between ethyl acetate and saturated aqueous NaHCO_3_. The organic layer was washed with brine, dried (Na_2_SO_4_), filtered and concentrated. The crude material was adsorbed onto silica gel and purified via ISCO using 1-50% ethyl acetate from hexanes to yield 542 mg (92%) of an off-white solid (**RTI-77**). ^1^H-NMR (CDCl_3_) δ 8.52 (br s, 1 H), 7.71-7.63 (m, 7 H), 7.53-7.47 (m, 5 H), 7.28-7.22 (m, 1 H), 5.24 (br s, 1 H), 3.94 (d, 2 H, J = 6 Hz), 1.49 (s, 9 H).

**<u>RTI-79:</u>** A solution of **RTI-77** (540 mg, 0.893 mmol) in dichloromethane (26 mL) was cooled to 0° C and treated with trifluoroacetic acid (1.4 mL, 18.8 mmol). The reaction warmed to room temperature and stirred for 18 hours. Upon completion, the mixture was concentrated and the residue was partitioned between ethyl acetate and 2 N aqueous NaOH. The aqueous layer was extracted with ethyl acetate and the combined organic layers were washed with brine, dried (Na_2_SO_4_), filtered and concentrated to yield 405 mg (90%) of a yellow solid (**RTI-79**), which required no further purification. ^1^H-NMR (CDCl_3_) δ 9.68 (br s, 1 H), 7.72 (d, 2 H, J = 9 Hz), 7.64 (dd, 4 H, J = 9 Hz), 7.55-7.49 (m, 5 H), 7.25 (d, 2 H, J = 6 Hz), 3.52 (s, 2 H).

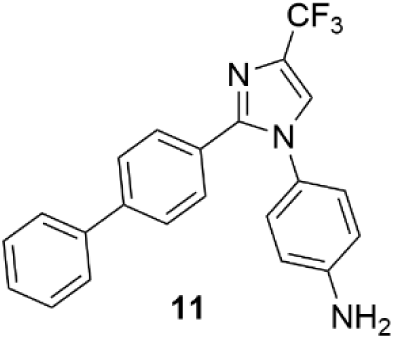

**<u>11</u>:** Using (1,1’-biphenyl)-4-carboxaldehyde as the starting material through the procedures to prepare **RTI-76**, **11** was isolated as a yellow solid (3.81 g). ^1^H-NMR (CDCl_3_) δ 7.61 (dd, 2 H, J = 6 Hz), 7.50 (dd, 4 H, J = 3 Hz, 6 Hz), 7.45-7.32 (m, 4 H), 7.05 (d, 2 H, J = 6 Hz), 6.71 (d, 2 H, J = 6 Hz), 3.89 (br s, 2 H).

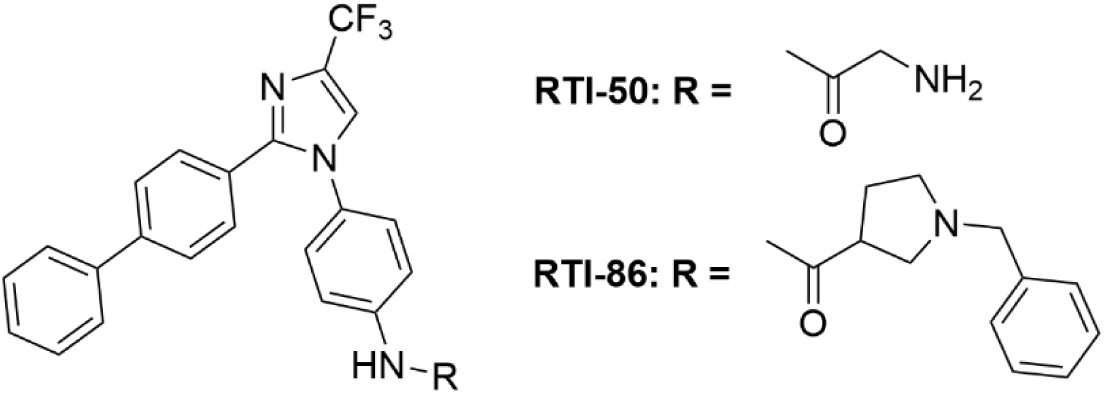

**<u>RTI-86</u>:** A mixture containing **11** (400 mg, 1.05 mmol), 1-(Phenylmethyl)-3-pyrrolidinecarboxylic acid (325 mg, 1.58 mmol), diisopropylethylamine (0.65 mL, 3.73 mmol) and Propylphosphonic anhydride solution, 50 wt. % in ethyl acetate (1.9 mL, 3.19 mmol) in anhydrous tetrahydrofuran (38 mL) was sealed tightly and stirred at room temperature for 18 hours. The solvent was concentrated to 20% volume and the residue was partitioned between ethyl acetate and saturated aqueous NaHCO_3_. The organic layer was washed with brine, dried (Na_2_SO_4_), filtered and concentrated. The crude material was adsorbed onto silica gel and purified via ISCO using 0-10% methanol from dichloromethane to yield 513 mg (86%) of an off-white solid (**RTI-86**). ^1^H-NMR (CDCl_3_) δ 9.77 (s, 1 H), 7.57 (dd, 4 H, J = 6 Hz, 9 Hz), 7.48 (dd, 6 H, J = 6 Hz), 7.42 (d, 1 H, J = 6 Hz), 7.37-7.32 (m, 5 H), 7.29-7.20 (m, 4 H), 3.73 (dd, 2 H, J = 6 Hz, 12 Hz), 3.16 (dd, 2 H, J = 6 Hz, 9 Hz), 2.95 (dd, 1 H, J = 6 Hz, 9 Hz), 2.42-2.33 (m, 3 H), 2.12-2.05 (m, 1 H). LC-MS, calculated for C_34_H_29_F_3_N_4_O (MH)^+^ 567.6; observed 567.2. Anal. Calculated for C_34_H_29_F_3_N_4_O; C, 72.07; H, 5.15; N, 9.88. Found: C, 71.97; H, 5.25; N, 9.82.

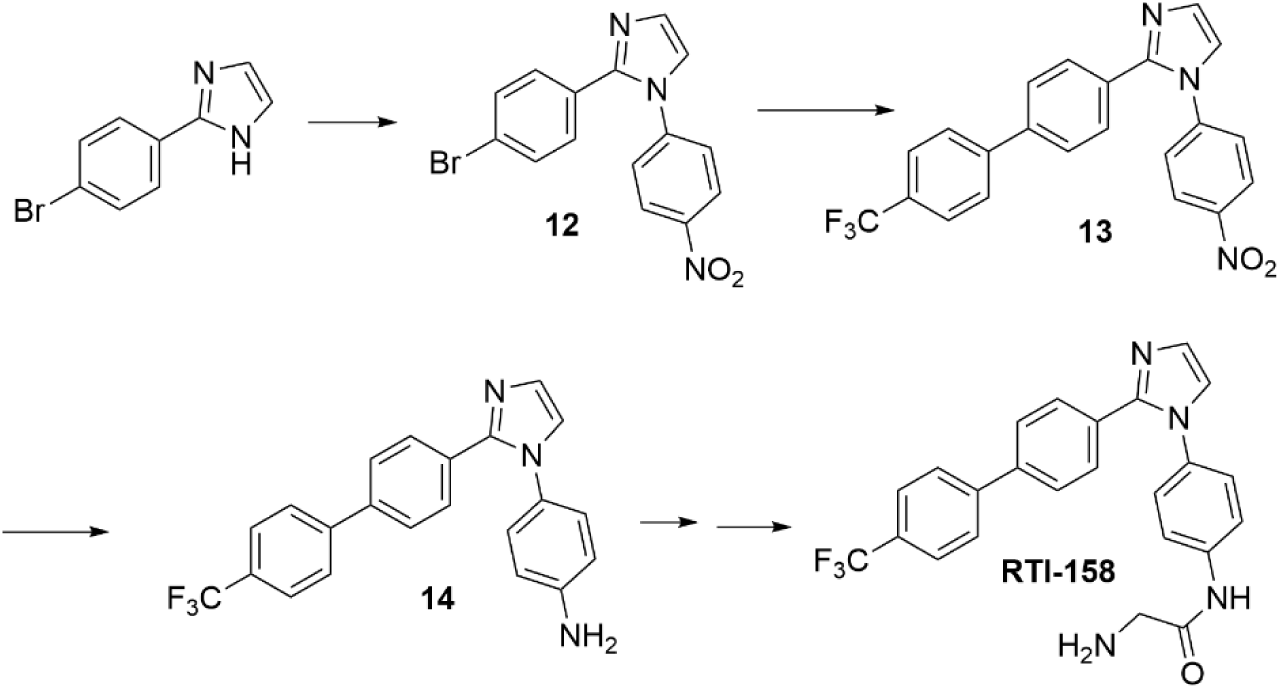

**<u>12</u>:** A mixture containing 2-(4-Bromophenyl)-1H-imidazole (500 mg, 2.25 mmol), 4-fluoronitrobenzene (950 mg, 6.74 mmol) and K_2_CO_3_ (930 mg, 6.74 mmol) in N, N-dimethylformamide (21 mL) was heated to 90° C for 21 hours. Upon cooling to room temperature, the poured into saturated aqueous LiCl solution and extracted with ethyl acetate. The combined organics were washed with water (5 x) and brine, dried (MgSO_4_), filtered and concentrated. The crude material was adsorbed onto silica gel and purified via ISCO using 0-5% methanol from dichloromethane to yield 700 mg (90%) of an off-white solid (**12**).

**<u>13</u>:** A mixture containing **12** (500 mg, 1.46 mmol), 4-trifluoromethylphenylboronic acid (330 mg, 1.75 mmol), palladium tetrakis(triphenylphosphine) (170 mg, 0.146 mmol) and 1.0 M aqueous Na_2_CO_3_ (4.37 mL, 4.37 mmol) in ethylene glycol dimethyl ether (15 mL) was heated to 100° C for 21 hours. Upon cooling to room temperature, the reaction was diluted with water and poured extracted with ethyl acetate. The combined organics were washed with brine, dried (MgSO_4_), filtered and concentrated. The crude material was adsorbed onto silica gel and purified via ISCO using 10-75% ethyl acetate from hexanes to yield 500 mg (84%) of an off-white solid (**13**).

**<u>14:</u>** A mixture containing **13** (500 mg, 1.22 mmol), stannous chloride dihydrate (1.85 g, 9.78 mmol) and 1.0 M HCl (24 mL, 24 mmol) in ethanol (40 mL) was heated to 50° C for 3.5 hours. Upon cooling to room temperature, the solvent was concentrated and the residue was diluted with ethyl acetate and 2 N NaOH. The aqueous layer was extracted with ethyl acetate and the combined organics were washed with water and brine, dried (Na_2_SO_4_), filtered and concentrated. The crude material was adsorbed onto silica gel and purified via ISCO using 0-5% methanol from dichloromethane to yield 470 mg (>100%) of an off-white solid (**14**).

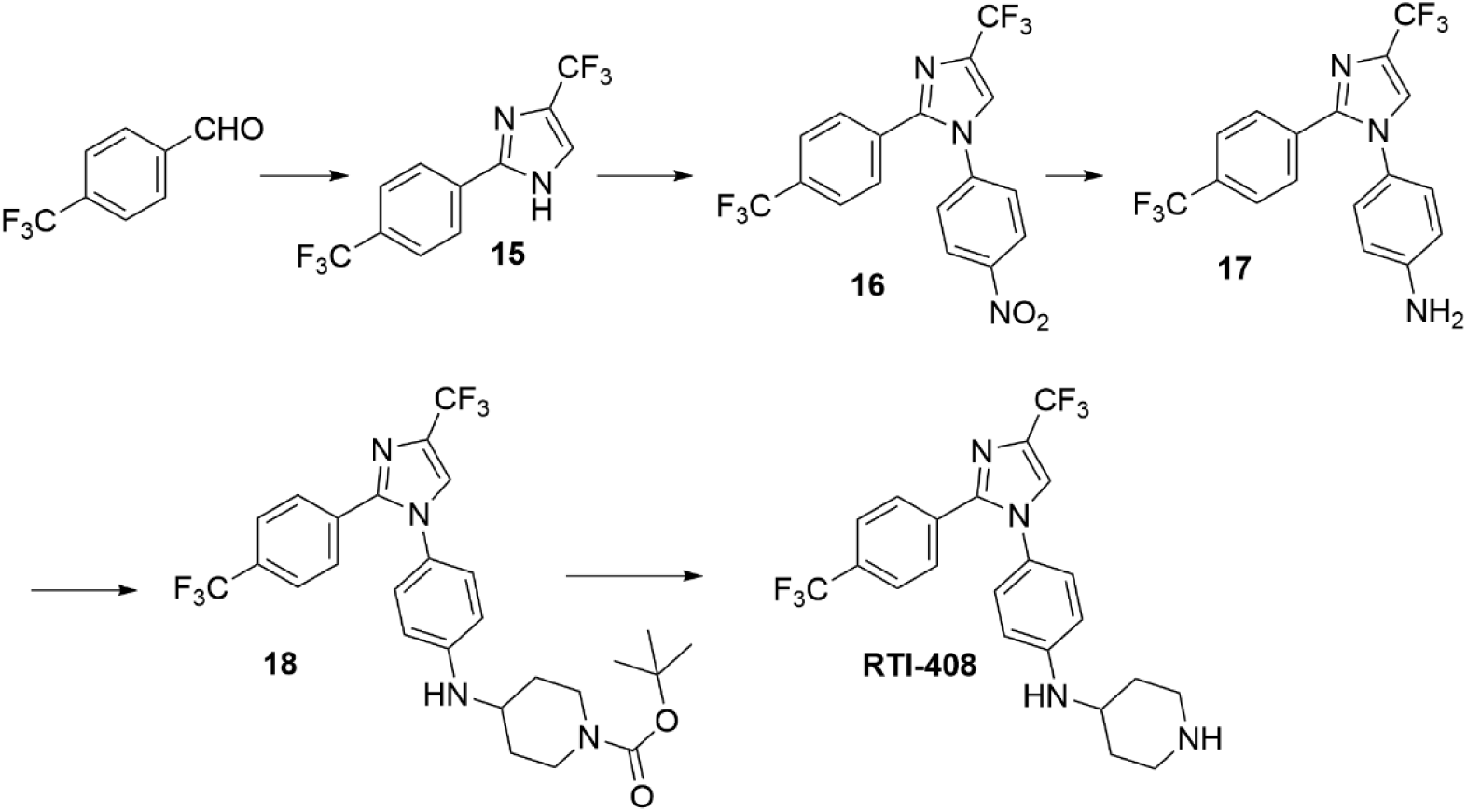

**<u>15</u>:** A mixture containing 1,1-Dibromo-3,3,3-trifluoroacetone (4.64 g, 17.2 mmol) and sodium acetate (1.88 g, 23.0 mmol) in water (42 mL) was heated to 95° C for 1 hour, then cooled to room temperature. 4’-Trifluoromethylbenzaldehyde (2.0 g, 11.4 mmol) in methanol (102 mL) was cooled to 0° C and treated with dropwise addition of ammonium hydroxide (50 mL). This mixture stirred for 10 minutes at 0° C and then the first mixture was added dropwise; the combined reaction slowly warmed to room temperature and stirred for 18 hours. The resulting solids were filtered, washed with water and dried and the filtrate was concentrated. The dried solids and filtrate residue were partitioned between dichloromethane and water. The organic layer was washed with brine, dried (Na_2_SO_4_) and concentrated. The crude material was adsorbed onto silica gel and purified via ISCO using 0-30% ethyl acetate from hexanes to yield 1.0 g (33%) of a yellow solid (**15**).

**<u>16</u>:** A mixture containing **15** (1.0 g, 3.57 mmol), 4-fluoronitrobenzene (1.52 g, 10.7 mmol) and K_2_CO_3_ (1.48 g, 10.7 mmol) in N, N-dimethylformamide (40 mL) was heated to 90° C for 21 hours. Upon cooling to room temperature, the poured into saturated aqueous LiCl solution and extracted with ethyl acetate. The combined organics were washed with water (5 x) and brine, dried (MgSO_4_), filtered and concentrated. The crude material was adsorbed onto silica gel and purified via ISCO using 0-5% methanol from dichloromethane to yield 1.0 g (70%) of an off-white solid (**16**).

**<u>17:</u>** A mixture containing **16** (1.0 g, 2.49 mmol), stannous chloride dihydrate (3.77 g, 19.9 mmol) and concentrated HCl (4.15 mL, 49.9 mmol) in ethanol (40 mL) was heated to 50° C for 3.5 hours. Upon cooling to room temperature, the solvent was concentrated and the residue was diluted with ethyl acetate and 2 N NaOH. The aqueous layer was extracted with ethyl acetate and the combined organics were washed with water and brine, dried (Na_2_SO_4_), filtered and concentrated. The crude material was adsorbed onto silica gel and purified via ISCO using 0-5% methanol from dichloromethane to yield 275 mg (70%) of an off-white solid (**17**).

### Triazole Scaffold

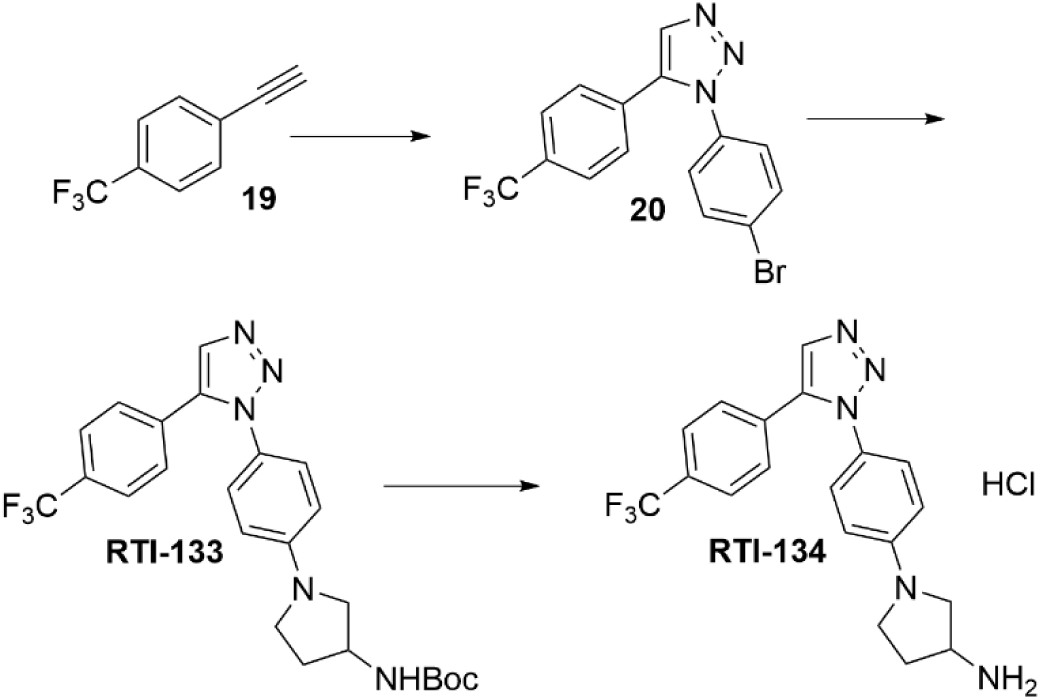

**<u>20:</u>** A mixture containing **19** (198 mg, 1.0 mmol) and 4-Trifluoromethylstyrene (179 mg, 1.05 mmol) in dimethylsulfoxide (3.3 mL) was treated with tetramethylammonium hydroxide (25% solution – 9.1 mg/37uL of water) and stirred at room temperature for 20 hours. The reaction was diluted with water (15 mL) and extracted with ethyl acetate. The organic layer was washed with water and brine, dried (Na_2_SO_4_), filtered and concentrated. The crude material was adsorbed onto silica gel and purified via ISCO using 5-30% ethyl acetate from hexanes to yield 350 mg (95%) of a yellow-brown solid (**20**).

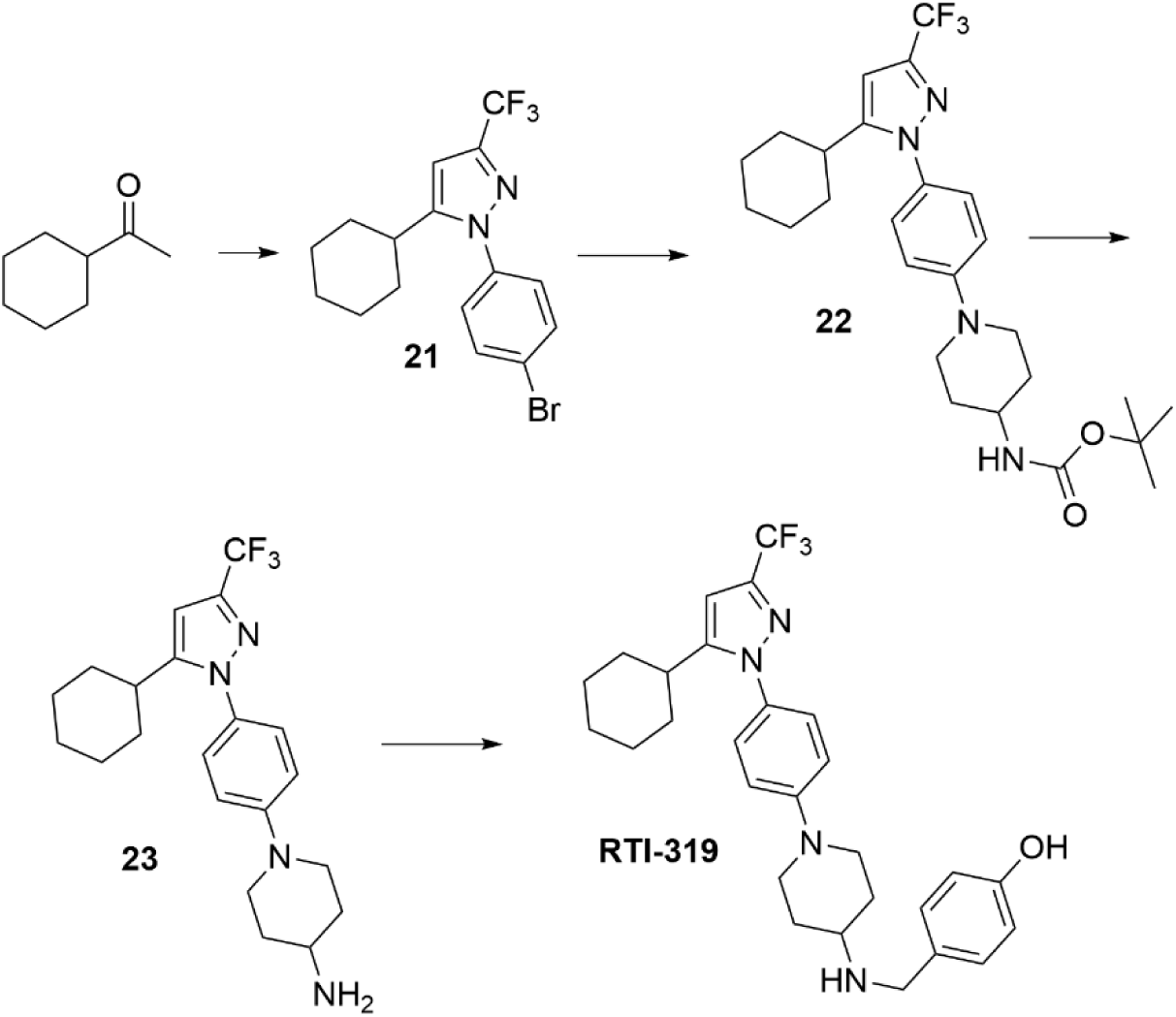

**<u>21</u>:** Into an oven dried flask was introduced sodium hydride (60% wt./mineral oil, 9.52 g, 237.6 mmol) and stirred in anhydrous THF (140 mL) for 5 minutes at room temperature. Ethyl trifluoroacetate (18.9 mL, 158.8 mmol) was added dropwise and this mixture stirred at room temperature for 10 minutes. A solution of cyclohexyl methyl ketone (10 g, 79.2 mmol) in anhydrous THF (40 mL) was added dropwise and the reaction mixture was refluxed for 3 hours. The reaction was concentrated and the residue was partitioned between ethyl acetate and water. The aqueous layer was extracted with ethyl acetate and the combined organics were washed with brine, dried (Na_2_SO_4_) and concentrated to yield a quantitative yield of a yellow oil, which was used without further purification. ^1^H-NMR (CD_3_OD) δ 5.24 (s, 1 H), 2.94-2.84 (m, 1 H), 1.82-1.66 (m, 4 H), 1.40-1.19 (m, 6 H).

A mixture containing the previous product (79.2 mmol) and 4-bromohydrazine hydrochloride (24 g, 103 mmol) in ethanol (1 L) was refluxed for 18 hours. The solvent was concentrated and the residue was partitioned between ethyl acetate and saturated aqueous NaHCO_3_. The aqueous layer was extracted with ethyl acetate and the combined organics were washed with brine, dried (MgSO_4_), filtered and concentrated. The crude material was purified over silica gel via a flush column using 3-5% ethyl acetate from hexanes to remove baseline impurities from the isomeric product mixture. Upon sitting at room temperature, a yellow solid (undesired isomer) precipitated from the mixture. The liquid portion of the mixture (rich in **21**) was decanted and the yellow solid was triturated in hexanes to remove all of **21** (with some of the undesired isomer) and filtered. This combined filtrate and decantate was concentrated, adsorbed onto silica gel and purified via ISCO using 2-15% dichloromethane from hexanes to yield 4.58 g (15%) of an orange oil (**21**). ^1^H-NMR (CDCl_3_) δ 7.59 (d, 2 H, J = 9 Hz), 7.37 (d, 2 H, J = 9 Hz), 6.62 (s, 1 H), 2.75-2.66 (m, 1 H), 2.09-1.99 (m, 2 H), 1.90-1.70 (m, 2 H), 1.53-1.21 (m, 6 H). Anal. Calculated for C_16_H_16_BrF_3_N_2_; C, 51.48; H, 4.32; N, 7.50. Found: C, 51.73; H, 4.51; N, 7.51.

**<u>22:</u>** The following were combined in a heavy-duty glass reactor: **21** (1.1 g, 2.95 mmol), 4-(tert-butoxycarbonylamino)piperidine (1.18 g, 5.89 mmol), BINAP (551 mg, 0.884 mmol), Pd_2_(dba)_3_ (540 mg, 0.589 mmol) and Cs_2_CO_3_ (1.92 g, 5.89 mmol) in anhydrous toluene (40 mL) and nitrogen gas was bubbled into the mixture for two minutes. The reactor was then sealed with a Teflon cap and heated to 110° C for 15 hours. Upon cooling, the mixture was filtered through Celite and the filter pad was rinsed with ethyl acetate. The filtrate was washed with water and brine, dried (Na_2_SO_4_), filtered and concentrated to obtain 3.51 g of a rust-colored gel. The crude material was purified over silica gel using 0-100% ethyl acetate from hexanes to yield 426 mg (29%) of a yellow solid (**22**). ^1^H-NMR (CDCl_3_) δ 7.31 (d, 2 H, J = 9 Hz), 6.93 (d, 2 H, J = 9 Hz), 6.55 (s, 1 H), 4.48 (br s, 1 H), 3.73-3.64 (m, 3 H), 2.91 (dd, 2 H, J = 3 Hz, 9 Hz), 2.74-2.67 (m, 1 H), 2.07-2.00 (m, 4 H), 1.84-1.67 (m, 7 H), 1.62-1.25 (m, 7 H), 1.46 (s, 9 H).

**<u>23:</u>** A solution of **22** (426 mg, 0.865 mmol) in dichloromethane (17 mL) was cooled to 0° C and treated with trifluoroacetic acid (0.65 mL, 8.75 mmol). The reaction warmed to room temperature and stirred for 15 hours. Upon completion, the mixture was concentrated and the residue was partitioned between ethyl acetate and 2 N aqueous NaOH. The aqueous layer was extracted with ethyl acetate and the combined organic layers were washed with brine, dried (Na_2_SO_4_), filtered and concentrated to yield 281 mg of **23** (>100%) of a yellow gel, which required no further purification. ^1^H-NMR (CDCl_3_) δ 7.30 (d, 2 H, J = 9 Hz), 6.94 (d, 2 H, J = 9 Hz), 6.55 (s, 1 H), 3.75-3.69 (m, 2 H), 2.89-2.80 (m, 2 H), 2.73-2.66 (m, 1 H), 2.10-1.71 (m, 7 H), 1.53-1.27 (m, 8 H). LC-MS, calculated for C_21_H_27_F_3_N_4_ (MH)^+^ 393.4; observed 393.0.

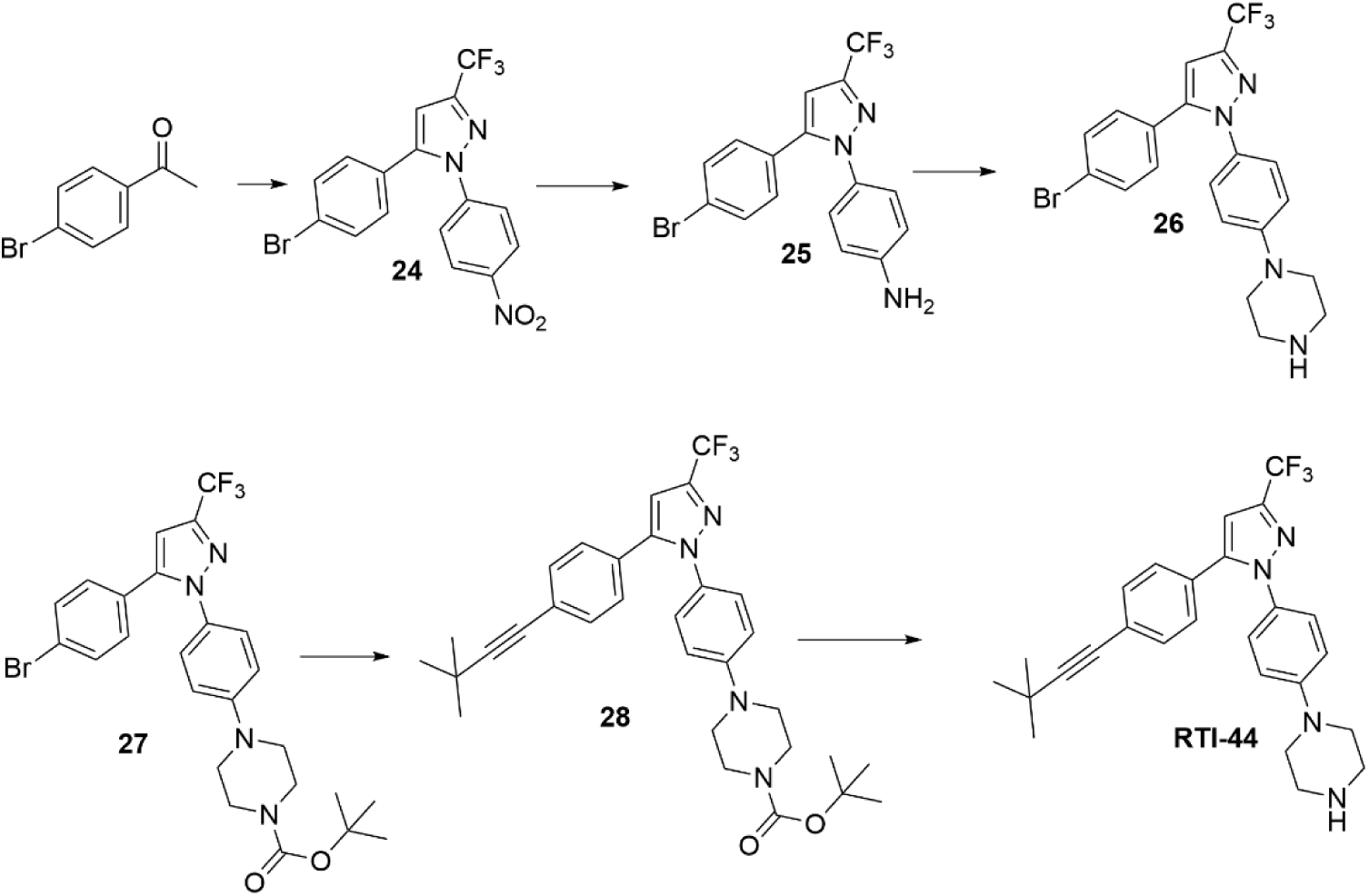

**<u>24</u>:** To a suspension of sodium hydride (60% wt./mineral oil, 1.69 g, 70.3 mmol) and stirred in anhydrous THF (50 mL) in a 3-neck flask fitted with a reflux condenser, ethyl trifluoroacetate (7.14 g, 50.2 mmol), in 6 mL of anhydrous THF, was added dropwise; this mixture stirred at room temperature for 10 minutes. A solution of p-Bromoacetophenone (5 g, 25.1 mmol) in anhydrous THF (25 mL) was added dropwise, over 30 minutes, and the reaction mixture was refluxed for 3 hours. The reaction was quenched acetic acid/water (5 mL, 1/1 ratio) and concentrated. The residue was partitioned between ethyl acetate and acetic acid/water and the combined organics were washed with brine, dried (Na_2_SO_4_) and concentrated to generate 6.5 g (88%) of a yellow solid, which was used without further purification.

A mixture containing the previous product (0.733 g, 2.5 mmol) and 4-nitrohydrazine hydrochloride (0.57 g, 3.0 mmol) in ethanol (25 mL) was refluxed for 2 hours. Another 47 mg (0.25 mmol) of 4-nitrohydrazine hydrochloride was added and the reaction refluxed for 15 hours. The solvent was concentrated and the residue was partitioned between ethyl acetate and water. The aqueous layer was extracted with ethyl acetate and the combined organics were washed with brine, dried (MgSO_4_), filtered and concentrated. The crude material was adsorbed onto silica gel and purified via ISCO using 0.5-10% ethyl acetate from hexanes to yield 0.69 g (67%) of a white solid (**24**).

**<u>25:</u>** A mixture containing **24** (82.4 mg, 0.20 mmol), stannous chloride dihydrate (133 mg, 0.70 mmol) and concentrated HCl (0.3 mL, 3.6 mmol) in ethanol (1.5 mL) was heated to 50° C for 2.5 hours. Upon cooling to room temperature, the solvent was concentrated and the residue was diluted with ethyl acetate and 2 N NaOH. The aqueous layer was extracted with ethyl acetate and the combined organics were washed with water and brine, dried (Na_2_SO_4_), filtered and concentrated to yield 75 mg (98%) of an off-white solid (**25**), which required no further purification.

**<u>26:</u>** A mixture containing **25** (38.2 mg, 0.10 mmol), Bis(2-chloroethyl)amine hydrochloride (35.6 mg, 0.20 mmol) and K_2_CO_3_ (27.6 mg, 0.20 mmol) in 1-methoxyethanol (1.0 mL) was heated to 175° C for 2 hours via microwaves. The reaction was heated for another 1 hour to enhance completion. Upon cooling to room temperature, the reaction was diluted with 2 N NaOH. The mixture was extracted with ethyl acetate and the combined organics were washed with water and brine, dried (Na_2_SO_4_), filtered and concentrated. The crude material was adsorbed onto silica gel and purified via ISCO using 0-15% (1% NH_4_OH/methanol) from dichloromethane to yield 27 mg (60%) of a colorless oil (**26**).

**<u>27:</u>** A mixture containing **26** (44 mg, 0.097 mmol), di-tert-butyl dicarbonate (23.4 mg, 0.107 mmol) in ethanol (1.0 mL) was heated to 30° C for 30 minutes. The reaction was concentrated and diluted with water. The mixture was extracted with ethyl acetate and the combined organics were washed with water and brine, dried (Na_2_SO_4_), filtered and concentrated. The crude material was adsorbed onto silica gel and purified via ISCO using 0-20% ethyl acetate from hexanes to yield 29 mg (54%) of a colorless oil (**27**).

**<u>28:</u>** The following were combined in an I-CHEM vial with a screw cap septum: **27** (43.8 mg, 0.08 mmol), 3,3-Dimethyl-1-butyne (16 mg, 0.16 mmol), Palladium tetrakis(triphenylphosphine) (9.2 mg, 0.008 mmol), CuI (3.0 mg, 0.016 mmol) in diisopropylamine (2 mL). The vial was repeatedly evacuated and backfilled with nitrogen. The reaction was heated to 70° C for 15 hours. Upon cooling, the mixture was filtered through Celite and the filter pad was rinsed with ethyl acetate. The filtrate was concentrated and the crude material was purified over silica gel using 0-18% ethyl acetate from hexanes to yield 43 mg (94%) of a colorless oil (**28**).

**<u>RTI-44:</u>** A solution of **28** (34 mg, 0.061 mmol) in dichloromethane (2 mL) was cooled to 0° C and treated with trifluoroacetic acid (0.2 mL, 2.69 mmol). The reaction warmed to room temperature and stirred for 15 hours. Upon completion, the mixture was concentrated and the residue was partitioned between ethyl acetate and 2 N aqueous NaOH. The aqueous layer was extracted with ethyl acetate and the combined organic layers were washed with brine, dried (Na_2_SO_4_), filtered and concentrated to yield 281 mg of **RTI-44** (>100%) of a yellow gel, which required no further purification. ^1^H-NMR (CD_3_OD) δ 7.67-7.61 (m, 2 H), 7.58-7.52 (m, 2 H), 7.30-7.17 (m, 4 H), 3.49-3.43 (m, 4 H), 3.37-3.32 (m, 4 H), 1.27 (s, 9 H).

## Notes

### Competing Interest Statement

The authors have declared no competing interest.

## REFERENCES

1. Seaman J, Mercer AJ, Sondorp HE, Herwaldt BL. Epidemic visceral leishmaniasis in southern Sudan: treatment of severely debilitated patients under wartime conditions and with limited resources. Ann Intern Med. 1996;124(7):664–72. Epub 1996/04/01. PubMed PMID: 8607595.

2. Paila YD, Saha B, Chattopadhyay A. Amphotericin B inhibits entry of Leishmania donovani into primary macrophages. Biochem Biophys Res Commun. 2010;399(3):429–33. Epub 2010/08/04. doi: S0006-291X(10)01419-1 [pii] 10.1016/j.bbrc.2010.07.099. PubMed PMID: 20678487.

3. Purkait B, Kumar A, Nandi N, Sardar AH, Das S, Kumar S, et al. Mechanism of amphotericin B resistance in clinical isolates of Leishmania donovani. Antimicrob Agents Chemother. 2012;56(2):1031–41. Epub 2011/11/30. doi: AAC.00030-11 [pii] 10.1128/AAC.00030-11. PubMed PMID: 22123699; PubMed Central PMCID: PMC3264217.

4. Garcia-Hernandez R, Manzano JI, Castanys S, Gamarro F. Leishmania donovani Develops Resistance to Drug Combinations. PLoS Negl Trop Dis. 2012;6(12):e1974. Epub 2013/01/04. doi: 10.1371/journal.pntd.0001974 PNTD-D-12-00833 [pii]. PubMed PMID: 23285310; PubMed Central PMCID: PMC3527373.

5. Zahid MSH, Varma DM, Johnson MM, Landavazo A, Bachelder EM, Blough BE, et al. Overcoming reduced antibiotic susceptibility in intracellular Salmonella enterica serovar Typhimurium using AR-12. FEMS Microbiol Lett. 2021;368(11). Epub 2021/06/06. doi: 10.1093/femsle/fnab062. PubMed PMID: 34089315; PubMed Central PMCID: PMCPMC8433491.

6. Franco LH, Fleuri AKA, Pellison NC, Quirino GFS, Horta CV, de Carvalho RVH, et al. Autophagy downstream of endosomal Toll-like receptor signaling in macrophages is a key mechanism for resistance to Leishmania major infection. J Biol Chem. 2017;292(32):13087–96. Epub 2017/06/14. doi: 10.1074/jbc.M117.780981. PubMed PMID: 28607148; PubMed Central PMCID: PMCPMC5555173.

7. Casanova JE. Bacterial Autophagy: Offense and Defense at the Host-Pathogen Interface. Cell Mol Gastroenterol Hepatol. 2017;4(2):237–43. Epub 2017/07/01. doi: 10.1016/j.jcmgh.2017.05.002. PubMed PMID: 28660242; PubMed Central PMCID: PMCPMC5480303.

8. Galluzzi L, Diotallevi A, De Santi M, Ceccarelli M, Vitale F, Brandi G, et al. Leishmania infantum Induces Mild Unfolded Protein Response in Infected Macrophages. PLoS One. 2016;11(12):e0168339. doi: 10.1371/journal.pone.0168339. PubMed PMID: 27978534; PubMed Central PMCID: PMCPMC5158320.

9. Collier MA, Peine KJ, Gautam S, Oghumu S, Varikuti S, Borteh H, et al. Host-mediated Leishmania donovani treatment using AR-12 encapsulated in acetalated dextran microparticles. International Journal of Pharmaceutics. 2016;499(1-2):186–94. doi: 10.1016/j.ijpharm.2016.01.004. PubMed PMID: WOS:000370048300019.

10. Zahid MSH, Johnson MM, Tokarski RJ, 2nd, Satoskar AR, Fuchs JR, Bachelder EM, et al. Evaluation of synergy between host and pathogen-directed therapies against intracellular Leishmania donovani. Int J Parasitol Drugs Drug Resist. 2019;10:125–32. Epub 2019/09/08. doi: 10.1016/j.ijpddr.2019.08.004. PubMed PMID: 31493763; PubMed Central PMCID: PMCPMC6731340.

11. Saha AK, Mukherjee T, Bhaduri A. Mechanism of action of amphotericin B on Leishmania donovani promastigotes. Mol Biochem Parasitol. 1986;19(3):195–200. Epub 1986/06/01. doi: 10.1016/0166-6851(86)90001-0. PubMed PMID: 3736592.

12. Ferraboschi P, Ciceri S, Grisenti P. Applications of Lysozyme, an Innate Immune Defense Factor, as an Alternative Antibiotic. Antibiotics (Basel). 2021;10(12). Epub 2021/12/25. doi: 10.3390/antibiotics10121534. PubMed PMID: 34943746; PubMed Central PMCID: PMCPMC8698798.

13. Valigurová A, Koláøová I. Unrevealing the Mystery of Latent Leishmaniasis: What Cells Can Host Leishmania? Pathogens. 2023;12(2). Epub 2023/02/26. doi: 10.3390/pathogens12020246. PubMed PMID: 36839518; PubMed Central PMCID: PMCPMC9967396.

14. Kumar P, Kumar R, Pandey H, Sundar S, Pai K. Studies on the arginase, 5’-nucleotidase and lysozyme activity by monocytes from visceral leishmaniasis patients. J Parasit Dis. 2012;36(1):19–25. Epub 2013/04/02. doi: 10.1007/s12639-011-0066-z. PubMed PMID: 23542635; PubMed Central PMCID: PMCPMC3284622.

15. McCracken NA, Liu H, Runnebohm AM, Wijeratne HRS, Wijeratne AB, Staschke KA, et al. Obtaining Functional Proteomics Insights From Thermal Proteome Profiling Through Optimized Melt Shift Calculation and Statistical Analysis With InflectSSP. Molecular & Cellular Proteomics. 2023;22(9):100630. doi: 10.1016/j.mcpro.2023.100630.

16. Mateus A, Kurzawa N, Becher I, Sridharan S, Helm D, Stein F, et al. Thermal proteome profiling for interrogating protein interactions. Mol Syst Biol. 2020;16(3):e9232. Epub 2020/03/07. doi: 10.15252/msb.20199232. PubMed PMID: 32133759; PubMed Central PMCID: PMCPMC7057112.

17. Lezama-Davila CM, Isaac-Marquez AP, Kapadia G, Owens K, Oghumu S, Beverley S, et al. Leishmanicidal activity of two naphthoquinones against Leishmania donovani. Biol Pharm Bull. 2012;35(10):1761–4. Epub 2012/10/06. doi: 10.1248/bpb.b12-00419. PubMed PMID: 23037165; PubMed Central PMCID: PMCPMC3693454.

18. Weischenfeldt J, Porse B. Bone Marrow-Derived Macrophages (BMM): Isolation and Applications. CSH Protoc. 2008;2008:pdb.prot5080. Epub 2008/01/01. doi: 10.1101/pdb.prot5080. PubMed PMID: 21356739.

19. Zahid MSH, Johnson MM, Tokarski RJ, Satoskar AR, Fuchs JR, Bachelder EM, et al. Evaluation of synergy between host and pathogen-directed therapies against intracellular Leishmania donovani. International Journal for Parasitology: Drugs and Drug Resistance. 2019;10:125–32. doi: 10.1016/j.ijpddr.2019.08.004.

20. Gaetani M, Sabatier P, Saei AA, Beusch CM, Yang Z, Lundström SL, et al. Proteome Integral Solubility Alteration: A High-Throughput Proteomics Assay for Target Deconvolution. J Proteome Res. 2019;18(11):4027–37. Epub 2019/09/24. doi: 10.1021/acs.jproteome.9b00500. PubMed PMID: 31545609.

21. Kubota K, Funabashi M, Ogura Y. Target deconvolution from phenotype-based drug discovery by using chemical proteomics approaches. Biochim Biophys Acta Proteins Proteom. 2019;1867(1):22–7. Epub 2018/11/06. doi: 10.1016/j.bbapap.2018.08.002. PubMed PMID: 30392561.

22. Cabrera A, Wiebelhaus N, Quan B, Ma R, Meng H, Fitzgerald MC. Comparative Analysis of Mass-Spectrometry-Based Proteomic Methods for Protein Target Discovery Using a One-Pot Approach. J Am Soc Mass Spectrom. 2020;31(2):217–26. Epub 2020/02/08. doi: 10.1021/jasms.9b00041. PubMed PMID: 32031398; PubMed Central PMCID: PMCPMC7441748.

23. Clausen BE, Burkhardt C, Reith W, Renkawitz R, Förster I. Conditional gene targeting in macrophages and granulocytes using LysMcre mice. Transgenic Res. 1999;8(4):265–77. Epub 2000/01/06. doi: 10.1023/a:1008942828960. PubMed PMID: 10621974.

